# A DNA barcode reference library of the French Polynesian shore fishes

**DOI:** 10.1101/595793

**Authors:** Erwan Delrieu-Trottin, Jeffrey T. Williams, Diane Pitassy, Amy Driskell, Nicolas Hubert, Jérémie Viviani, Thomas H. Cribb, Benoit Espiau, René Galzin, Michel Kulbicki, Thierry Lison de Loma, Christopher Meyer, Johann Mourier, Gérard Mou-Tham, Valeriano Parravicini, Patrick Plantard, Pierre Sasal, Gilles Siu, Nathalie Tolou, Michel Veuille, Lee Weigt, Serge Planes

## Abstract

The emergence of DNA barcoding and metabarcoding opened new ways to study biological diversity, however, the completion of DNA barcode libraries is fundamental for such approaches to succeed. This dataset is a DNA barcode reference library (fragment of Cytochrome Oxydase I gene) for 2,190 specimens representing at least 540 species of shore fishes collected over 10 years at 154 sites across the four volcanic archipelagos of French Polynesia; the Austral, Gambier, Marquesas and Society Islands, a 5,000,000 km^2^ area. At present, 65% of the known shore fish species of these archipelagoes possess a DNA barcode associated with preserved, photographed, tissue sampled and cataloged specimens, and extensive collection locality data. This dataset represents one of the most comprehensive DNA barcoding efforts for a vertebrate fauna to date. Considering the challenges associated with the conservation of coral reef fishes and the difficulties of accurately identifying species using morphological characters, this publicly available library is expected to be helpful for both authorities and academics in various fields.

## Background & Summary

DNA barcoding aims to identify individuals to the species level by using a short and standardized portion of a gene as a species tag^1^. This standardized procedure has revolutionized how biodiversity can be surveyed as the identification of a species then becomes independent of the level of taxonomic expertise of the collector^2^, the life stage of the species^3,4^ or the state of conservation of the specimen^5,6^. Due to its large spectrum of potential applications, DNA barcoding has been employed in a large array of scientific fields such as taxonomy^7^, biogeography, biodiversity inventories^8^ and ecology^9^; but see Hubert and Hanner for a review^10^. In the genomic era, this approach has been successfully applied to the simultaneous identification of multiple samples (*i.e*. the metabarcoding approach), extending its applications to surveys of whole ecological communities^11^, but also monitoring species diet^12,13^, identifying the presence of specific species in a region^14^, or studying changes in the community through time by sampling environmental DNA^15,16^.

By design, DNA barcoding has proved to be fast and accurate, but its accuracy is highly dependent on the completeness of DNA barcode reference libraries. These libraries turn surveys of Operational Taxonomic Units (OTUs) into species surveys through the assignment of species names to OTUs^17,18^, hence giving meaning to data for ecologists, evolutionary biologists and stakeholders. Taxonomists increasingly provide DNA barcodes of new species they are describing; but thousands of species of shore fishes still lack this diagnostic molecular marker.

In the South Pacific, an early initiative led by the CRIOBE Laboratory was successfully carried out for French Polynesian coral reef fishes at the scale of one island, Moorea (Society Island)^19^. The fish fauna of Moorea’s waters is one of the best known of the region given the historical operation of research laboratories and long term surveys^20,21^. The Moorea project revealed a high level of cryptic diversity in Moorea’s fishes^19^ and motivated the CRIOBE Laboratory to extend this biodiversity survey of shore fishes to the remaining islands of French Polynesia. French Polynesia (FP) is a 5,000,000 km^2^ region located between 7° and 27° South Latitude that constitutes a priority area for conducting a barcoding survey. This region is species rich due to its position at the junction of several biogeographic areas with varying levels of endemism. For example, the Marquesas Islands (northeastern FP) rank as the third highest region of endemism for coral reef fishes in the Indo-Pacific (13.7%^22^). The Austral Islands (southwestern FP) and Gambier Islands (southeastern FP) host numerous southern subtropical endemic species^23–25^. Finally, the Society Islands (western FP) possess the highest species richness (877 species) and the highest number of widespread species in French Polynesia^26^.

Here, we present the result of a large-scale effort to DNA barcode the shore fishes in French Polynesia. Conducted between 2008 and 2014, a total of 154 sites were inventoried across these four archipelagoes. Islands of varying ages and topographies were visited ranging from low-lying atolls to high islands surrounded by a barrier reef, or solely fringing reefs. Furthermore, inventories were conducted across different habitats at each island (*i.e.* sand bank, coral reefs, rubble, rocky, etc.). In total, 2,190 specimens were identified, preserved, photographed, tissue sampled, DNA barcoded and cataloged with extensive metadata to build a library representing at least 540 species, 232 genera and 61 families of fishes (Fig. 1). Merged with previous sampling efforts at Moorea, a total of 3,131 specimens now possess a DNA barcode representing at least 645 nominal species for a coverage of approximately 65% of the known shore fish species diversity of these four archipelagoes. These biodiversity surveys have already resulted in the publication of updated species checklists^22,26^ and in the description of 17 new species^27–34^. This comprehensive library for French Polynesia shore fishes will certainly benefit a wide community of users with different interests, ranging from basic to applied science, and including fisheries management, functional ecology, taxonomy and conservation. Furthermore, many newly detected taxa for science are revealed here, along with complete collection data and DNA barcodes, which should facilitate their formal description as new species. While shedding new light on the species diversity of the Pacific region, this publicly available library is expected to fuel the development of DNA barcode libraries in the Pacific Ocean and to provide more accurate results for the growing number of studies using DNA metabarcoding in the Indo-West Pacific.

**Figure 1.**
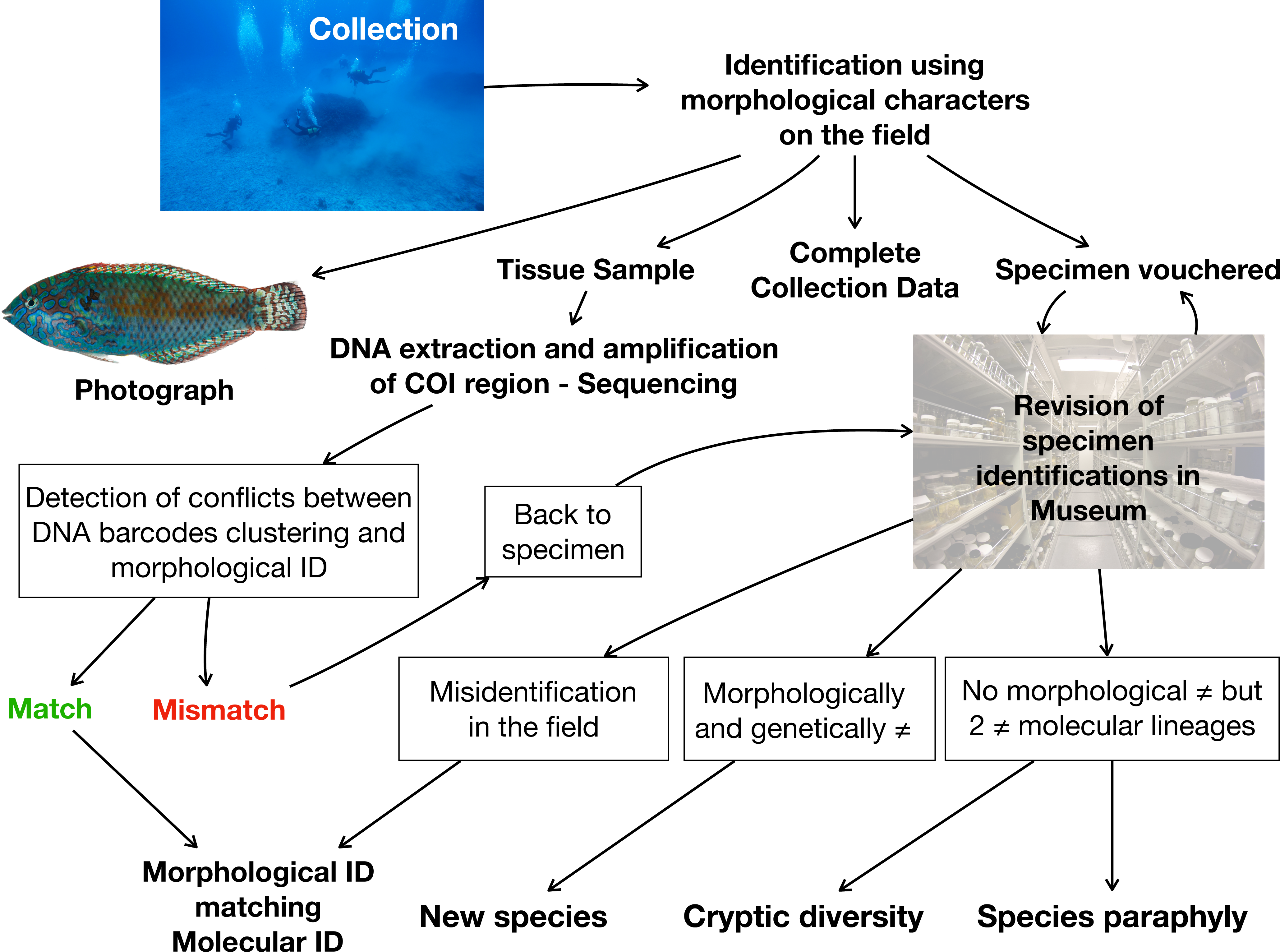
Overview of data generation. From collection of specimen to the validation of data generation.

## Methods

### Sampling strategy

We explored a diversity of habitats across the four corners of French Polynesia with shallow and deep SCUBA dives (down to 50–55 m) for a total of 154 sampled sites (Fig. 2, Table 1). A total of 2,190 specimens, representing at least 540 species, 232 genera and 61 families (Fig. 3a) have been collected across four archipelagos representing the four corners of French Polynesia (FP), through six scientific expeditions: Marquesas Islands (1) in 2008 at Mohotani and (2) in 2011 at every island of the archipelago aboard the M.V. Braveheart (Clark Bank, Motu One, Hatutaa, Eiao, Motu Iti, Nuku-Hiva, Ua-Huka, Ua-Pou, Fatu-Huku, Hiva-Oa, Tahuata, Fatu-Hiva; 52 sites), (3) in 2010 at Gambier Islands aboard the M.V. Claymore (Mangareva, Taravai, Akamaru, and all along the barrier reef; 53 sites), (4) at Austral Islands in 2013 aboard the Golden Shadow (Raivavae, Tubuai, Rurutu, Rimatara, Maria Islands; 25 sites), (5) at westernmost atolls of the Society Islands in 2014 aboard the M.V. Braveheart (Manuae and Maupiha’a; 20 sites). A sixth scientific expedition took place on Moorea’s deep reefs in 2008 (Society Islands) as a small scale scientific expedition that included the exploration and sampling of some of the deep reefs of Moorea (53 to 56 m depth; 4 sites) (Fig. 2).

**Table 1.** Overview of the dataset. Number of specimens and species collected for each scientific expedition. Sampling effort expressed in number of sampling days and number of sites.

| BOLD project | Geographical location | No. of specimen collected | No. of species collected | Sampling effort (No. of sampling days / No. of sites) |
| --- | --- | --- | --- | --- |
| AUSTR | Austral Islands | 560 | 263 | 12 / 25 |
| GAMBA | Gambier Islands | 705 | 290 | 18 / 53 |
| MARQ | Marquesas Islands | 386 | 182 | 18 / 41 |
| MOH | Marquesas Islands | 190 | 107 | 5 / 11 |
| MOOP | Society Islands | 42 | 27 | 4 / 4 |
| SCIL | Society Islands | 309 | 213 | 8 / 20 |

**Figure 2.**
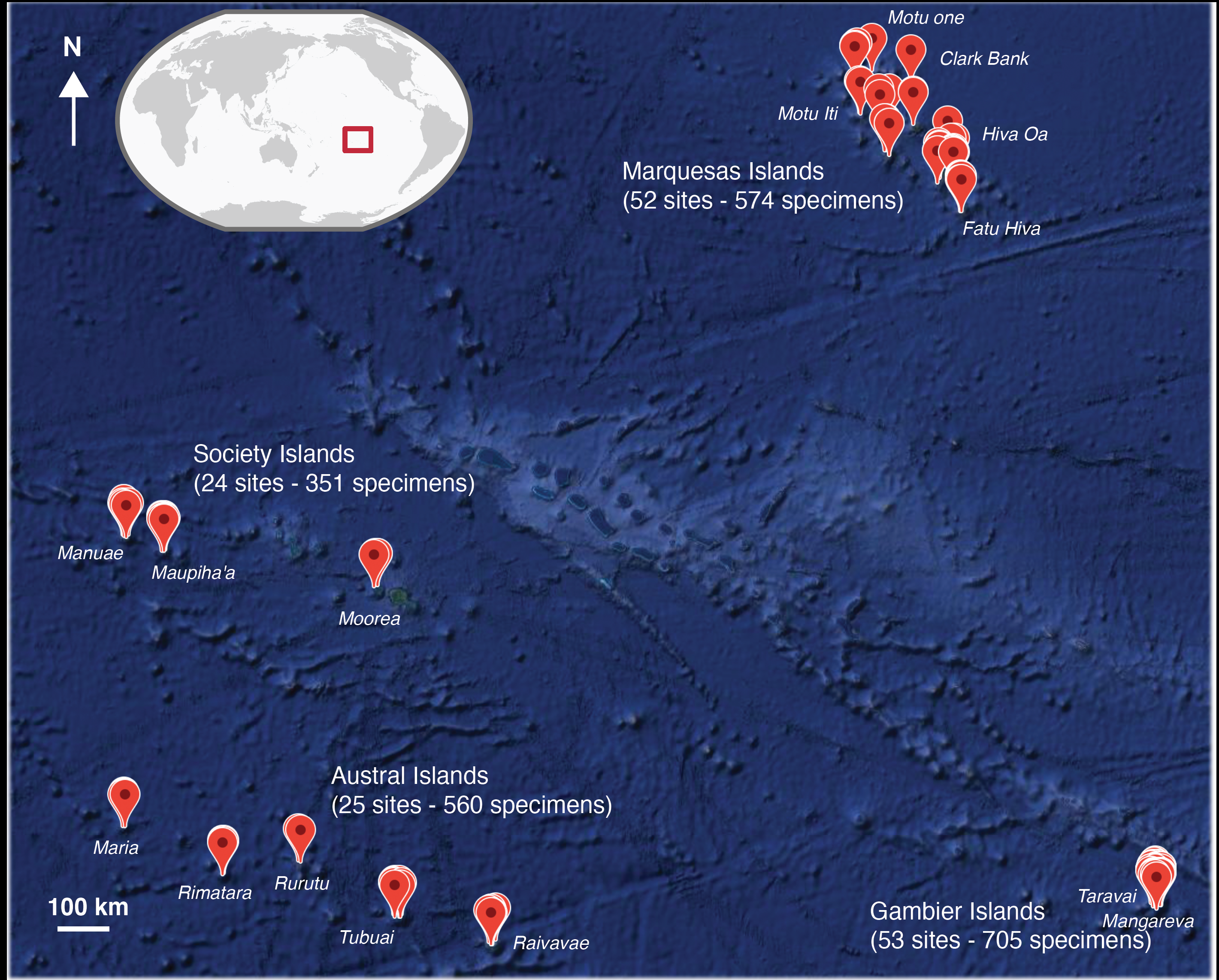
Sampling localities across French Polynesia. The number of sampling sites and the number of specimens collected are displayed for each archipelago. Several sampling localities may be represented by a single dot due to the geographic scale of French Polynesia. Map data: Google, DigitalGlobe.

**Figure 3.**
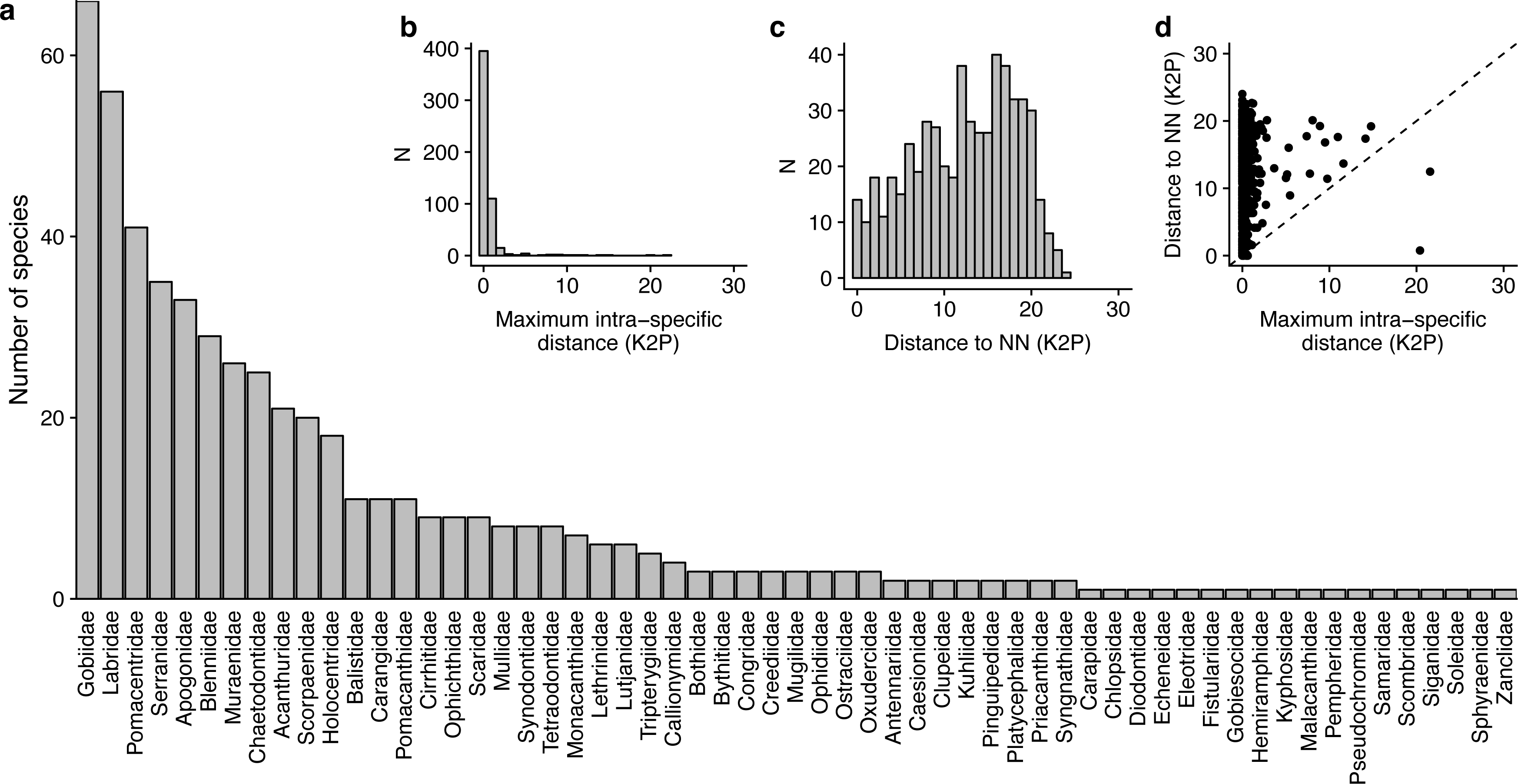
Species diversity and distribution of genetic distance across the DNA barcode library. (a) Species diversity by family for the four archipelagoes sampled; (b) Distribution of maximum intraspecific distances (K2P, percent); (c) Distribution of nearest neighbor distances (K2P, percent); (d) Relationship between maximum intraspecific and nearest neighbor distances. Points above the diagonal line indicate species with a barcode gap.

### Specimen collection

Specimens were captured using rotenone (powdered root of the Derris plant) and spear guns while SCUBA diving. These complementary sampling methods^35^ allowed us to sample both the cryptic and small fish fauna as well as the larger specimens of species not susceptible to rotenone collecting. Four individuals per species were collected on average. Fishes were sorted and identified onboard to the species level using identification keys and taxonomic references^23,36^ and representative specimens of all species collected were photographed in a fish photo tank to capture fresh color patterns, labeled and tissue sampled for genetic analyses (fin clip or muscle biopsies preserved in 96% ethanol). The photographed/sampled voucher specimens were preserved in 10% formalin (3.7% formaldehyde solution) and later transferred into 75% ethanol for permanent archival storage. Preserved voucher specimens and tissues were deposited and cataloged into the fish collection at the Museum Support Center, National Museum of Natural History, Smithsonian Institution, Suitland, Maryland, USA. Nomenclature follows Randall^23^ and we followed recent taxonomic changes using the California Academy of Sciences Online Eschmeyer’s Catalog of Fishes^37^.

### DNA barcode sequencing

We extracted whole genomic DNA using QIAxtractor (QIAGEN, Crawley) and Autogen AutoGenPrep 965 according to manufacturer’s protocols. A 655bp fragment of the cytochrome oxidase I gene (COI) was amplified using Fish COI primers FISHCOILBC (TCAACYAATCAYAAAGATATYGGCAC) and FISHCOIHBC (ACTTCYGGGTGRCCRAARAATCA) and Polymerase Chain Reaction (PCR) and Sanger sequencing protocols as in Weigt et al.^38^. PCR products were Sanger sequenced bidirectionally and run on an ABI3730XL in the Laboratories of Analytical Biology (National Museum of Natural History, Smithsonian Institution). Sequences were edited using Sequencher 5.4 (Gene Codes) and aligned with Clustal W as implemented in Barcode Of Life Datasystem (BOLD, http://www.boldsystems.org). Alignments were unambiguous with no indels or frameshift mutations. A total of 2,190 DNA barcodes have been generated.

### Specimen identification

All morphological identifications were revised as needed after the specimens were deposited in the archival specimen collection to confirm initial identifications made in the field. Specimens of specific groups like Antennaridae, Bythitidae, Chlopsidae or Muraenidae were revised by additional taxonomist specialists (David Smith, John McCosker, Leslie W. Knapp, Werner Schwarzhans). After the morphological identification, we used the Taxon-ID Tree tool and Barcode Index Numbers (BIN) discordance tools as implemented in the Sequence Analysis module of BOLD to check every identification using the DNA barcodes generated. The Taxon-ID tool consists of the construction of a neighbor-joining (NJ) tree using K2P (Kimura 2 Parameter) distances by BOLD to provide a graphic representation of the species divergence^39^. The BIN discordance tool uses the Refined Single Linkage algorithm (RESL^40^) to provide a total number of OTUs.

## Data Records

This library is composed of three main components: (1) voucher specimens archived in the national fish collection at the Smithsonian Institution (Washington, DC), which were photographed in the field, (2) complete collection data associated with each voucher specimen, and (3) DNA barcodes (Fig. 1).

All photographs, voucher collection numbers, DNA barcodes and collection data are publicly available in BOLD^41^ in the Container INDOF “Fish of French Polynesia” or by scientific expedition (“AUSTR”, “GAMBA”, “MARQ”, “MOH”, “MOOP” and “SCILL”) and in Figshare^42^. DNA barcodes have also been made available in GenBank^43^ and this database is accessible through the CRIOBE portal (http://fishbardb.criobe.pf).

The library fulfills the BARCODE data standard^44,45^ which requires: 1) Species name, 2) Voucher data, 3) Collection data, 4) Identifier of the specimen, 5) COI sequence of at least 500 bp, 6) PCR primers used to generate the amplicon, 7) Trace files. In BOLD, each record in a project represents a voucher specimen with its photographs, voucher collection numbers, associated sequences and extensive collection data related to (1) the Voucher: Sample ID, Field ID, Museum ID, Institution Storing; (2) the Taxonomy: Phylum, Class, Order, Family, Subfamily, Genus, species, Identifier, Identifier E-mail, Taxonomy Notes; (3) Specimen Details: Sex, Reproduction, Life Stage, FAO Zone, Notes such as sizes of the specimens, Voucher Status, and (4) Collection Data: Collectors, Collection Date, Continent, Country/Ocean*, State/Province, Region, Sector, Exact Site, GPS Coordinates, Elevation, Depth, Depth Precision, GPS Source, and Collection Notes^42^.

## Technical Validation

To test the robustness of our library, we first computed the distribution of the interspecific and intraspecific variability for all the described species (Fig. 3b, c, d). We found that there is little to no overlap in the distribution of divergence within and between species for the vast majority of the species identified morphologically (mean intra-specific divergence 0.66, min: 0.00, max: 21.56; mean inter-specific divergence 12.28, min: 0.00, max: 24.01). The RESL algorithm identified more BINs (617) than nominal species identified morphologically (540). The morphological reexamination of specimens in light of these results suggest that 65 taxa could be new species for science awaiting a formal description (Online-only Table 1) as they are morphologically distinguishable from other species and possess unique BIN numbers. Taxonomic paraphyly (*i.e.* potentially cryptic species) has been found for 18 additional species (Table 2) as they are divided in 37 different BINs, while no morphological character has been found so far to distinguish them. Finally, mixed genealogies between sister-species were observed for 17 species (Table 3), mostly between some of the Marquesan endemics and their closest relatives that are not currently observed in the Marquesas Islands. Considering the maternal inheritance of the mitochondrial genes and the very shallow genealogies involved (maximum K2P genetic distances lower than 2%), both incomplete lineage sorting and past introgressive hybridization might be responsible of the mixing of species genealogies in those 17 cases. In summary, 94% of the BINs match species identified using morphological characters, meaning that it was possible to successfully identify a species using DNA barcodes in 94% of the cases.

**Table 2.** Potential cryptic species. Species with number of specimens collected displaying taxonomic paraphyly most likely representing undescribed cryptic species. Sample ID includes sampling location (AUST: Austral Islands, GAMB: Gambier Islands, MARQ and MOH: Marquesas Islands, SCIL and MOOP: Society Islands).

| **BINs** | **Taxa** | **No. of specimens** |
| --- | --- | --- |
| BOLD:AAF8427 | *Apogon crassiceps* | 2 |
| BOLD:ABW7007 | *Apogon crassiceps* | 4 |
| BOLD:ACE7901 | *Apogon crassiceps* | 1 |
| BOLD:ACX1964 | *Apogon doryssa* | 1 |
| BOLD:ABW8494 | *Apogon doryssa* | 2 |
| BOLD:AAF5636 | *Aporops bilinearis* | 1 |
| BOLD:AAF5637 | *Aporops bilinearis* | 4 |
| BOLD:AAD2580 | *Centropyge flavissima* | 2 |
| BOLD:AAD9019 | *Centropyge flavissima* | 6 |
| BOLD:ACD1956 | *Fusigobius duospilus* | 5 |
| BOLD:AAD1050 | *Fusigobius duospilus* | 1 |
| BOLD:AAA6306 | *Gnatholepis cauerensis* | 9 |
| BOLD:AAC6155 | *Gnatholepis cauerensis* | 5 |
| BOLD:ACC5235 | *Gymnothorax melatremus* | 3 |
| BOLD:AAC8364 | *Gymnothorax melatremus* | 5 |
| BOLD:AAF0704 | *Leiuranus semicinctus* | 3 |
| BOLD:AAL6561 | *Leiuranus semicinctus* | 2 |
| BOLD:ACD1820 | *Myrophis microchir* | 1 |
| BOLD:AAE0976 | *Myrophis microchir* | 2 |
| BOLD:AAB3862 | *Parupeneus multifasciatus* | 6 |
| BOLD:ACD1989 | *Parupeneus multifasciatus* | 3 |
| BOLD:ACD1988 | *Priolepis triops* | 3 |
| BOLD:AAX7961 | *Priolepis triops* | 1 |
| BOLD:AAB4082 | *Pristiapogon kallopterus* | 1 |
| BOLD:ABZ7996 | *Pristiapogon kallopterus* | 7 |
| BOLD:ACC5180 | *Pseudocheilinus octotaenia* | 10 |
| BOLD:AAD3038 | *Pseudocheilinus octotaenia* | 9 |
| BOLD:AAB4821 | *Pterocaesio tile* | 4 |
| BOLD:ACK9118 | *Pterocaesio tile* | 1 |
| BOLD:ACP9778 | *Scolecenchelys gymnota* | 1 |
| BOLD:AAJ8783 | *Scolecenchelys gymnota* | 2 |
| BOLD:AAC7090 | *Stegastes fasciolatus* | 11 |
| BOLD:ABZ0285 | *Stegastes fasciolatus* | 2 |
| BOLD:ACC5053 | *Uropterygius kamar* | 1 |
| BOLD:ACC5109 | *Uropterygius kamar* | 1 |
| BOLD:ACD1642 | *Uropterygius macrocephalus* | 1 |
| BOLD:AAU1965 | *Uropterygius macrocephalus* | 2 |

**Table 3.** Species displaying either incomplete lineage sorting or shallow inter-species divergence. Mean and Maximum intra-Species distances (Mean Intra-Sp and Max Intra-Sp), and Kimura 2 Parameter distances from the Nearest Neighbour (NN).

| Family | Species | Mean Intra-Sp | Max Intra-Sp | Nearest Neighbour | Nearest Species | Distance to NN |
| --- | --- | --- | --- | --- | --- | --- |
| Acanthuridae | *Acanthurus reversus* | 0.08 | 0.15 | AUSTR453-13 | *Acanthurus olivaceus* | 0 |
| Holocentridae | *Myripristis earlei* | 0.28 | 0.62 | SCILL065-15 | *Myripristis berndti* | 0 |
| Monacanthidae | *Pervagor marginalis* | 0.36 | 0.62 | SCILL083-15 | *Pervagor aspricaudus* | 0 |
| Tetraodontidae | *Canthigaster criobe* | 0 | 0 | MOH030-16 | *Canthigaster janthinoptera* | 0 |
| Mullidae | *Mulloidichthys mimicus* | 0.52 | 0.52 | AUSTR089-13 | *Mulloidichthys vanicolensis* | 0.17 |
| Pomacentridae | *Chromis abrupta* | 0 | 0 | SCILL209-15 | *Chromis margaritifer* | 0.31 |
| Labridae | *Coris marquesensis* | 0 | 0 | SCILL040-15 | *Coris gaimard* | 0.46 |
| Apogonidae | *Ostorhinchus relativus* | N/A | 0 | SCILL142-15 | *Ostorhinchus angustatus* | 0.93 |
| Tetraodontidae | *Canthigaster rapaensis* | 0.21 | 0.31 | MARQ456-12 | *Canthigaster marquesensis* | 1.1 |
| Pomacentridae | *Abudefduf conformis* | 0.15 | 0.15 | GAMBA844-12 | *Abudefduf sexfasciatus* | 1.24 |
| Monacanthidae | *Cantherhines nukuhiva* | 0.15 | 0.31 | GAMBA711-12 | *Cantherhines sandwichiensis* | 1.4 |
| Pomacentridae | *Plectroglyphidodon sagmarius* | 0.08 | 0.15 | AUSTR222-13 | *Plectroglyphidodon imparipennis* | 1.56 |
| Holocentridae | *Sargocentron caudimaculatum* | 0.68 | 1.1 | SCILL104-15 | *Sargocentron tiere* | 1.57 |
| Acanthuridae | *Zebrasoma rostratum* | 0 | 0 | AUSTR376-13 | *Zebrasoma scopas* | 1.72 |
| Apogonidae | *Apogon marquesensis* | 0.23 | 0.31 | GAMBA657-12 | *Apogon susanae* | 1.88 |
| Chaetodontidae | *Chaetodon flavirostris* | 0.08 | 0.15 | SCILL269-15 | *Chaetodon lunula* | 1.88 |
| Chaetodontidae | *Chaetodon lunula* | 0.1 | 0.15 | GAMBA555-12 | *Chaetodon flavirostris* | 1.88 |

## Usage Notes

This Barcode release dataset is freely available to use in barcoding or metabarcoding surveys for specimen identification. Several approaches can be considered:

1. directly downloading the sequences in fasta format, and working offline by merging this dataset with an ongoing barcoding project;
2. working online, through the BOLD website (registration is free), and merging the Container INDOF “Fish of French Polynesia” or parts of the scientific expeditions (Table 1) with an ongoing BOLD project;
3. through online identification tools, as data are indexed in both BOLD and Genbank databases. This library will be considered when any queries of molecular identification will be made through the identification engine of BOLD (http://www.boldsystems.org/index.php/IDS-OpenIdEngine) or the standard nucleotide Basic Local Alignment Search Tool (BLAST, https://blast.ncbi.nlm.nih.gov/). In the same manner, this dataset should also be indexed in the MIDORI database^46,47^. Composed of both endemic and widespread species, this library is expected to benefit a large community from academics to authorities who use molecular data to monitor and survey biodiversity.

## Acknowledgements

This work was financially supported by the French National Agency for Marine Protected Area in France (‘Pakaihi I Te Moana’ expedition), the ANR IMODEL and Contrat de Projet Etat-Territoire in French Polynesia and the French Ministry for Environment, Sustainable Development and Transport (MEDDTL) (‘CORALSPOT’ expeditions), the Living Oceans Foundation (‘Australs’ expedition) and the LabEx CORAIL and the GOPS, (‘Scilly’ expedition) and the Gordon and Betty Moore Foundation (Mo‘orea Biocode Project). Additional funding was provided by the IFRECOR in French Polynesia and the TOTAL Foundation. We are grateful to T. Frogier, P. Mery and the Centre Plongée Marquises (Xavier (Pipapo) and Marie Curvat), for their field assistance in the Gambier, the Marquesas, the Australs and Scilly Islands along with the crew of the *Claymore II, Braveheart* and the *Golden Shadow*. We thank the Ministère de l’Environnement de Polynésie, the Délégation à la Recherche Polynésie, the Mairie of Nuku-Hiva, and the people of the Marquesas Islands for their kind and generous support of the project as we traveled throughout the islands. We thank Jerry Finan, Erika Wilbur, Shirleen Smith, Kris Murphy, David Smith and Sandra Raredon of the Division of Fishes (National Museum of Natural History) for assistance in preparations for the trip and processing specimens and Jeffrey Hunt and Kenneth Macdonald III and Meaghan Parker Forney of the Laboratories of Analytical Biology (Smithsonian Institution) for assistance in molecular analysis of samples. Finally, we thank the staff of the CRIOBE and particularly Yannick Chancerelle for logistical support in French Polynesia. Specimens were collected under the permit “Permanent agreement, Délégation à la Recherche, French Polynesia”.

## Author contributions

E.D.T. drafted the first manuscript, J.T.W., D.P., A.D., and N.H. provided extensive edits and comments. E.D.T, J.T.W., T.C., R.G., M.K., T.L.M., J.M., G.M.-T., V.P., P.P., P.S., G.S., N.T., M.V. and S.P. collected fish specimens. E.D.T., J.T.W., D.P., A.D., N.H., J.V., B.E., C. M., L.W. and S.P. produced DNA barcodes and cleaned the database. R.G. and S.P. financed the scientific expeditions. C.M. and S.P. financed the sequencing. All authors read and approved the final manuscript.

---

**Online-only Table 1.**
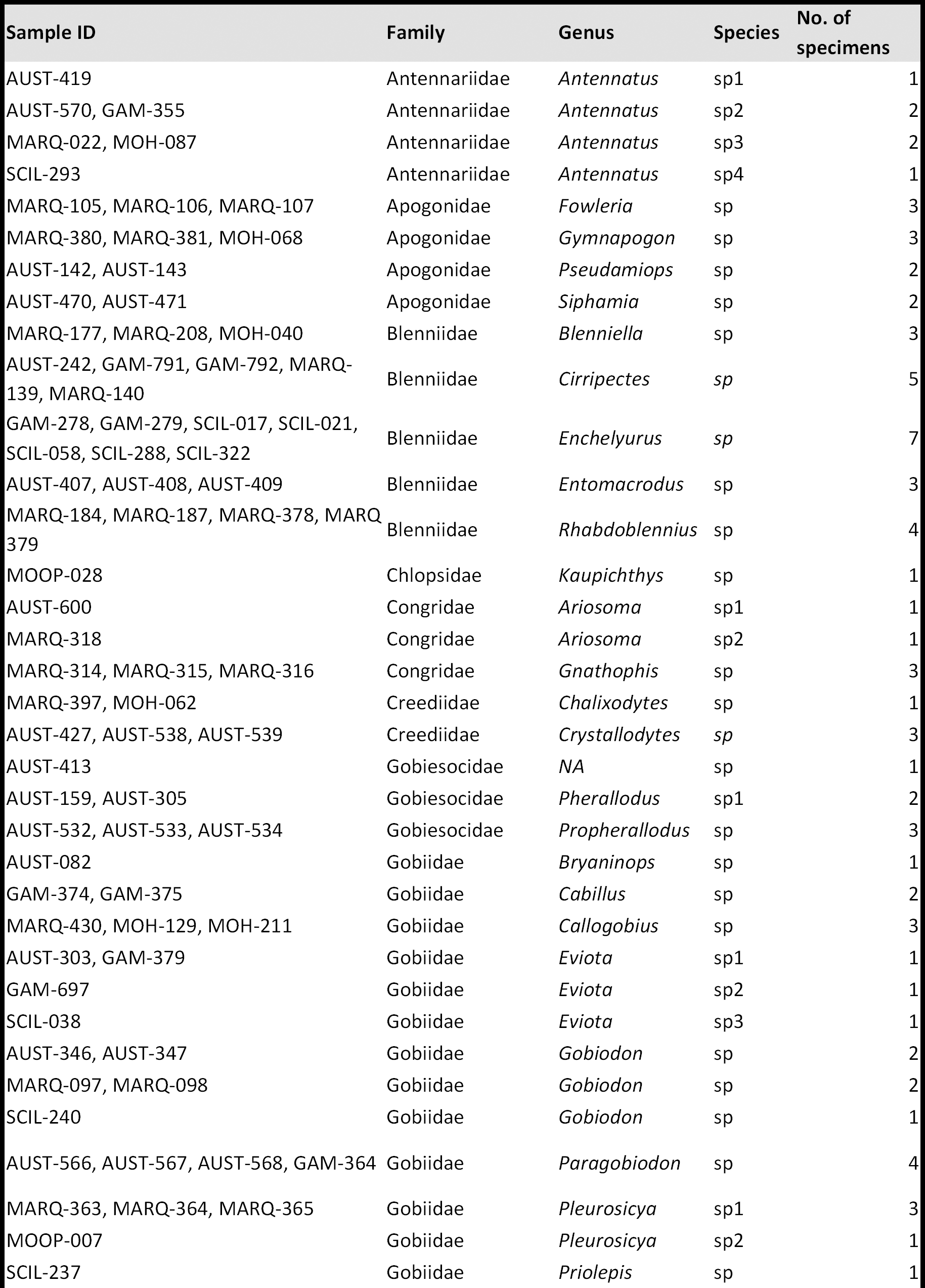
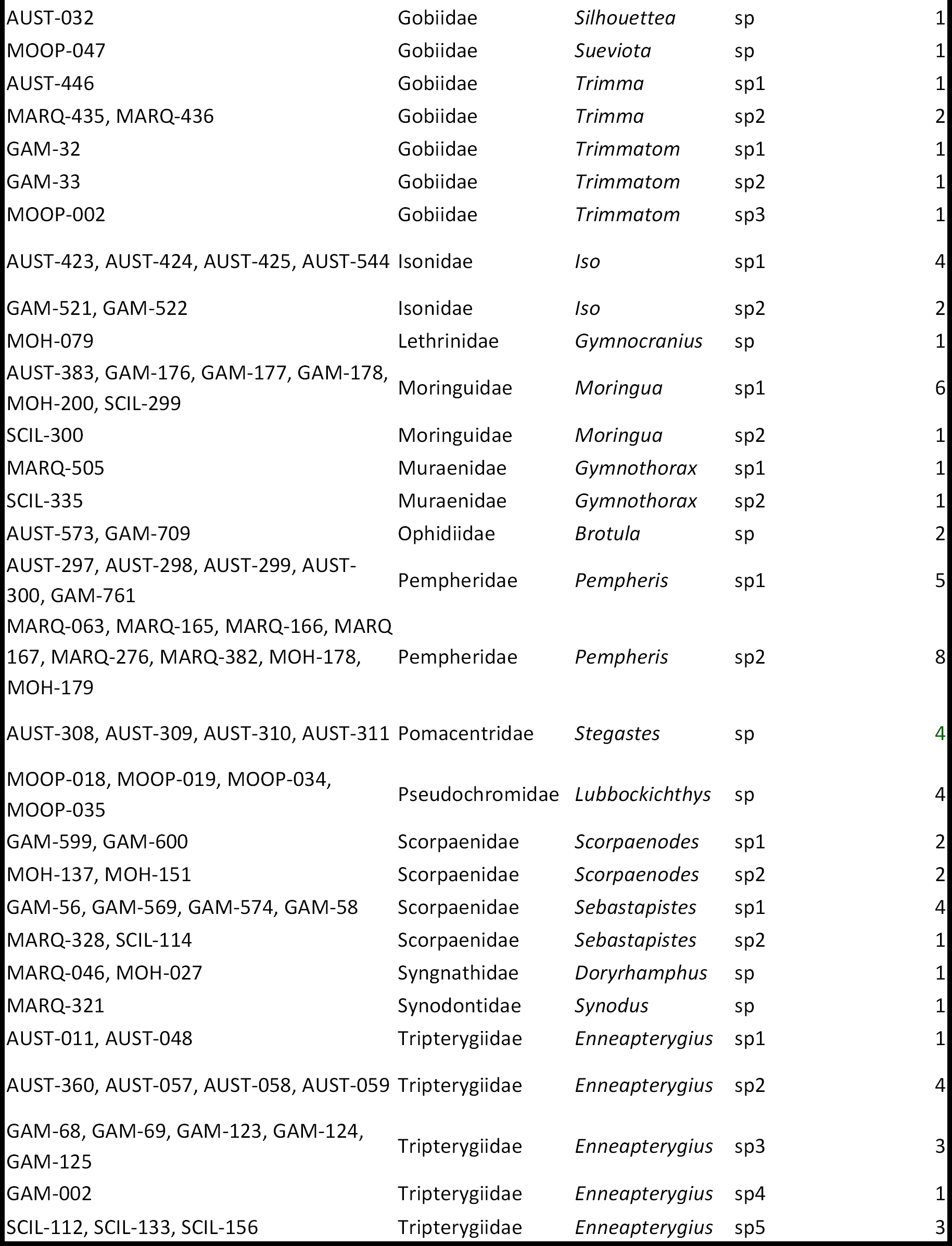
Potential new species detected in the dataset. Specimens which were identified only to the genus level and which represent potentially new species waiting to be described. Number of specimens included in each Barcode Index Number (BIN).

## References

1. Hebert, P. D. N. & Gregory, T. R. The promise of DNA barcoding for taxonomy. Syst. Biol. 54, 852–859 (2005).

2. Garnett, S. T. & Christidis, L. Taxonomy anarchy hampers conservation. Nat. News 546, 25 (2017).

3. Hubert, N., Delrieu-Trottin, E., Irisson, J. O., Meyer, C. & Planes, S. Identifying coral reef fish larvae through DNA barcoding: A test case with the families Acanthuridae and Holocentridae. Mol. Phylogenet. Evol. 55, 1195–1203 (2010).

4. Ko, H.-L. et al. Evaluating the accuracy of morphological identification of larval fishes by applying DNA barcoding. PLoS One 8, e53451 (2013).

5. Wong, E. H.-K. & Hanner, R. H. DNA barcoding detects market substitution in North American seafood. Food Res. Int. 41, 828–837 (2008).

6. Holmes, B. H., Steinke, D. & Ward, R. D. Identification of shark and ray fins using DNA barcoding. Fish. Res. 95, 280–288 (2009).

7. Riedel, A., Sagata, K., Suhardjono, Y. R., Tänzler, R. & Balke, M. Integrative taxonomy on the fast track-towards more sustainability in biodiversity research. Front. Zool. 10, 15 (2013).

8. Monaghan, M. T. et al. Accelerated species inventory on Madagascar using coalescent-based models of species delineation. Syst. Biol. 58, 298–311 (2009).

9. Tänzler, R., Sagata, K., Surbakti, S., Balke, M. & Riedel, A. DNA barcoding for community ecology-how to tackle a hyperdiverse, mostly undescribed Melanesian fauna. PLoS One 7, e28832 (2012).

10. Hubert, N. & Hanner, R. DNA barcoding, species delineation and taxonomy: a historical perspective. DNA barcodes 3, 44–58 (2015).

11. Leray, M. & Knowlton, N. DNA barcoding and metabarcoding of standardized samples reveal patterns of marine benthic diversity. Proc. Natl. Acad. Sci. 112, 2076–2081 (2015).

12. Leray, M., Boehm, J. T., Mills, S. C. & Meyer, C. P. Moorea BIOCODE barcode library as a tool for understanding predator-prey interactions: insights into the diet of common predatory coral reef fishes. Coral Reefs 31, 383–388 (2012).

13. Leray, M., Meyer, C. P. & Mills, S. C. Metabarcoding dietary analysis of coral dwelling predatory fish demonstrates the minor contribution of coral mutualists to their highly partitioned, generalist diet. PeerJ 3, e1047 (2015).

14. Bakker, J. et al. Environmental DNA reveals tropical shark diversity in contrasting levels of anthropogenic impact. Sci. Rep. 7, 16886 (2017).

15. Stat, M. et al. Ecosystem biomonitoring with eDNA: metabarcoding across the tree of life in a tropical marine environment. Sci. Rep. 7, 12240 (2017).

16. DiBattista, J. D. et al. Assessing the utility of eDNA as a tool to survey reef-fish communities in the Red Sea. Coral Reefs 36, 1245–1252 (2017).

17. Hebert, P. D. N. et al. A DNA ‘Barcode Blitz’: Rapid digitization and sequencing of a natural history collection. PLoS One 8, e68535 (2013).

18. Schmidt, S., Schmid-Egger, C., Morinière, J., Haszprunar, G. & Hebert, P. D. N. DNA barcoding largely supports 250 years of classical taxonomy: identifications for Central European bees (Hymenoptera, Apoidea *partim*). Mol. Ecol. Resour. 15, 985–1000 (2015).

19. Hubert, N. et al. Cryptic diversity in indo-pacific coral-reef fishes revealed by DNA-barcoding provides new support to the centre-of-overlap hypothesis. PLoS One 7, (2012).

20. Lamy, T., Legendre, P., Chancerelle, Y., Siu, G. & Claudet, J. Understanding the spatio-temporal response of coral reef fish communities to natural disturbances: insights from beta-diversity decomposition. PLoS One 10, e0138696 (2015).

21. Galzin, R. et al. Long term monitoring of coral and fish assemblages (1983 - 2014) in Tiahura reefs, Moorea, French Polynesia. Cybium 40, 31–41 (2016).

22. Delrieu-Trottin, E. et al. Shore fishes of the Marquesas Islands, an updated checklist with new records and new percentage of endemic species. Check List 11, (2015).

23. Randall, J. E. Reef and Shore Fishes of the South Pacific: New Caledonia to Tahiti and the Pitcairn Islands. 1, (University of Hawai’i Press Honolulu, 2005)

24. Randall, J. E. & Cea, A. Shore Fishes of Easter Island. (University of Hawai’i Press, 2011)

25. Delrieu-Trottin, E. et al. Evidence of cryptic species in the blenniid *Cirripectes alboapicalis* species complex, with zoogeographic implications for the South Pacific. Zookeys 810, 127–138 (2018).

26. Siu, G. et al. Shore fishes of French polynesia. Cybium 41, 1–34 (2017).

27. Williams, J. T., Delrieu-Trottin, E. & Planes, S. A new species of Indo-Pacific fish, *Canthigaster criobe*, with comments on other Canthigaster (Tetraodontiformes: Tetraodontidae) at the Gambier Archipelago. Zootaxa 3523, 80–88 (2012).

28. Tornabene, L., Ahmadia, G. N. & Williams, J. T. Four new species of dwarfgobies (Teleostei: Gobiidae: *Eviota*) from the Austral, Gambier, Marquesas and Society Archipelagos, French Polynesia. Syst. Biodivers. 11, 363–380 (2013).

29. Williams, J. T., Delrieu-Trottin, E. & Planes, S. Two new fish species of the subfamily Anthiinae (Perciformes, Serranidae) from the Marquesas. Zootaxa 3647, 167–180 (2013).

30. Delrieu-Trottin, E., Williams, J. T. & Planes, S. *Macropharyngodon pakoko*, a new species of wrasse (Teleostei: Labridae) endemic to the Marquesas Islands, French polynesia. Zootaxa 3857, 433–443 (2014).

31. McCosker, J. E. & Hibino, Y. A review of the finless snake eels of the genus *Apterichtus* (Anguilliformes: Ophichthidae), with the description of five new species. Zootaxa 3941, 49–78 (2015).

32. Polanco, F. A., Acero, P. A. & Betancur-R., R. No longer a circumtropical species: revision of the lizardfishes in the *Trachinocephalus myops* species complex, with description of a new species from the Marquesas Islands. J. Fish Biol. 89, 1302–1323 (2016).

33. Viviani, J., Williams, J. T. & Planes, S. Two new pygmygobies (Percomorpha: Gobiidae: *Trimma*) from French Polynesia. J. Ocean Sci. Found. 23, 1–11 (2016).

34. Williams, J. T. & Viviani, J. *Pseudogramma polyacantha* complex (Serranidae, tribe Grammistini): DNA barcoding results lead to the discovery of three cryptic species, including two new species from French Polynesia. Zootaxa 4111, 246–260 (2016).

35. Williams, J. T. et al. Checklist of the shorefishes of Wallis Islands (Wallis and Futuna French Territories, South-Central Pacific). Cybium 30, 247–260 (2006).

36. Bacchet, P., Zysman, T. & Lefevre, Y. Guide des poissons de Tahiti et ses iles. (Au vent des îles, 2006)

37. Eschmeyer, W. N., Fricke, R. & van der Laan, R. Eschmeyer’s Catalog of Fishes electronic version. http://researcharchive.calacademy.org/research/Ichthyology/catalog/fishcatmain.asp (2018).

38. Weigt, L. A., Driskell, A. C., Baldwin, C. C. & Ormos, A. in DNA barcodes 109–126 (Springer, 2012)

39. Ratnasingham, S. & Hebert, P. D. N. BOLD: The Barcode of Life Data System (www.barcodinglife.org). Mol. Ecol. Notes 7, p355–364 (2007).

40. Ratnasingham, S. & Hebert, P. D. N. A DNA-based registry for all animal species: the Barcode Index Number (BIN) system. PLoS One 8, e66213 (2013).

41. BOLD DS-INDOF https://doi.org/10.5883/DS-INDOF (2019).

42. Delrieu-Trottin et al. Figshare https://doi.org/10.6084/m9.figshare.7923905 (2019).

43. NCBI Nucleotide https://www.ncbi.nlm.nih.gov/nucleotide/ Accession numbers KC567661 - KC567663, KC684990, KC684991, KU905709-KU905727, KY570698, KY570703 - KY570705, KY570708, KY683549, MH707846 - MH707881, MK566774 - MK567153, MK656969 - MK658713 (2019)

44. Hubert, N. et al. Identifying Canadian freshwater fishes through DNA barcodes. PLoS One 3, (2008).

45. Hubert, N. & Hanner, R. DNA Barcoding, species delineation and taxonomy: a historical perspective. DNA Barcodes 3, 44–58 (2015).

46. Machida, R. J., Leray, M., Ho, S. L. & Knowlton, N. Metazoan mitochondrial gene sequence reference datasets for taxonomic assignment of environmental samples. Sci. Data 4, 1–7 (2017).

47. Leray, M., Ho, S. L., Lin, I. J. & Machida, R. J. MIDORI server: a webserver for taxonomic assignment of unknown metazoan mitochondrial-encoded sequences using a curated database. Bioinformatics 34, 3753–3754 (2018).

