## Supplementary Figure for "A DNA barcode reference library of the French Polynesian shore fishes"

### BOLD TaxonID Tree

| | | |
|---|---|---|
| Title | : | Tree Result - DS-INDOF (2192 records selected) |
| Date | : | 22-Mar-2019 |
| Data Type | : | Nucleotide |
| Distance Model | : | Kimura 2 Parameter |
| Marker | : | COI-5P |
| Colourization | : | [blue]=Stop Codons [red]=Contamination or misidentification |
| Label | : | Sample ID |
| Label | : | Process ID |
| Label | : | Taxon |
| Label | : | GenBank Accession |
| Sequence Count | : | 2190 |
| Species count | : | 540 |
| Genus count | : | 232 |
| Family count | : | 61 |
| Unidentified | : | 157 |
| BIN Count | : | 617 |

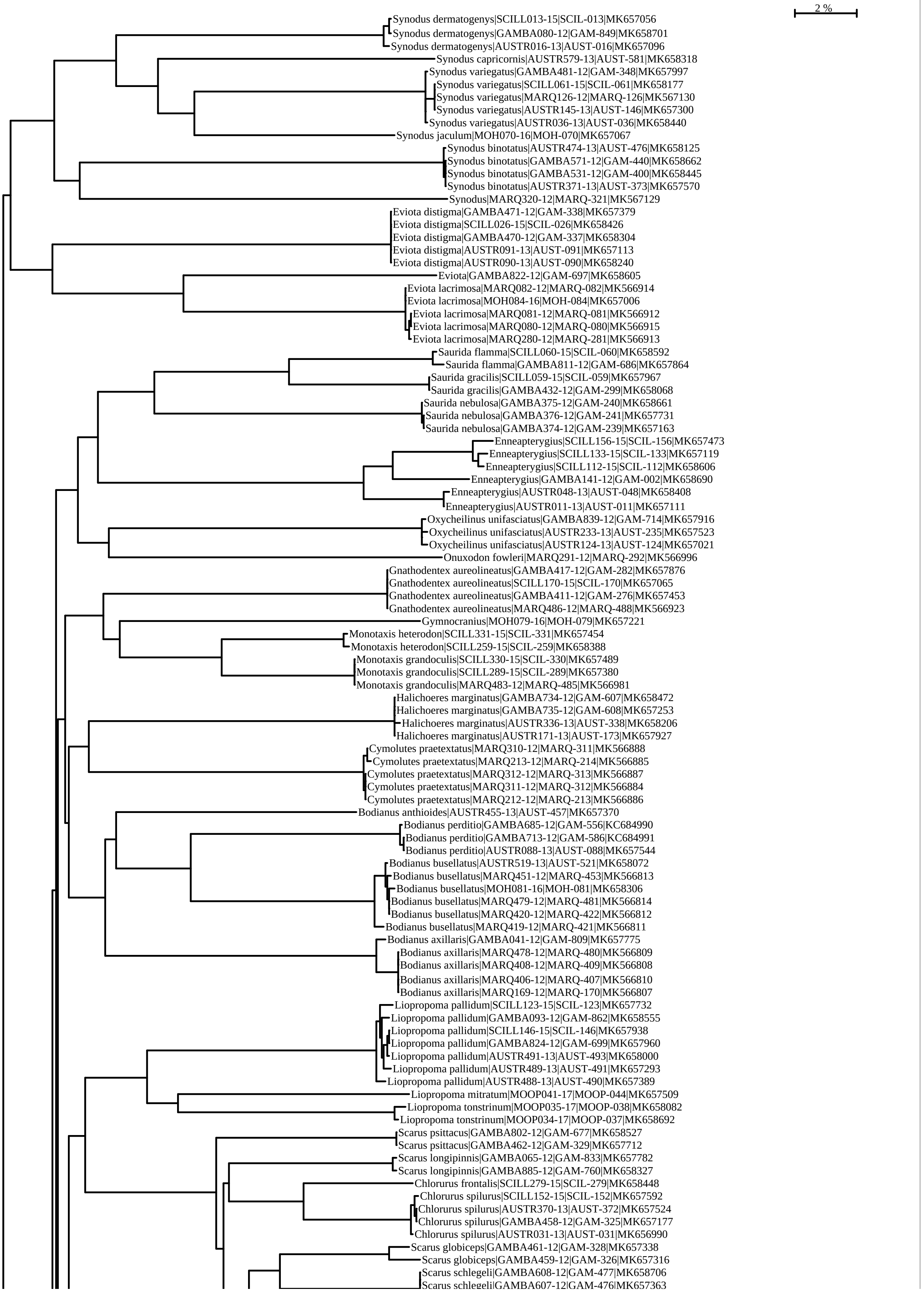
2 %
Synodus dermatogenys|SCILL013-15|SCIL-013|MK657056
Synodus dermatogenys|GAMBA080-12|GAM-849|MK658701
Synodus dermatogenys|AUSTR016-13|AUST-016|MK657096
Synodus capricornis|AUSTR579-13|AUST-581|MK658318
Synodus variegatus|GAMBA481-12|GAM-348|MK657997
Synodus variegatus|SCILL061-15|SCIL-061|MK658177
Synodus variegatus|MARQ126-12|MARQ-126|MK567130
Synodus variegatus|AUSTR145-13|AUST-146|MK657300
Synodus variegatus|AUSTR036-13|AUST-036|MK658440
Synodus jaculum|MOH070-16|MOH-070|MK657067
Synodus binotatus|AUSTR474-13|AUST-476|MK658125
Synodus binotatus|GAMBA571-12|GAM-440|MK658662
Synodus binotatus|GAMBA531-12|GAM-400|MK658445
Synodus binotatus|AUSTR371-13|AUST-373|MK657570
Synodus|MARQ320-12|MARQ-321|MK567129
Eviota distigma|GAMBA471-12|GAM-338|MK657379
Eviota distigma|SCILL026-15|SCIL-026|MK658426
Eviota distigma|GAMBA470-12|GAM-337|MK658304
Eviota distigma|AUSTR091-13|AUST-091|MK657113
Eviota distigma|AUSTR090-13|AUST-090|MK658240
Eviota|GAMBA822-12|GAM-697|MK658605
Eviota lacrimosa|MARQ082-12|MARQ-082|MK566914
Eviota lacrimosa|MOH084-16|MOH-084|MK657006
Eviota lacrimosa|MARQ081-12|MARQ-081|MK566912
Eviota lacrimosa|MARQ080-12|MARQ-080|MK566915
Eviota lacrimosa|MARQ280-12|MARQ-281|MK566913
Saurida flamma|SCILL060-15|SCIL-060|MK658592
Saurida flamma|GAMBA811-12|GAM-686|MK657864
Saurida gracilis|SCILL059-15|SCIL-059|MK657967
Saurida gracilis|GAMBA432-12|GAM-299|MK658068
Saurida nebulosa|GAMBA375-12|GAM-240|MK658661
Saurida nebulosa|GAMBA376-12|GAM-241|MK657731
Saurida nebulosa|GAMBA374-12|GAM-239|MK657163
Enneapterygius|SCILL156-15|SCIL-156|MK657473
Enneapterygius|SCILL133-15|SCIL-133|MK657119
Enneapterygius|SCILL112-15|SCIL-112|MK658606
Enneapterygius|GAMBA141-12|GAM-002|MK658690
Enneapterygius|AUSTR048-13|AUST-048|MK658408
Enneapterygius|AUSTR011-13|AUST-011|MK657111
Oxycheilinus unifasciatus|GAMBA839-12|GAM-714|MK657916
Oxycheilinus unifasciatus|AUSTR233-13|AUST-235|MK657523
Oxycheilinus unifasciatus|AUSTR124-13|AUST-124|MK657021
Onuxodon fowleri|MARQ291-12|MARQ-292|MK566996
Gnathodentex aureolineatus|GAMBA417-12|GAM-282|MK657876
Gnathodentex aureolineatus|SCILL170-15|SCIL-170|MK657065
Gnathodentex aureolineatus|GAMBA411-12|GAM-276|MK657453
Gnathodentex aureolineatus|MARQ486-12|MARQ-488|MK566923
Gymnocranius|MOH079-16|MOH-079|MK657221
Monotaxis heterodon|SCILL331-15|SCIL-331|MK657454
Monotaxis heterodon|SCILL259-15|SCIL-259|MK658388
Monotaxis grandoculis|SCILL330-15|SCIL-330|MK657489
Monotaxis grandoculis|SCILL289-15|SCIL-289|MK657380
Monotaxis grandoculis|MARQ483-12|MARQ-485|MK566981
Halichoeres marginatus|GAMBA734-12|GAM-607|MK658472
Halichoeres marginatus|GAMBA735-12|GAM-608|MK657253
Halichoeres marginatus|AUSTR336-13|AUST-338|MK658206
Halichoeres marginatus|AUSTR171-13|AUST-173|MK657927
Cymolutes praetextatus|MARQ310-12|MARQ-311|MK566888
Cymolutes praetextatus|MARQ213-12|MARQ-214|MK566885
Cymolutes praetextatus|MARQ312-12|MARQ-313|MK566887
Cymolutes praetextatus|MARQ311-12|MARQ-312|MK566884
Cymolutes praetextatus|MARQ212-12|MARQ-213|MK566886
Bodianus anthioides|AUSTR455-13|AUST-457|MK657370
Bodianus perditio|GAMBA685-12|GAM-556|KC684990
Bodianus perditio|GAMBA713-12|GAM-586|KC684991
Bodianus perditio|AUSTR088-13|AUST-088|MK657544
Bodianus busellatus|AUSTR519-13|AUST-521|MK658072
Bodianus busellatus|MARQ451-12|MARQ-453|MK566813
Bodianus busellatus|MOH081-16|MOH-081|MK658306
Bodianus busellatus|MARQ479-12|MARQ-481|MK566814
Bodianus busellatus|MARQ420-12|MARQ-422|MK566812
Bodianus busellatus|MARQ419-12|MARQ-421|MK566811
Bodianus axillaris|GAMBA041-12|GAM-809|MK657775
Bodianus axillaris|MARQ478-12|MARQ-480|MK566809
Bodianus axillaris|MARQ408-12|MARQ-409|MK566808
Bodianus axillaris|MARQ406-12|MARQ-407|MK566810
Bodianus axillaris|MARQ169-12|MARQ-170|MK566807
Liopropoma pallidum|SCILL123-15|SCIL-123|MK657732
Liopropoma pallidum|GAMBA093-12|GAM-862|MK658555
Liopropoma pallidum|SCILL146-15|SCIL-146|MK657938
Liopropoma pallidum|GAMBA824-12|GAM-699|MK657960
Liopropoma pallidum|AUSTR491-13|AUST-493|MK658000
Liopropoma pallidum|AUSTR489-13|AUST-491|MK657293
Liopropoma pallidum|AUSTR488-13|AUST-490|MK657389
Liopropoma mitratum|MOOP041-17|MOOP-044|MK657509
Liopropoma tonstrinum|MOOP035-17|MOOP-038|MK658082
Liopropoma tonstrinum|MOOP034-17|MOOP-037|MK658692
Scarus psittacus|GAMBA802-12|GAM-677|MK658527
Scarus psittacus|GAMBA462-12|GAM-329|MK657712
Scarus longipinnis|GAMBA065-12|GAM-833|MK657782
Scarus longipinnis|GAMBA885-12|GAM-760|MK658327
Chlorurus frontalis|SCILL279-15|SCIL-279|MK658448
Chlorurus spilurus|SCILL152-15|SCIL-152|MK657592
Chlorurus spilurus|AUSTR370-13|AUST-372|MK657524
Chlorurus spilurus|GAMBA458-12|GAM-325|MK657177
Chlorurus spilurus|AUSTR031-13|AUST-031|MK656990
Scarus globiceps|GAMBA461-12|GAM-328|MK657338
Scarus globiceps|GAMBA459-12|GAM-326|MK657316
Scarus schlegeli|GAMBA608-12|GAM-477|MK658706
Scarus schlegeli|GAMBA607-12|GAM-476|MK657363

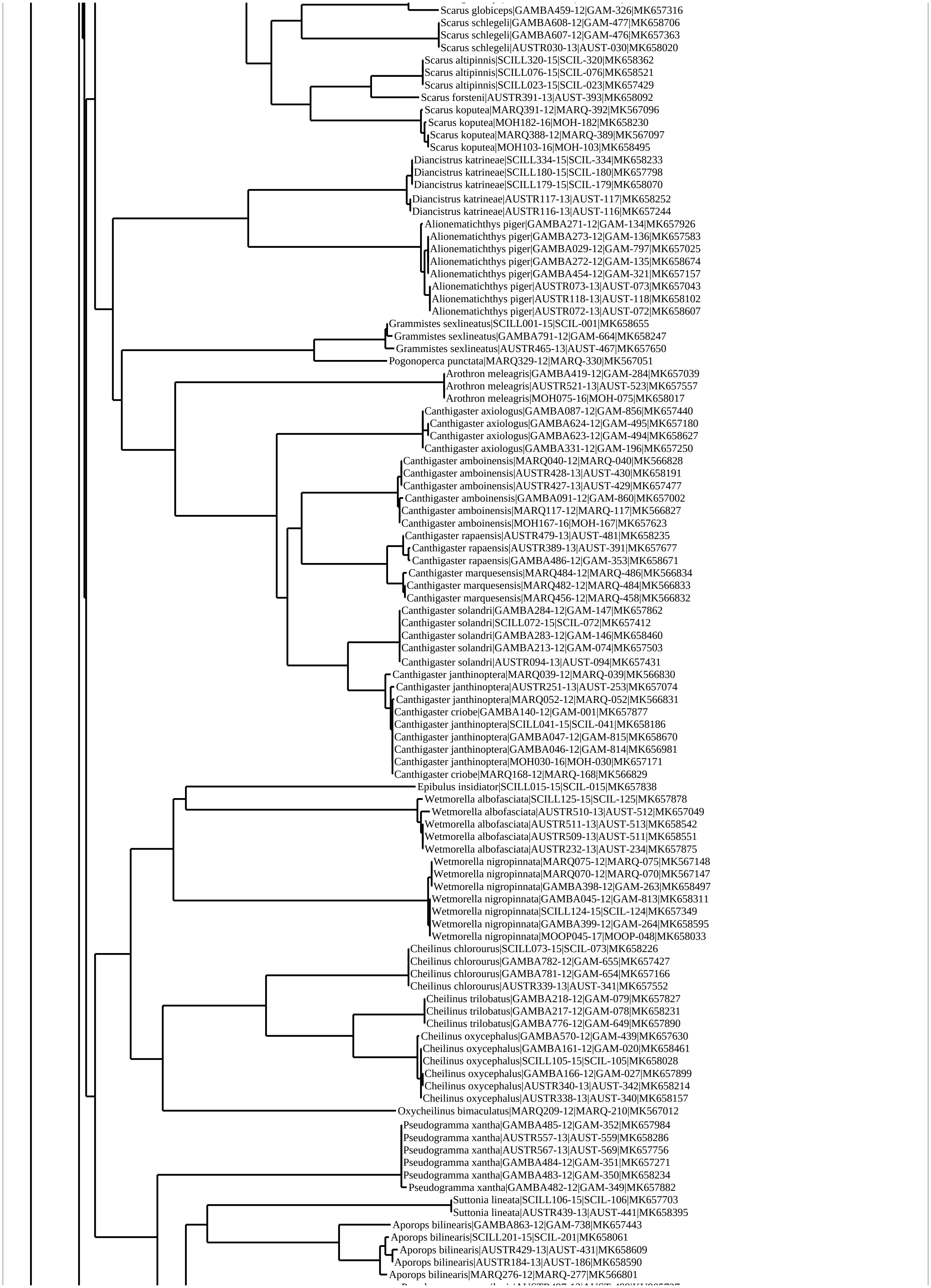
Scarus globiceps|GAMBA459-12|GAM-326|MK657316
Scarus schlegeli|GAMBA608-12|GAM-477|MK658706
Scarus schlegeli|GAMBA607-12|GAM-476|MK657363
Scarus schlegeli|AUSTR030-13|AUST-030|MK658020
Scarus altipinnis|SCILL320-15|SCIL-320|MK658362
Scarus altipinnis|SCILL076-15|SCIL-076|MK658521
Scarus altipinnis|SCILL023-15|SCIL-023|MK657429
Scarus forsteni|AUSTR391-13|AUST-393|MK658092
Scarus koputea|MARQ391-12|MARQ-392|MK567096
Scarus koputea|MOH182-16|MOH-182|MK658230
Scarus koputea|MARQ388-12|MARQ-389|MK567097
Scarus koputea|MOH103-16|MOH-103|MK658495
Diancistrus katrineae|SCILL334-15|SCIL-334|MK658233
Diancistrus katrineae|SCILL180-15|SCIL-180|MK657798
Diancistrus katrineae|SCILL179-15|SCIL-179|MK658070
Diancistrus katrineae|AUSTR117-13|AUST-117|MK658252
Diancistrus katrineae|AUSTR116-13|AUST-116|MK657244
Alionematichthys piger|GAMBA271-12|GAM-134|MK657926
Alionematichthys piger|GAMBA273-12|GAM-136|MK657583
Alionematichthys piger|GAMBA029-12|GAM-797|MK657025
Alionematichthys piger|GAMBA272-12|GAM-135|MK658674
Alionematichthys piger|GAMBA454-12|GAM-321|MK657157
Alionematichthys piger|AUSTR073-13|AUST-073|MK657043
Alionematichthys piger|AUSTR118-13|AUST-118|MK658102
Alionematichthys piger|AUSTR072-13|AUST-072|MK658607
Grammistes sexlineatus|SCILL001-15|SCIL-001|MK658655
Grammistes sexlineatus|GAMBA791-12|GAM-664|MK658247
Grammistes sexlineatus|AUSTR465-13|AUST-467|MK657650
Pogonoperca punctata|MARQ329-12|MARQ-330|MK567051
Arothron meleagris|GAMBA419-12|GAM-284|MK657039
Arothron meleagris|AUSTR521-13|AUST-523|MK657557
Arothron meleagris|MOH075-16|MOH-075|MK658017
Canthigaster axiologus|GAMBA087-12|GAM-856|MK657440
Canthigaster axiologus|GAMBA624-12|GAM-495|MK657180
Canthigaster axiologus|GAMBA623-12|GAM-494|MK658627
Canthigaster axiologus|GAMBA331-12|GAM-196|MK657250
Canthigaster amboinensis|MARQ040-12|MARQ-040|MK566828
Canthigaster amboinensis|AUSTR428-13|AUST-430|MK658191
Canthigaster amboinensis|AUSTR427-13|AUST-429|MK657477
Canthigaster amboinensis|GAMBA091-12|GAM-860|MK657002
Canthigaster amboinensis|MARQ117-12|MARQ-117|MK566827
Canthigaster amboinensis|MOH167-16|MOH-167|MK657623
Canthigaster rapaensis|AUSTR479-13|AUST-481|MK658235
Canthigaster rapaensis|AUSTR389-13|AUST-391|MK657677
Canthigaster rapaensis|GAMBA486-12|GAM-353|MK658671
Canthigaster marquesensis|MARQ484-12|MARQ-486|MK566834
Canthigaster marquesensis|MARQ482-12|MARQ-484|MK566833
Canthigaster marquesensis|MARQ456-12|MARQ-458|MK566832
Canthigaster solandri|GAMBA284-12|GAM-147|MK657862
Canthigaster solandri|SCILL072-15|SCIL-072|MK657412
Canthigaster solandri|GAMBA283-12|GAM-146|MK658460
Canthigaster solandri|GAMBA213-12|GAM-074|MK657503
Canthigaster solandri|AUSTR094-13|AUST-094|MK657431
Canthigaster janthinoptera|MARQ039-12|MARQ-039|MK566830
Canthigaster janthinoptera|AUSTR251-13|AUST-253|MK657074
Canthigaster janthinoptera|MARQ052-12|MARQ-052|MK566831
Canthigaster criobe|GAMBA140-12|GAM-001|MK657877
Canthigaster janthinoptera|SCILL041-15|SCIL-041|MK658186
Canthigaster janthinoptera|GAMBA047-12|GAM-815|MK658670
Canthigaster janthinoptera|GAMBA046-12|GAM-814|MK656981
Canthigaster janthinoptera|MOH030-16|MOH-030|MK657171
Canthigaster criobe|MARQ168-12|MARQ-168|MK566829
Epibulus insidiator|SCILL015-15|SCIL-015|MK657838
Wetmorella albofasciata|SCILL125-15|SCIL-125|MK657878
Wetmorella albofasciata|AUSTR510-13|AUST-512|MK657049
Wetmorella albofasciata|AUSTR511-13|AUST-513|MK658542
Wetmorella albofasciata|AUSTR509-13|AUST-511|MK658551
Wetmorella albofasciata|AUSTR232-13|AUST-234|MK657875
Wetmorella nigropinnata|MARQ075-12|MARQ-075|MK567148
Wetmorella nigropinnata|MARQ070-12|MARQ-070|MK567147
Wetmorella nigropinnata|GAMBA398-12|GAM-263|MK658497
Wetmorella nigropinnata|GAMBA045-12|GAM-813|MK658311
Wetmorella nigropinnata|SCILL124-15|SCIL-124|MK657349
Wetmorella nigropinnata|GAMBA399-12|GAM-264|MK658595
Wetmorella nigropinnata|MOOP045-17|MOOP-048|MK658033
Cheilinus chlorourus|SCILL073-15|SCIL-073|MK658226
Cheilinus chlorourus|GAMBA782-12|GAM-655|MK657427
Cheilinus chlorourus|GAMBA781-12|GAM-654|MK657166
Cheilinus chlorourus|AUSTR339-13|AUST-341|MK657552
Cheilinus trilobatus|GAMBA218-12|GAM-079|MK657827
Cheilinus trilobatus|GAMBA217-12|GAM-078|MK658231
Cheilinus trilobatus|GAMBA776-12|GAM-649|MK657890
Cheilinus oxycephalus|GAMBA570-12|GAM-439|MK657630
Cheilinus oxycephalus|GAMBA161-12|GAM-020|MK658461
Cheilinus oxycephalus|SCILL105-15|SCIL-105|MK658028
Cheilinus oxycephalus|GAMBA166-12|GAM-027|MK657899
Cheilinus oxycephalus|AUSTR340-13|AUST-342|MK658214
Cheilinus oxycephalus|AUSTR338-13|AUST-340|MK658157
Oxycheilinus bimaculatus|MARQ209-12|MARQ-210|MK567012
Pseudogramma xantha|GAMBA485-12|GAM-352|MK657984
Pseudogramma xantha|AUSTR557-13|AUST-559|MK658286
Pseudogramma xantha|AUSTR567-13|AUST-569|MK657756
Pseudogramma xantha|GAMBA484-12|GAM-351|MK657271
Pseudogramma xantha|GAMBA483-12|GAM-350|MK658234
Pseudogramma xantha|GAMBA482-12|GAM-349|MK657882
Suttonia lineata|SCILL106-15|SCIL-106|MK657703
Suttonia lineata|AUSTR439-13|AUST-441|MK658395
Aporops bilinearis|GAMBA863-12|GAM-738|MK657443
Aporops bilinearis|SCILL201-15|SCIL-201|MK658061
Aporops bilinearis|AUSTR429-13|AUST-431|MK658609
Aporops bilinearis|AUSTR184-13|AUST-186|MK658590
Aporops bilinearis|MARQ276-12|MARQ-277|MK566801

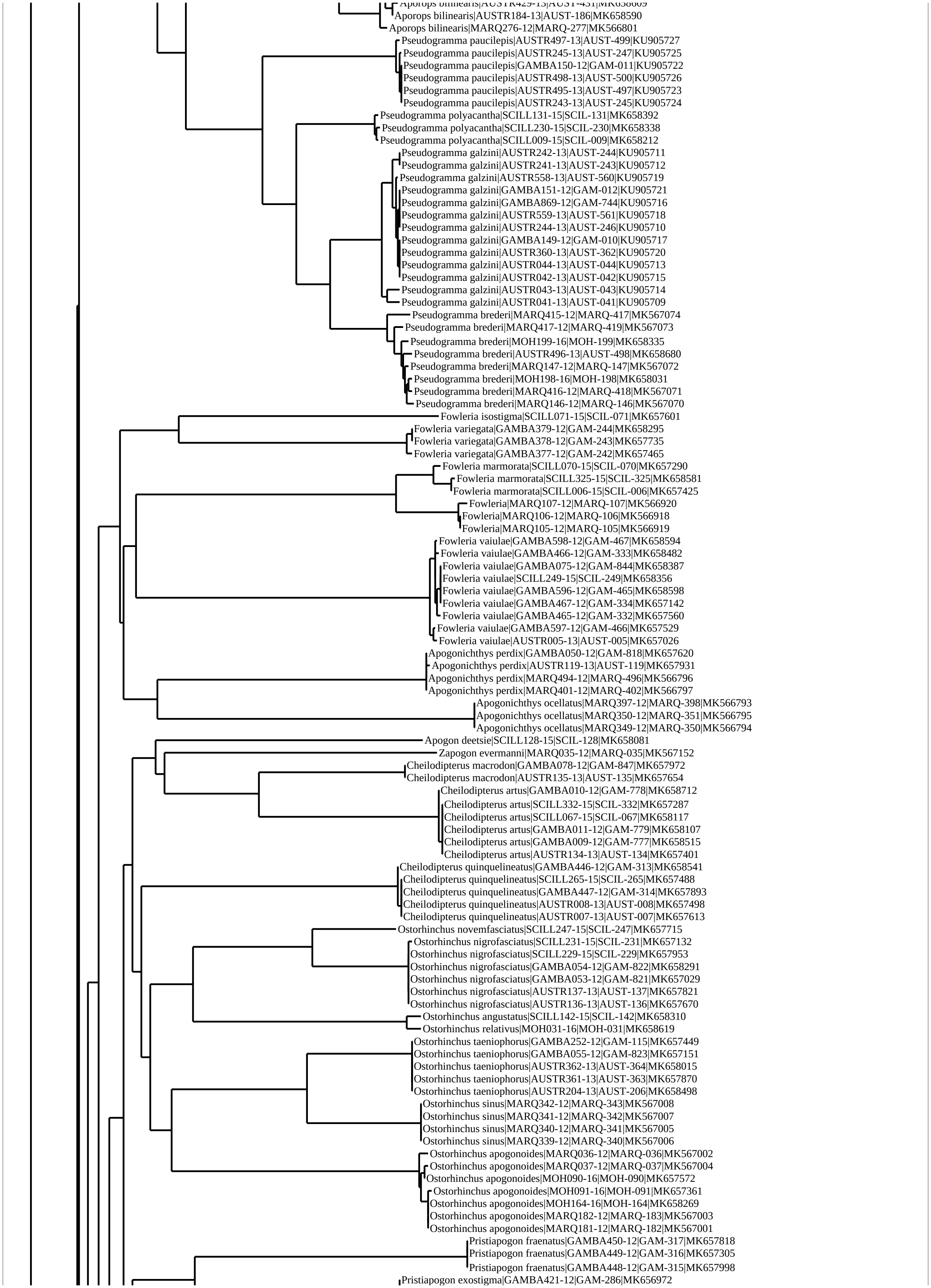
Aporops bilinearis|AUSTR184-13|AUST-186|MK658590
Aporops bilinearis|MARQ276-12|MARQ-277|MK566801
Pseudogramma paucilepis|AUSTR497-13|AUST-499|KU905727
Pseudogramma paucilepis|AUSTR245-13|AUST-247|KU905725
Pseudogramma paucilepis|GAMBA150-12|GAM-011|KU905722
Pseudogramma paucilepis|AUSTR498-13|AUST-500|KU905726
Pseudogramma paucilepis|AUSTR495-13|AUST-497|KU905723
Pseudogramma paucilepis|AUSTR243-13|AUST-245|KU905724
Pseudogramma polyacantha|SCILL131-15|SCIL-131|MK658392
Pseudogramma polyacantha|SCILL230-15|SCIL-230|MK658338
Pseudogramma polyacantha|SCILL009-15|SCIL-009|MK658212
Pseudogramma galzini|AUSTR242-13|AUST-244|KU905711
Pseudogramma galzini|AUSTR241-13|AUST-243|KU905712
Pseudogramma galzini|AUSTR558-13|AUST-560|KU905719
Pseudogramma galzini|GAMBA151-12|GAM-012|KU905721
Pseudogramma galzini|GAMBA869-12|GAM-744|KU905716
Pseudogramma galzini|AUSTR559-13|AUST-561|KU905718
Pseudogramma galzini|AUSTR244-13|AUST-246|KU905710
Pseudogramma galzini|GAMBA149-12|GAM-010|KU905717
Pseudogramma galzini|AUSTR360-13|AUST-362|KU905720
Pseudogramma galzini|AUSTR044-13|AUST-044|KU905713
Pseudogramma galzini|AUSTR042-13|AUST-042|KU905715
Pseudogramma galzini|AUSTR043-13|AUST-043|KU905714
Pseudogramma galzini|AUSTR041-13|AUST-041|KU905709
Pseudogramma brederi|MARQ415-12|MARQ-417|MK567074
Pseudogramma brederi|MARQ417-12|MARQ-419|MK567073
Pseudogramma brederi|MOH199-16|MOH-199|MK658335
Pseudogramma brederi|AUSTR496-13|AUST-498|MK658680
Pseudogramma brederi|MARQ147-12|MARQ-147|MK567072
Pseudogramma brederi|MOH198-16|MOH-198|MK658031
Pseudogramma brederi|MARQ416-12|MARQ-418|MK567071
Pseudogramma brederi|MARQ146-12|MARQ-146|MK567070
Fowleria isostigma|SCILL071-15|SCIL-071|MK657601
Fowleria variegata|GAMBA379-12|GAM-244|MK658295
Fowleria variegata|GAMBA378-12|GAM-243|MK657735
Fowleria variegata|GAMBA377-12|GAM-242|MK657465
Fowleria marmorata|SCILL070-15|SCIL-070|MK657290
Fowleria marmorata|SCILL325-15|SCIL-325|MK658581
Fowleria marmorata|SCILL006-15|SCIL-006|MK657425
Fowleria|MARQ107-12|MARQ-107|MK566920
Fowleria|MARQ106-12|MARQ-106|MK566918
Fowleria|MARQ105-12|MARQ-105|MK566919
Fowleria vaiulae|GAMBA598-12|GAM-467|MK658594
Fowleria vaiulae|GAMBA466-12|GAM-333|MK658482
Fowleria vaiulae|GAMBA075-12|GAM-844|MK658387
Fowleria vaiulae|SCILL249-15|SCIL-249|MK658356
Fowleria vaiulae|GAMBA596-12|GAM-465|MK658598
Fowleria vaiulae|GAMBA467-12|GAM-334|MK657142
Fowleria vaiulae|GAMBA465-12|GAM-332|MK657560
Fowleria vaiulae|GAMBA597-12|GAM-466|MK657529
Fowleria vaiulae|AUSTR005-13|AUST-005|MK657026
Apogonichthys perdix|GAMBA050-12|GAM-818|MK657620
Apogonichthys perdix|AUSTR119-13|AUST-119|MK657931
Apogonichthys perdix|MARQ494-12|MARQ-496|MK566796
Apogonichthys perdix|MARQ401-12|MARQ-402|MK566797
Apogonichthys ocellatus|MARQ397-12|MARQ-398|MK566793
Apogonichthys ocellatus|MARQ350-12|MARQ-351|MK566795
Apogonichthys ocellatus|MARQ349-12|MARQ-350|MK566794
Apogon deetsie|SCILL128-15|SCIL-128|MK658081
Zapogon evermanni|MARQ035-12|MARQ-035|MK567152
Cheilodipterus macrodon|GAMBA078-12|GAM-847|MK657972
Cheilodipterus macrodon|AUSTR135-13|AUST-135|MK657654
Cheilodipterus artus|GAMBA010-12|GAM-778|MK658712
Cheilodipterus artus|SCILL332-15|SCIL-332|MK657287
Cheilodipterus artus|SCILL067-15|SCIL-067|MK658117
Cheilodipterus artus|GAMBA011-12|GAM-779|MK658107
Cheilodipterus artus|GAMBA009-12|GAM-777|MK658515
Cheilodipterus artus|AUSTR134-13|AUST-134|MK657401
Cheilodipterus quinquelineatus|GAMBA446-12|GAM-313|MK658541
Cheilodipterus quinquelineatus|SCILL265-15|SCIL-265|MK657488
Cheilodipterus quinquelineatus|GAMBA447-12|GAM-314|MK657893
Cheilodipterus quinquelineatus|AUSTR008-13|AUST-008|MK657498
Cheilodipterus quinquelineatus|AUSTR007-13|AUST-007|MK657613
Ostorhinchus novemfasciatus|SCILL247-15|SCIL-247|MK657715
Ostorhinchus nigrofasciatus|SCILL231-15|SCIL-231|MK657132
Ostorhinchus nigrofasciatus|SCILL229-15|SCIL-229|MK657953
Ostorhinchus nigrofasciatus|GAMBA054-12|GAM-822|MK658291
Ostorhinchus nigrofasciatus|GAMBA053-12|GAM-821|MK657029
Ostorhinchus nigrofasciatus|AUSTR137-13|AUST-137|MK657821
Ostorhinchus nigrofasciatus|AUSTR136-13|AUST-136|MK657670
Ostorhinchus angustatus|SCILL142-15|SCIL-142|MK658310
Ostorhinchus relativus|MOH031-16|MOH-031|MK658619
Ostorhinchus taeniophorus|GAMBA252-12|GAM-115|MK657449
Ostorhinchus taeniophorus|GAMBA055-12|GAM-823|MK657151
Ostorhinchus taeniophorus|AUSTR362-13|AUST-364|MK658015
Ostorhinchus taeniophorus|AUSTR361-13|AUST-363|MK657870
Ostorhinchus taeniophorus|AUSTR204-13|AUST-206|MK658498
Ostorhinchus sinus|MARQ342-12|MARQ-343|MK567008
Ostorhinchus sinus|MARQ341-12|MARQ-342|MK567007
Ostorhinchus sinus|MARQ340-12|MARQ-341|MK567005
Ostorhinchus sinus|MARQ339-12|MARQ-340|MK567006
Ostorhinchus apogonoides|MARQ036-12|MARQ-036|MK567002
Ostorhinchus apogonoides|MARQ037-12|MARQ-037|MK567004
Ostorhinchus apogonoides|MOH090-16|MOH-090|MK657572
Ostorhinchus apogonoides|MOH091-16|MOH-091|MK657361
Ostorhinchus apogonoides|MOH164-16|MOH-164|MK658269
Ostorhinchus apogonoides|MARQ182-12|MARQ-183|MK567003
Ostorhinchus apogonoides|MARQ181-12|MARQ-182|MK567001
Pristiapogon fraenatus|GAMBA450-12|GAM-317|MK657818
Pristiapogon fraenatus|GAMBA449-12|GAM-316|MK657305
Pristiapogon fraenatus|GAMBA448-12|GAM-315|MK657998
Pristiapogon exostigma|GAMBA421-12|GAM-286|MK656972

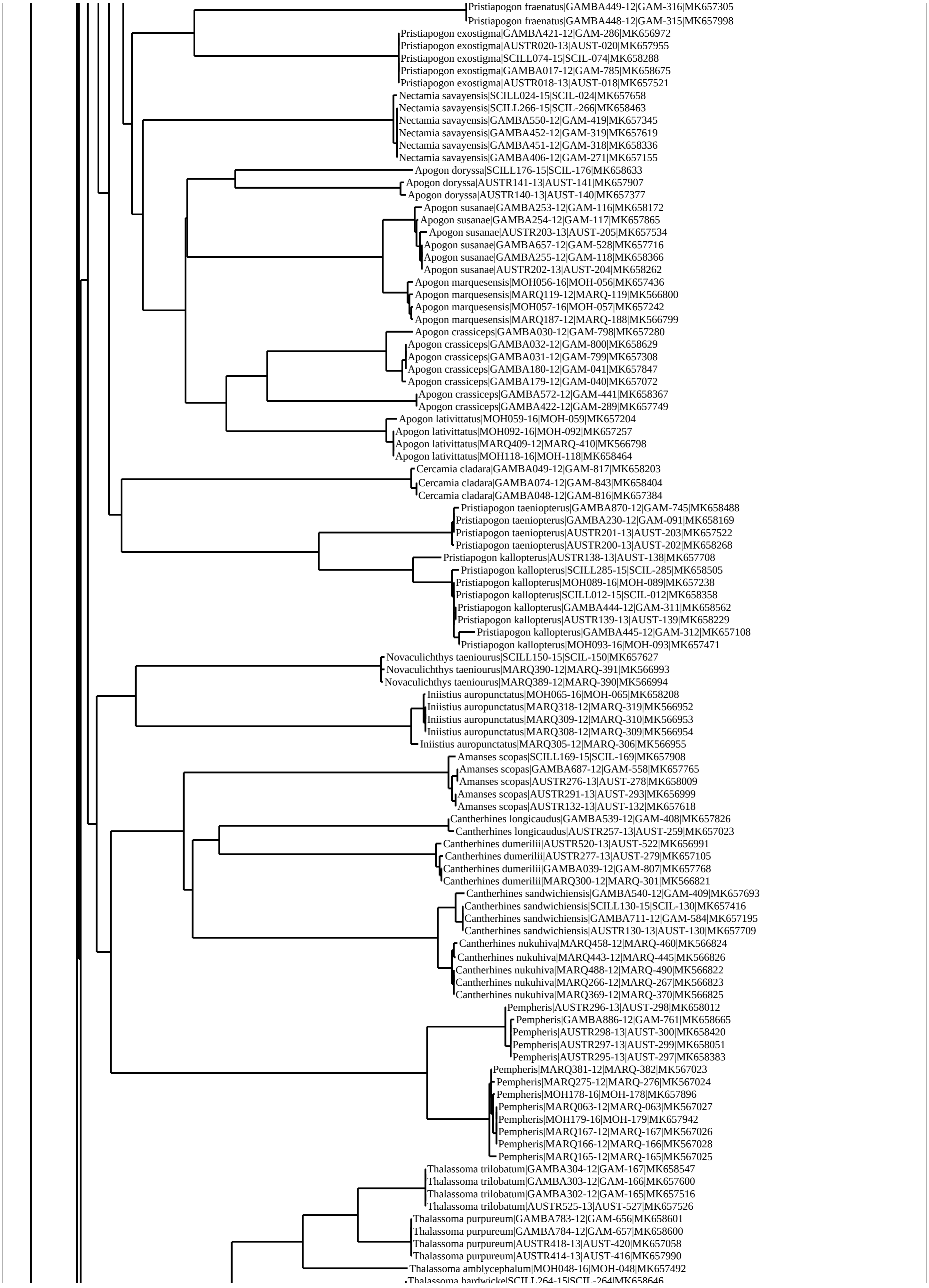
Pristiapogon fraenatus|GAMBA449-12|GAM-316|MK657305
Pristiapogon fraenatus|GAMBA448-12|GAM-315|MK657998
Pristiapogon exostigma|GAMBA421-12|GAM-286|MK656972
Pristiapogon exostigma|AUSTR020-13|AUST-020|MK657955
Pristiapogon exostigma|SCILL074-15|SCIL-074|MK658288
Pristiapogon exostigma|GAMBA017-12|GAM-785|MK658675
Pristiapogon exostigma|AUSTR018-13|AUST-018|MK657521
Nectamia savayensis|SCILL024-15|SCIL-024|MK657658
Nectamia savayensis|SCILL266-15|SCIL-266|MK658463
Nectamia savayensis|GAMBA550-12|GAM-419|MK657345
Nectamia savayensis|GAMBA452-12|GAM-319|MK657619
Nectamia savayensis|GAMBA451-12|GAM-318|MK658336
Nectamia savayensis|GAMBA406-12|GAM-271|MK657155
Apogon doryssa|SCILL176-15|SCIL-176|MK658633
Apogon doryssa|AUSTR141-13|AUST-141|MK657907
Apogon doryssa|AUSTR140-13|AUST-140|MK657377
Apogon susanae|GAMBA253-12|GAM-116|MK658172
Apogon susanae|GAMBA254-12|GAM-117|MK657865
Apogon susanae|AUSTR203-13|AUST-205|MK657534
Apogon susanae|GAMBA657-12|GAM-528|MK657716
Apogon susanae|GAMBA255-12|GAM-118|MK658366
Apogon susanae|AUSTR202-13|AUST-204|MK658262
Apogon marquesensis|MOH056-16|MOH-056|MK657436
Apogon marquesensis|MARQ119-12|MARQ-119|MK566800
Apogon marquesensis|MOH057-16|MOH-057|MK657242
Apogon marquesensis|MARQ187-12|MARQ-188|MK566799
Apogon crassiceps|GAMBA030-12|GAM-798|MK657280
Apogon crassiceps|GAMBA032-12|GAM-800|MK658629
Apogon crassiceps|GAMBA031-12|GAM-799|MK657308
Apogon crassiceps|GAMBA180-12|GAM-041|MK657847
Apogon crassiceps|GAMBA179-12|GAM-040|MK657072
Apogon crassiceps|GAMBA572-12|GAM-441|MK658367
Apogon crassiceps|GAMBA422-12|GAM-289|MK657749
Apogon lativittatus|MOH059-16|MOH-059|MK657204
Apogon lativittatus|MOH092-16|MOH-092|MK657257
Apogon lativittatus|MARQ409-12|MARQ-410|MK566798
Apogon lativittatus|MOH118-16|MOH-118|MK658464
Cercamia cladara|GAMBA049-12|GAM-817|MK658203
Cercamia cladara|GAMBA074-12|GAM-843|MK658404
Cercamia cladara|GAMBA048-12|GAM-816|MK657384
Pristiapogon taeniopterus|GAMBA870-12|GAM-745|MK658488
Pristiapogon taeniopterus|GAMBA230-12|GAM-091|MK658169
Pristiapogon taeniopterus|AUSTR201-13|AUST-203|MK657522
Pristiapogon taeniopterus|AUSTR200-13|AUST-202|MK658268
Pristiapogon kallopterus|AUSTR138-13|AUST-138|MK657708
Pristiapogon kallopterus|SCILL285-15|SCIL-285|MK658505
Pristiapogon kallopterus|MOH089-16|MOH-089|MK657238
Pristiapogon kallopterus|SCILL012-15|SCIL-012|MK658358
Pristiapogon kallopterus|GAMBA444-12|GAM-311|MK658562
Pristiapogon kallopterus|AUSTR139-13|AUST-139|MK658229
Pristiapogon kallopterus|GAMBA445-12|GAM-312|MK657108
Pristiapogon kallopterus|MOH093-16|MOH-093|MK657471
Novaculichthys taeniourus|SCILL150-15|SCIL-150|MK657627
Novaculichthys taeniourus|MARQ390-12|MARQ-391|MK566993
Novaculichthys taeniourus|MARQ389-12|MARQ-390|MK566994
Iniistius auropunctatus|MOH065-16|MOH-065|MK658208
Iniistius auropunctatus|MARQ318-12|MARQ-319|MK566952
Iniistius auropunctatus|MARQ309-12|MARQ-310|MK566953
Iniistius auropunctatus|MARQ308-12|MARQ-309|MK566954
Iniistius auropunctatus|MARQ305-12|MARQ-306|MK566955
Amanses scopas|SCILL169-15|SCIL-169|MK657908
Amanses scopas|GAMBA687-12|GAM-558|MK657765
Amanses scopas|AUSTR276-13|AUST-278|MK658009
Amanses scopas|AUSTR291-13|AUST-293|MK656999
Amanses scopas|AUSTR132-13|AUST-132|MK657618
Cantherhines longicaudus|GAMBA539-12|GAM-408|MK657826
Cantherhines longicaudus|AUSTR257-13|AUST-259|MK657023
Cantherhines dumerilii|AUSTR520-13|AUST-522|MK656991
Cantherhines dumerilii|AUSTR277-13|AUST-279|MK657105
Cantherhines dumerilii|GAMBA039-12|GAM-807|MK657768
Cantherhines dumerilii|MARQ300-12|MARQ-301|MK566821
Cantherhines sandwichiensis|GAMBA540-12|GAM-409|MK657693
Cantherhines sandwichiensis|SCILL130-15|SCIL-130|MK657416
Cantherhines sandwichiensis|GAMBA711-12|GAM-584|MK657195
Cantherhines sandwichiensis|AUSTR130-13|AUST-130|MK657709
Cantherhines nukuhiva|MARQ458-12|MARQ-460|MK566824
Cantherhines nukuhiva|MARQ443-12|MARQ-445|MK566826
Cantherhines nukuhiva|MARQ488-12|MARQ-490|MK566822
Cantherhines nukuhiva|MARQ266-12|MARQ-267|MK566823
Cantherhines nukuhiva|MARQ369-12|MARQ-370|MK566825
Pempheris|AUSTR296-13|AUST-298|MK658012
Pempheris|GAMBA886-12|GAM-761|MK658665
Pempheris|AUSTR298-13|AUST-300|MK658420
Pempheris|AUSTR297-13|AUST-299|MK658051
Pempheris|AUSTR295-13|AUST-297|MK658383
Pempheris|MARQ381-12|MARQ-382|MK567023
Pempheris|MARQ275-12|MARQ-276|MK567024
Pempheris|MOH178-16|MOH-178|MK657896
Pempheris|MARQ063-12|MARQ-063|MK567027
Pempheris|MOH179-16|MOH-179|MK657942
Pempheris|MARQ167-12|MARQ-167|MK567026
Pempheris|MARQ166-12|MARQ-166|MK567028
Pempheris|MARQ165-12|MARQ-165|MK567025
Thalassoma trilobatum|GAMBA304-12|GAM-167|MK658547
Thalassoma trilobatum|GAMBA303-12|GAM-166|MK657600
Thalassoma trilobatum|GAMBA302-12|GAM-165|MK657516
Thalassoma trilobatum|AUSTR525-13|AUST-527|MK657526
Thalassoma purpureum|GAMBA783-12|GAM-656|MK658601
Thalassoma purpureum|GAMBA784-12|GAM-657|MK658600
Thalassoma purpureum|AUSTR418-13|AUST-420|MK657058
Thalassoma purpureum|AUSTR414-13|AUST-416|MK657990
Thalassoma amblycephalum|MOH048-16|MOH-048|MK657492
Thalassoma hardwicke|SCILL264-15|SCIL-264|MK658646

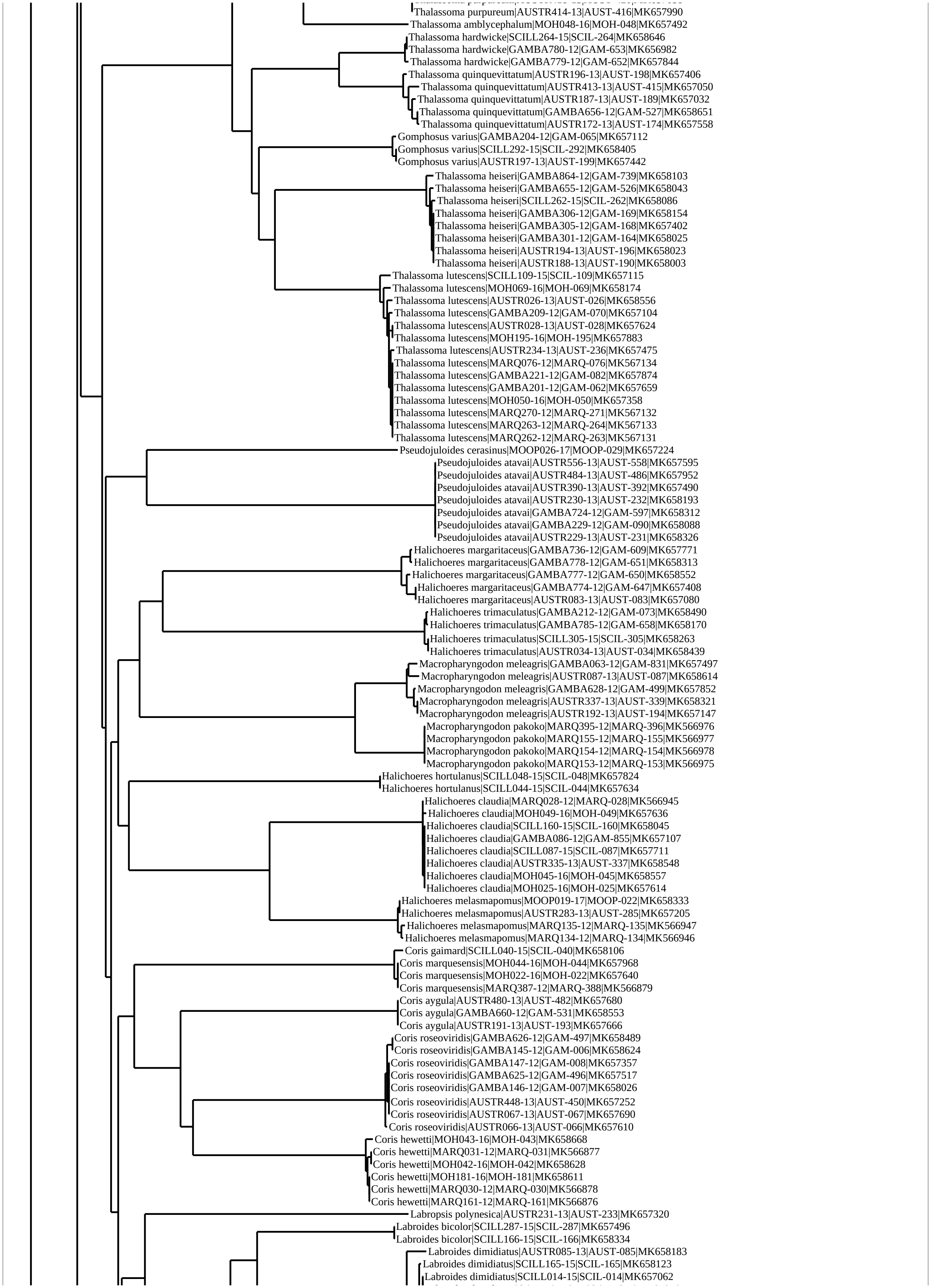
Thalassoma purpureum|AUSTR414-13|AUST-416|MK657990
Thalassoma amblycephalum|MOH048-16|MOH-048|MK657492
Thalassoma hardwicke|SCILL264-15|SCIL-264|MK658646
Thalassoma hardwicke|GAMBA780-12|GAM-653|MK656982
Thalassoma hardwicke|GAMBA779-12|GAM-652|MK657844
Thalassoma quinquevittatum|AUSTR196-13|AUST-198|MK657406
Thalassoma quinquevittatum|AUSTR413-13|AUST-415|MK657050
Thalassoma quinquevittatum|AUSTR187-13|AUST-189|MK657032
Thalassoma quinquevittatum|GAMBA656-12|GAM-527|MK658651
Thalassoma quinquevittatum|AUSTR172-13|AUST-174|MK657558
Gomphosus varius|GAMBA204-12|GAM-065|MK657112
Gomphosus varius|SCILL292-15|SCIL-292|MK658405
Gomphosus varius|AUSTR197-13|AUST-199|MK657442
Thalassoma heiseri|GAMBA864-12|GAM-739|MK658103
Thalassoma heiseri|GAMBA655-12|GAM-526|MK658043
Thalassoma heiseri|SCILL262-15|SCIL-262|MK658086
Thalassoma heiseri|GAMBA306-12|GAM-169|MK658154
Thalassoma heiseri|GAMBA305-12|GAM-168|MK657402
Thalassoma heiseri|GAMBA301-12|GAM-164|MK658025
Thalassoma heiseri|AUSTR194-13|AUST-196|MK658023
Thalassoma heiseri|AUSTR188-13|AUST-190|MK658003
Thalassoma lutescens|SCILL109-15|SCIL-109|MK657115
Thalassoma lutescens|MOH069-16|MOH-069|MK658174
Thalassoma lutescens|AUSTR026-13|AUST-026|MK658556
Thalassoma lutescens|GAMBA209-12|GAM-070|MK657104
Thalassoma lutescens|AUSTR028-13|AUST-028|MK657624
Thalassoma lutescens|MOH195-16|MOH-195|MK657883
Thalassoma lutescens|AUSTR234-13|AUST-236|MK657475
Thalassoma lutescens|MARQ076-12|MARQ-076|MK567134
Thalassoma lutescens|GAMBA221-12|GAM-082|MK657874
Thalassoma lutescens|GAMBA201-12|GAM-062|MK657659
Thalassoma lutescens|MOH050-16|MOH-050|MK657358
Thalassoma lutescens|MARQ270-12|MARQ-271|MK567132
Thalassoma lutescens|MARQ263-12|MARQ-264|MK567133
Thalassoma lutescens|MARQ262-12|MARQ-263|MK567131
Pseudojuloides cerasinus|MOOP026-17|MOOP-029|MK657224
Pseudojuloides atavai|AUSTR556-13|AUST-558|MK657595
Pseudojuloides atavai|AUSTR484-13|AUST-486|MK657952
Pseudojuloides atavai|AUSTR390-13|AUST-392|MK657490
Pseudojuloides atavai|AUSTR230-13|AUST-232|MK658193
Pseudojuloides atavai|GAMBA724-12|GAM-597|MK658312
Pseudojuloides atavai|GAMBA229-12|GAM-090|MK658088
Pseudojuloides atavai|AUSTR229-13|AUST-231|MK658326
Halichoeres margaritaceus|GAMBA736-12|GAM-609|MK657771
Halichoeres margaritaceus|GAMBA778-12|GAM-651|MK658313
Halichoeres margaritaceus|GAMBA777-12|GAM-650|MK658552
Halichoeres margaritaceus|GAMBA774-12|GAM-647|MK657408
Halichoeres margaritaceus|AUSTR083-13|AUST-083|MK657080
Halichoeres trimaculatus|GAMBA212-12|GAM-073|MK658490
Halichoeres trimaculatus|GAMBA785-12|GAM-658|MK658170
Halichoeres trimaculatus|SCILL305-15|SCIL-305|MK658263
Halichoeres trimaculatus|AUSTR034-13|AUST-034|MK658439
Macropharyngodon meleagris|GAMBA063-12|GAM-831|MK657497
Macropharyngodon meleagris|AUSTR087-13|AUST-087|MK658614
Macropharyngodon meleagris|GAMBA628-12|GAM-499|MK657852
Macropharyngodon meleagris|AUSTR337-13|AUST-339|MK658321
Macropharyngodon meleagris|AUSTR192-13|AUST-194|MK657147
Macropharyngodon pakoko|MARQ395-12|MARQ-396|MK566976
Macropharyngodon pakoko|MARQ155-12|MARQ-155|MK566977
Macropharyngodon pakoko|MARQ154-12|MARQ-154|MK566978
Macropharyngodon pakoko|MARQ153-12|MARQ-153|MK566975
Halichoeres hortulanus|SCILL048-15|SCIL-048|MK657824
Halichoeres hortulanus|SCILL044-15|SCIL-044|MK657634
Halichoeres claudia|MARQ028-12|MARQ-028|MK566945
Halichoeres claudia|MOH049-16|MOH-049|MK657636
Halichoeres claudia|SCILL160-15|SCIL-160|MK658045
Halichoeres claudia|GAMBA086-12|GAM-855|MK657107
Halichoeres claudia|SCILL087-15|SCIL-087|MK657711
Halichoeres claudia|AUSTR335-13|AUST-337|MK658548
Halichoeres claudia|MOH045-16|MOH-045|MK658557
Halichoeres claudia|MOH025-16|MOH-025|MK657614
Halichoeres melasmapomus|MOOP019-17|MOOP-022|MK658333
Halichoeres melasmapomus|AUSTR283-13|AUST-285|MK657205
Halichoeres melasmapomus|MARQ135-12|MARQ-135|MK566947
Halichoeres melasmapomus|MARQ134-12|MARQ-134|MK566946
Coris gaimard|SCILL040-15|SCIL-040|MK658106
Coris marquesensis|MOH044-16|MOH-044|MK657968
Coris marquesensis|MOH022-16|MOH-022|MK657640
Coris marquesensis|MARQ387-12|MARQ-388|MK566879
Coris aygula|AUSTR480-13|AUST-482|MK657680
Coris aygula|GAMBA660-12|GAM-531|MK658553
Coris aygula|AUSTR191-13|AUST-193|MK657666
Coris roseoviridis|GAMBA626-12|GAM-497|MK658489
Coris roseoviridis|GAMBA145-12|GAM-006|MK658624
Coris roseoviridis|GAMBA147-12|GAM-008|MK657357
Coris roseoviridis|GAMBA625-12|GAM-496|MK657517
Coris roseoviridis|GAMBA146-12|GAM-007|MK658026
Coris roseoviridis|AUSTR448-13|AUST-450|MK657252
Coris roseoviridis|AUSTR067-13|AUST-067|MK657690
Coris roseoviridis|AUSTR066-13|AUST-066|MK657610
Coris hewetti|MOH043-16|MOH-043|MK658668
Coris hewetti|MARQ031-12|MARQ-031|MK566877
Coris hewetti|MOH042-16|MOH-042|MK658628
Coris hewetti|MOH181-16|MOH-181|MK658611
Coris hewetti|MARQ030-12|MARQ-030|MK566878
Coris hewetti|MARQ161-12|MARQ-161|MK566876
Labropsis polynesica|AUSTR231-13|AUST-233|MK657320
Labroides bicolor|SCILL287-15|SCIL-287|MK657496
Labroides bicolor|SCILL166-15|SCIL-166|MK658334
Labroides dimidiatus|AUSTR085-13|AUST-085|MK658183
Labroides dimidiatus|SCILL165-15|SCIL-165|MK658123
Labroides dimidiatus|SCILL014-15|SCIL-014|MK657062

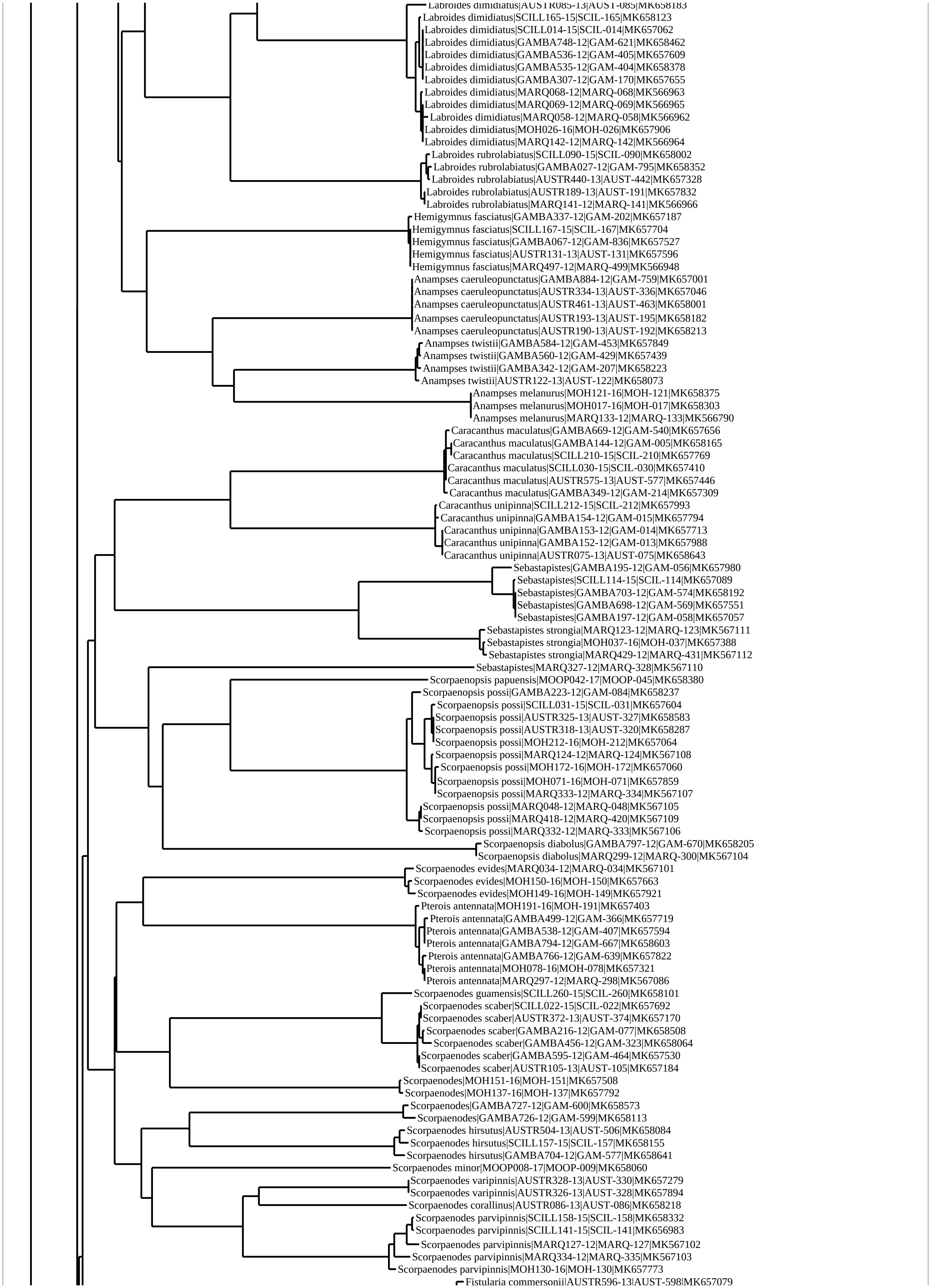
Labroides dimidiatus|AUSTR085-13|AUST-085|MK658183
Labroides dimidiatus|SCILL165-15|SCIL-165|MK658123
Labroides dimidiatus|SCILL014-15|SCIL-014|MK657062
Labroides dimidiatus|GAMBA748-12|GAM-621|MK658462
Labroides dimidiatus|GAMBA536-12|GAM-405|MK657609
Labroides dimidiatus|GAMBA535-12|GAM-404|MK658378
Labroides dimidiatus|GAMBA307-12|GAM-170|MK657655
Labroides dimidiatus|MARQ068-12|MARQ-068|MK566963
Labroides dimidiatus|MARQ069-12|MARQ-069|MK566965
Labroides dimidiatus|MARQ058-12|MARQ-058|MK566962
Labroides dimidiatus|MOH026-16|MOH-026|MK657906
Labroides dimidiatus|MARQ142-12|MARQ-142|MK566964
Labroides rubrolabiatus|SCILL090-15|SCIL-090|MK658002
Labroides rubrolabiatus|GAMBA027-12|GAM-795|MK658352
Labroides rubrolabiatus|AUSTR440-13|AUST-442|MK657328
Labroides rubrolabiatus|AUSTR189-13|AUST-191|MK657832
Labroides rubrolabiatus|MARQ141-12|MARQ-141|MK566966
Hemigymnus fasciatus|GAMBA337-12|GAM-202|MK657187
Hemigymnus fasciatus|SCILL167-15|SCIL-167|MK657704
Hemigymnus fasciatus|GAMBA067-12|GAM-836|MK657527
Hemigymnus fasciatus|AUSTR131-13|AUST-131|MK657596
Hemigymnus fasciatus|MARQ497-12|MARQ-499|MK566948
Anampses caeruleopunctatus|GAMBA884-12|GAM-759|MK657001
Anampses caeruleopunctatus|AUSTR334-13|AUST-336|MK657046
Anampses caeruleopunctatus|AUSTR461-13|AUST-463|MK658001
Anampses caeruleopunctatus|AUSTR193-13|AUST-195|MK658182
Anampses caeruleopunctatus|AUSTR190-13|AUST-192|MK658213
Anampses twistii|GAMBA584-12|GAM-453|MK657849
Anampses twistii|GAMBA560-12|GAM-429|MK657439
Anampses twistii|GAMBA342-12|GAM-207|MK658223
Anampses twistii|AUSTR122-13|AUST-122|MK658073
Anampses melanurus|MOH121-16|MOH-121|MK658375
Anampses melanurus|MOH017-16|MOH-017|MK658303
Anampses melanurus|MARQ133-12|MARQ-133|MK566790
Caracanthus maculatus|GAMBA669-12|GAM-540|MK657656
Caracanthus maculatus|GAMBA144-12|GAM-005|MK658165
Caracanthus maculatus|SCILL210-15|SCIL-210|MK657769
Caracanthus maculatus|SCILL030-15|SCIL-030|MK657410
Caracanthus maculatus|AUSTR575-13|AUST-577|MK657446
Caracanthus maculatus|GAMBA349-12|GAM-214|MK657309
Caracanthus unipinna|SCILL212-15|SCIL-212|MK657993
Caracanthus unipinna|GAMBA154-12|GAM-015|MK657794
Caracanthus unipinna|GAMBA153-12|GAM-014|MK657713
Caracanthus unipinna|GAMBA152-12|GAM-013|MK657988
Caracanthus unipinna|AUSTR075-13|AUST-075|MK658643
Sebastapistes|GAMBA195-12|GAM-056|MK657980
Sebastapistes|SCILL114-15|SCIL-114|MK657089
Sebastapistes|GAMBA703-12|GAM-574|MK658192
Sebastapistes|GAMBA698-12|GAM-569|MK657551
Sebastapistes|GAMBA197-12|GAM-058|MK657057
Sebastapistes strongia|MARQ123-12|MARQ-123|MK567111
Sebastapistes strongia|MOH037-16|MOH-037|MK657388
Sebastapistes strongia|MARQ429-12|MARQ-431|MK567112
Sebastapistes|MARQ327-12|MARQ-328|MK567110
Scorpaenopsis papuensis|MOOP042-17|MOOP-045|MK658380
Scorpaenopsis possi|GAMBA223-12|GAM-084|MK658237
Scorpaenopsis possi|SCILL031-15|SCIL-031|MK657604
Scorpaenopsis possi|AUSTR325-13|AUST-327|MK658583
Scorpaenopsis possi|AUSTR318-13|AUST-320|MK658287
Scorpaenopsis possi|MOH212-16|MOH-212|MK657064
Scorpaenopsis possi|MARQ124-12|MARQ-124|MK567108
Scorpaenopsis possi|MOH172-16|MOH-172|MK657060
Scorpaenopsis possi|MOH071-16|MOH-071|MK657859
Scorpaenopsis possi|MARQ333-12|MARQ-334|MK567107
Scorpaenopsis possi|MARQ048-12|MARQ-048|MK567105
Scorpaenopsis possi|MARQ418-12|MARQ-420|MK567109
Scorpaenopsis possi|MARQ332-12|MARQ-333|MK567106
Scorpaenopsis diabolus|GAMBA797-12|GAM-670|MK658205
Scorpaenopsis diabolus|MARQ299-12|MARQ-300|MK567104
Scorpaenodes evides|MARQ034-12|MARQ-034|MK567101
Scorpaenodes evides|MOH150-16|MOH-150|MK657663
Scorpaenodes evides|MOH149-16|MOH-149|MK657921
Pterois antennata|MOH191-16|MOH-191|MK657403
Pterois antennata|GAMBA499-12|GAM-366|MK657719
Pterois antennata|GAMBA538-12|GAM-407|MK657594
Pterois antennata|GAMBA794-12|GAM-667|MK658603
Pterois antennata|GAMBA766-12|GAM-639|MK657822
Pterois antennata|MOH078-16|MOH-078|MK657321
Pterois antennata|MARQ297-12|MARQ-298|MK567086
Scorpaenodes guamensis|SCILL260-15|SCIL-260|MK658101
Scorpaenodes scaber|SCILL022-15|SCIL-022|MK657692
Scorpaenodes scaber|AUSTR372-13|AUST-374|MK657170
Scorpaenodes scaber|GAMBA216-12|GAM-077|MK658508
Scorpaenodes scaber|GAMBA456-12|GAM-323|MK658064
Scorpaenodes scaber|GAMBA595-12|GAM-464|MK657530
Scorpaenodes scaber|AUSTR105-13|AUST-105|MK657184
Scorpaenodes|MOH151-16|MOH-151|MK657508
Scorpaenodes|MOH137-16|MOH-137|MK657792
Scorpaenodes|GAMBA727-12|GAM-600|MK658573
Scorpaenodes|GAMBA726-12|GAM-599|MK658113
Scorpaenodes hirsutus|AUSTR504-13|AUST-506|MK658084
Scorpaenodes hirsutus|SCILL157-15|SCIL-157|MK658155
Scorpaenodes hirsutus|GAMBA704-12|GAM-577|MK658641
Scorpaenodes minor|MOOP008-17|MOOP-009|MK658060
Scorpaenodes varipinnis|AUSTR328-13|AUST-330|MK657279
Scorpaenodes varipinnis|AUSTR326-13|AUST-328|MK657894
Scorpaenodes corallinus|AUSTR086-13|AUST-086|MK658218
Scorpaenodes parvipinnis|SCILL158-15|SCIL-158|MK658332
Scorpaenodes parvipinnis|SCILL141-15|SCIL-141|MK656983
Scorpaenodes parvipinnis|MARQ127-12|MARQ-127|MK567102
Scorpaenodes parvipinnis|MARQ334-12|MARQ-335|MK567103
Scorpaenodes parvipinnis|MOH130-16|MOH-130|MK657773
Fistularia commersonii|AUSTR596-13|AUST-598|MK657079

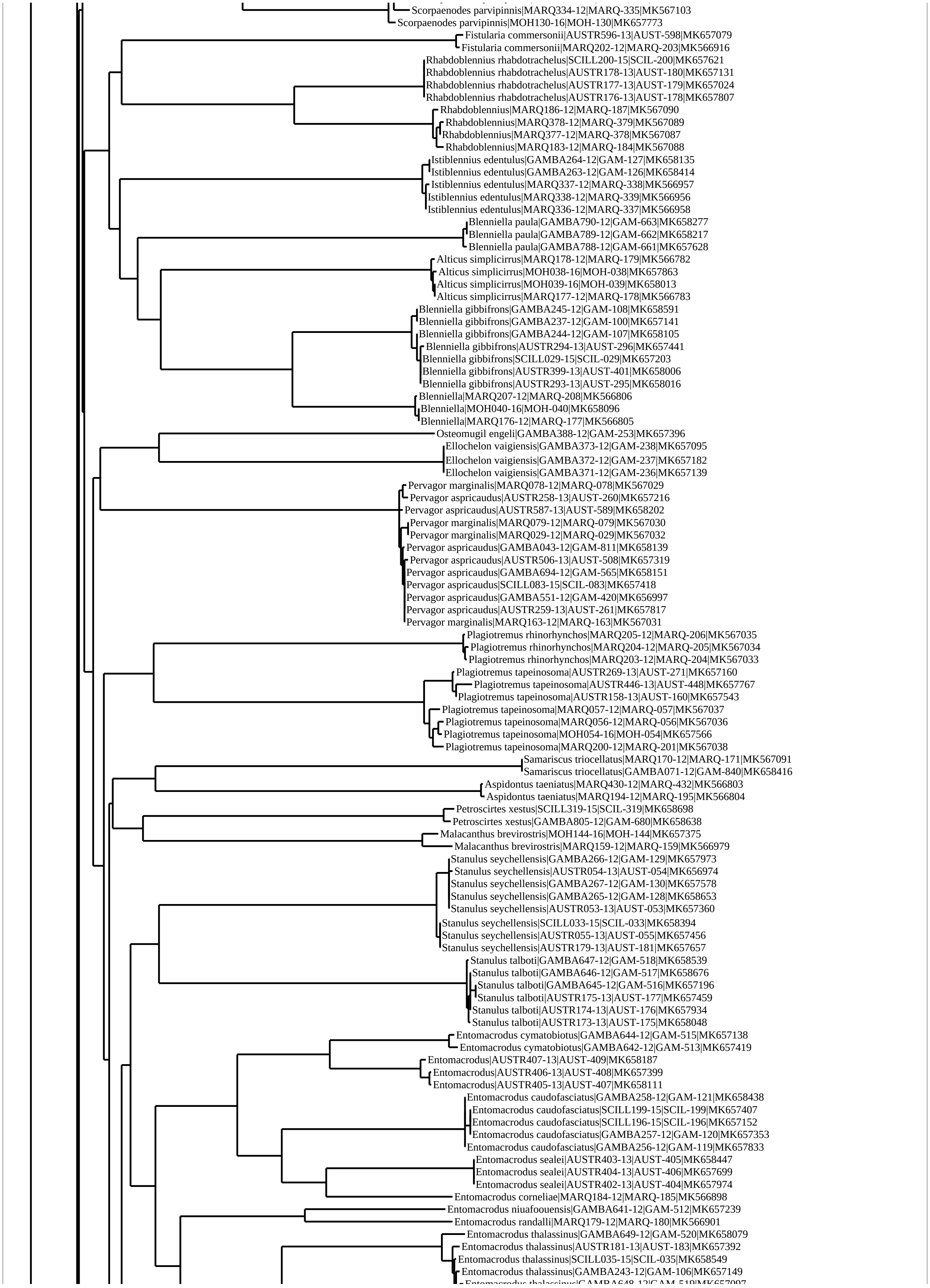
Scorpaenodes parvipinnis|MARQ334-12|MARQ-335|MK567103
Scorpaenodes parvipinnis|MOH130-16|MOH-130|MK657773
Fistularia commersonii|AUSTR596-13|AUST-598|MK657079
Fistularia commersonii|MARQ202-12|MARQ-203|MK566916
Rhabdoblennius rhabdotrachelus|SCILL200-15|SCIL-200|MK657621
Rhabdoblennius rhabdotrachelus|AUSTR178-13|AUST-180|MK657131
Rhabdoblennius rhabdotrachelus|AUSTR177-13|AUST-179|MK657024
Rhabdoblennius rhabdotrachelus|AUSTR176-13|AUST-178|MK657807
Rhabdoblennius|MARQ186-12|MARQ-187|MK567090
Rhabdoblennius|MARQ378-12|MARQ-379|MK567089
Rhabdoblennius|MARQ377-12|MARQ-378|MK567087
Rhabdoblennius|MARQ183-12|MARQ-184|MK567088
Istiblennius edentulus|GAMBA264-12|GAM-127|MK658135
Istiblennius edentulus|GAMBA263-12|GAM-126|MK658414
Istiblennius edentulus|MARQ337-12|MARQ-338|MK566957
Istiblennius edentulus|MARQ338-12|MARQ-339|MK566956
Istiblennius edentulus|MARQ336-12|MARQ-337|MK566958
Blenniella paula|GAMBA790-12|GAM-663|MK658277
Blenniella paula|GAMBA789-12|GAM-662|MK658217
Blenniella paula|GAMBA788-12|GAM-661|MK657628
Alticus simplicirrus|MARQ178-12|MARQ-179|MK566782
Alticus simplicirrus|MOH038-16|MOH-038|MK657863
Alticus simplicirrus|MOH039-16|MOH-039|MK658013
Alticus simplicirrus|MARQ177-12|MARQ-178|MK566783
Blenniella gibbifrons|GAMBA245-12|GAM-108|MK658591
Blenniella gibbifrons|GAMBA237-12|GAM-100|MK657141
Blenniella gibbifrons|GAMBA244-12|GAM-107|MK658105
Blenniella gibbifrons|AUSTR294-13|AUST-296|MK657441
Blenniella gibbifrons|SCILL029-15|SCIL-029|MK657203
Blenniella gibbifrons|AUSTR399-13|AUST-401|MK658006
Blenniella gibbifrons|AUSTR293-13|AUST-295|MK658016
Blenniella|MARQ207-12|MARQ-208|MK566806
Blenniella|MOH040-16|MOH-040|MK658096
Blenniella|MARQ176-12|MARQ-177|MK566805
Osteomugil engeli|GAMBA388-12|GAM-253|MK657396
Ellochelon vaigiensis|GAMBA373-12|GAM-238|MK657095
Ellochelon vaigiensis|GAMBA372-12|GAM-237|MK657182
Ellochelon vaigiensis|GAMBA371-12|GAM-236|MK657139
Pervagor marginalis|MARQ078-12|MARQ-078|MK567029
Pervagor aspricaudus|AUSTR258-13|AUST-260|MK657216
Pervagor aspricaudus|AUSTR587-13|AUST-589|MK658202
Pervagor marginalis|MARQ079-12|MARQ-079|MK567030
Pervagor marginalis|MARQ029-12|MARQ-029|MK567032
Pervagor aspricaudus|GAMBA043-12|GAM-811|MK658139
Pervagor aspricaudus|AUSTR506-13|AUST-508|MK657319
Pervagor aspricaudus|GAMBA694-12|GAM-565|MK658151
Pervagor aspricaudus|SCILL083-15|SCIL-083|MK657418
Pervagor aspricaudus|GAMBA551-12|GAM-420|MK656997
Pervagor aspricaudus|AUSTR259-13|AUST-261|MK657817
Pervagor marginalis|MARQ163-12|MARQ-163|MK567031
Plagiotremus rhinorhynchos|MARQ205-12|MARQ-206|MK567035
Plagiotremus rhinorhynchos|MARQ204-12|MARQ-205|MK567034
Plagiotremus rhinorhynchos|MARQ203-12|MARQ-204|MK567033
Plagiotremus tapeinosoma|AUSTR269-13|AUST-271|MK657160
Plagiotremus tapeinosoma|AUSTR446-13|AUST-448|MK657767
Plagiotremus tapeinosoma|AUSTR158-13|AUST-160|MK657543
Plagiotremus tapeinosoma|MARQ057-12|MARQ-057|MK567037
Plagiotremus tapeinosoma|MARQ056-12|MARQ-056|MK567036
Plagiotremus tapeinosoma|MOH054-16|MOH-054|MK657566
Plagiotremus tapeinosoma|MARQ200-12|MARQ-201|MK567038
Samariscus triocellatus|MARQ170-12|MARQ-171|MK567091
Samariscus triocellatus|GAMBA071-12|GAM-840|MK658416
Aspidontus taeniatus|MARQ430-12|MARQ-432|MK566803
Aspidontus taeniatus|MARQ194-12|MARQ-195|MK566804
Petroscirtes xestus|SCILL319-15|SCIL-319|MK658698
Petroscirtes xestus|GAMBA805-12|GAM-680|MK658638
Malacanthus brevirostris|MOH144-16|MOH-144|MK657375
Malacanthus brevirostris|MARQ159-12|MARQ-159|MK566979
Stanulus seychellensis|GAMBA266-12|GAM-129|MK657973
Stanulus seychellensis|AUSTR054-13|AUST-054|MK656974
Stanulus seychellensis|GAMBA267-12|GAM-130|MK657578
Stanulus seychellensis|GAMBA265-12|GAM-128|MK658653
Stanulus seychellensis|AUSTR053-13|AUST-053|MK657360
Stanulus seychellensis|SCILL033-15|SCIL-033|MK658394
Stanulus seychellensis|AUSTR055-13|AUST-055|MK657456
Stanulus seychellensis|AUSTR179-13|AUST-181|MK657657
Stanulus talboti|GAMBA647-12|GAM-518|MK658539
Stanulus talboti|GAMBA646-12|GAM-517|MK658676
Stanulus talboti|GAMBA645-12|GAM-516|MK657196
Stanulus talboti|AUSTR175-13|AUST-177|MK657459
Stanulus talboti|AUSTR174-13|AUST-176|MK657934
Stanulus talboti|AUSTR173-13|AUST-175|MK658048
Entomacrodus cymatobiotus|GAMBA644-12|GAM-515|MK657138
Entomacrodus cymatobiotus|GAMBA642-12|GAM-513|MK657419
Entomacrodus|AUSTR407-13|AUST-409|MK658187
Entomacrodus|AUSTR406-13|AUST-408|MK657399
Entomacrodus|AUSTR405-13|AUST-407|MK658111
Entomacrodus caudofasciatus|GAMBA258-12|GAM-121|MK658438
Entomacrodus caudofasciatus|SCILL199-15|SCIL-199|MK657407
Entomacrodus caudofasciatus|SCILL196-15|SCIL-196|MK657152
Entomacrodus caudofasciatus|GAMBA257-12|GAM-120|MK657353
Entomacrodus caudofasciatus|GAMBA256-12|GAM-119|MK657833
Entomacrodus sealei|AUSTR403-13|AUST-405|MK658447
Entomacrodus sealei|AUSTR404-13|AUST-406|MK657699
Entomacrodus sealei|AUSTR402-13|AUST-404|MK657974
Entomacrodus corneliae|MARQ184-12|MARQ-185|MK566898
Entomacrodus niuafoouensis|GAMBA641-12|GAM-512|MK657239
Entomacrodus randalli|MARQ179-12|MARQ-180|MK566901
Entomacrodus thalassinus|GAMBA649-12|GAM-520|MK658079
Entomacrodus thalassinus|AUSTR181-13|AUST-183|MK657392
Entomacrodus thalassinus|SCILL035-15|SCIL-035|MK658549
Entomacrodus thalassinus|GAMBA243-12|GAM-106|MK657149

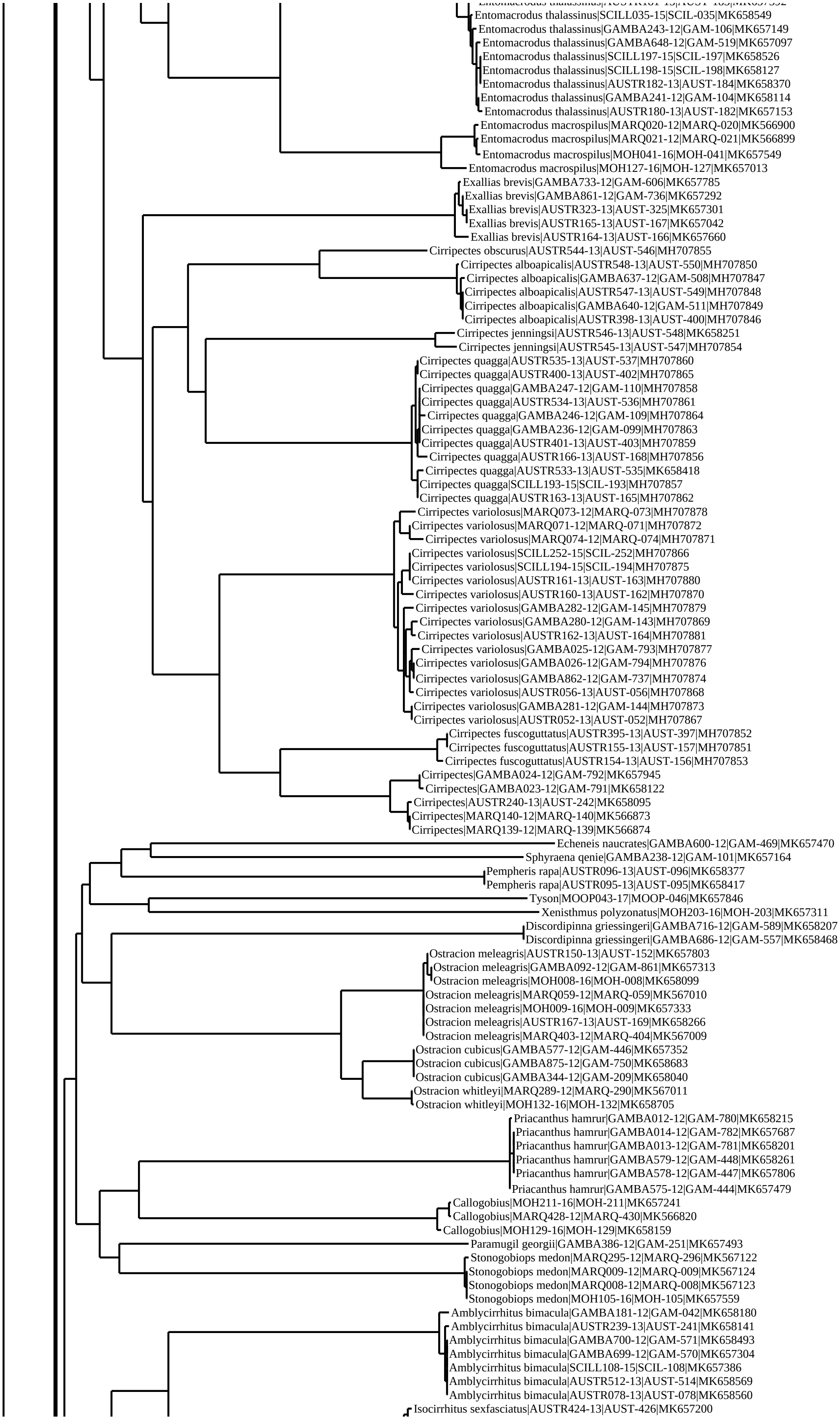
Entomacrodus thalassinus|SCILL035-15|SCIL-035|MK658549
Entomacrodus thalassinus|GAMBA243-12|GAM-106|MK657149
Entomacrodus thalassinus|GAMBA648-12|GAM-519|MK657097
Entomacrodus thalassinus|SCILL197-15|SCIL-197|MK658526
Entomacrodus thalassinus|SCILL198-15|SCIL-198|MK658127
Entomacrodus thalassinus|AUSTR182-13|AUST-184|MK658370
Entomacrodus thalassinus|GAMBA241-12|GAM-104|MK658114
Entomacrodus thalassinus|AUSTR180-13|AUST-182|MK657153
Entomacrodus macrospilus|MARQ020-12|MARQ-020|MK566900
Entomacrodus macrospilus|MARQ021-12|MARQ-021|MK566899
Entomacrodus macrospilus|MOH041-16|MOH-041|MK657549
Entomacrodus macrospilus|MOH127-16|MOH-127|MK657013
Exallias brevis|GAMBA733-12|GAM-606|MK657785
Exallias brevis|GAMBA861-12|GAM-736|MK657292
Exallias brevis|AUSTR323-13|AUST-325|MK657301
Exallias brevis|AUSTR165-13|AUST-167|MK657042
Exallias brevis|AUSTR164-13|AUST-166|MK657660
Cirripectes obscurus|AUSTR544-13|AUST-546|MH707855
Cirripectes alboapicalis|AUSTR548-13|AUST-550|MH707850
Cirripectes alboapicalis|GAMBA637-12|GAM-508|MH707847
Cirripectes alboapicalis|AUSTR547-13|AUST-549|MH707848
Cirripectes alboapicalis|GAMBA640-12|GAM-511|MH707849
Cirripectes alboapicalis|AUSTR398-13|AUST-400|MH707846
Cirripectes jenningsi|AUSTR546-13|AUST-548|MK658251
Cirripectes jenningsi|AUSTR545-13|AUST-547|MH707854
Cirripectes quagga|AUSTR535-13|AUST-537|MH707860
Cirripectes quagga|AUSTR400-13|AUST-402|MH707865
Cirripectes quagga|GAMBA247-12|GAM-110|MH707858
Cirripectes quagga|AUSTR534-13|AUST-536|MH707861
Cirripectes quagga|GAMBA246-12|GAM-109|MH707864
Cirripectes quagga|GAMBA236-12|GAM-099|MH707863
Cirripectes quagga|AUSTR401-13|AUST-403|MH707859
Cirripectes quagga|AUSTR166-13|AUST-168|MH707856
Cirripectes quagga|AUSTR533-13|AUST-535|MK658418
Cirripectes quagga|SCILL193-15|SCIL-193|MH707857
Cirripectes quagga|AUSTR163-13|AUST-165|MH707862
Cirripectes variolosus|MARQ073-12|MARQ-073|MH707878
Cirripectes variolosus|MARQ071-12|MARQ-071|MH707872
Cirripectes variolosus|MARQ074-12|MARQ-074|MH707871
Cirripectes variolosus|SCILL252-15|SCIL-252|MH707866
Cirripectes variolosus|SCILL194-15|SCIL-194|MH707875
Cirripectes variolosus|AUSTR161-13|AUST-163|MH707880
Cirripectes variolosus|AUSTR160-13|AUST-162|MH707870
Cirripectes variolosus|GAMBA282-12|GAM-145|MH707879
Cirripectes variolosus|GAMBA280-12|GAM-143|MH707869
Cirripectes variolosus|AUSTR162-13|AUST-164|MH707881
Cirripectes variolosus|GAMBA025-12|GAM-793|MH707877
Cirripectes variolosus|GAMBA026-12|GAM-794|MH707876
Cirripectes variolosus|GAMBA862-12|GAM-737|MH707874
Cirripectes variolosus|AUSTR056-13|AUST-056|MH707868
Cirripectes variolosus|GAMBA281-12|GAM-144|MH707873
Cirripectes variolosus|AUSTR052-13|AUST-052|MH707867
Cirripectes fuscoguttatus|AUSTR395-13|AUST-397|MH707852
Cirripectes fuscoguttatus|AUSTR155-13|AUST-157|MH707851
Cirripectes fuscoguttatus|AUSTR154-13|AUST-156|MH707853
Cirripectes|GAMBA024-12|GAM-792|MK657945
Cirripectes|GAMBA023-12|GAM-791|MK658122
Cirripectes|AUSTR240-13|AUST-242|MK658095
Cirripectes|MARQ140-12|MARQ-140|MK566873
Cirripectes|MARQ139-12|MARQ-139|MK566874
Echeneis naucrates|GAMBA600-12|GAM-469|MK657470
Sphyraena qenie|GAMBA238-12|GAM-101|MK657164
Pempheris rapa|AUSTR096-13|AUST-096|MK658377
Pempheris rapa|AUSTR095-13|AUST-095|MK658417
Tyson|MOOP043-17|MOOP-046|MK657846
Xenisthmus polyzonatus|MOH203-16|MOH-203|MK657311
Discordipinna griessingeri|GAMBA716-12|GAM-589|MK658207
Discordipinna griessingeri|GAMBA686-12|GAM-557|MK658468
Ostracion meleagris|AUSTR150-13|AUST-152|MK657803
Ostracion meleagris|GAMBA092-12|GAM-861|MK657313
Ostracion meleagris|MOH008-16|MOH-008|MK658099
Ostracion meleagris|MARQ059-12|MARQ-059|MK567010
Ostracion meleagris|MOH009-16|MOH-009|MK657333
Ostracion meleagris|AUSTR167-13|AUST-169|MK658266
Ostracion meleagris|MARQ403-12|MARQ-404|MK567009
Ostracion cubicus|GAMBA577-12|GAM-446|MK657352
Ostracion cubicus|GAMBA875-12|GAM-750|MK658683
Ostracion cubicus|GAMBA344-12|GAM-209|MK658040
Ostracion whitleyi|MARQ289-12|MARQ-290|MK567011
Ostracion whitleyi|MOH132-16|MOH-132|MK658705
Priacanthus hamrur|GAMBA012-12|GAM-780|MK658215
Priacanthus hamrur|GAMBA014-12|GAM-782|MK657687
Priacanthus hamrur|GAMBA013-12|GAM-781|MK658201
Priacanthus hamrur|GAMBA579-12|GAM-448|MK658261
Priacanthus hamrur|GAMBA578-12|GAM-447|MK657806
Priacanthus hamrur|GAMBA575-12|GAM-444|MK657479
Callogobius|MOH211-16|MOH-211|MK657241
Callogobius|MARQ428-12|MARQ-430|MK566820
Callogobius|MOH129-16|MOH-129|MK658159
Paramugil georgii|GAMBA386-12|GAM-251|MK657493
Stonogobiops medon|MARQ295-12|MARQ-296|MK567122
Stonogobiops medon|MARQ009-12|MARQ-009|MK567124
Stonogobiops medon|MARQ008-12|MARQ-008|MK567123
Stonogobiops medon|MOH105-16|MOH-105|MK657559
Amblycirrhitus bimacula|GAMBA181-12|GAM-042|MK658180
Amblycirrhitus bimacula|AUSTR239-13|AUST-241|MK658141
Amblycirrhitus bimacula|GAMBA700-12|GAM-571|MK658493
Amblycirrhitus bimacula|GAMBA699-12|GAM-570|MK657304
Amblycirrhitus bimacula|SCILL108-15|SCIL-108|MK657386
Amblycirrhitus bimacula|AUSTR512-13|AUST-514|MK658569
Amblycirrhitus bimacula|AUSTR078-13|AUST-078|MK658560
Isocirrhitus sexfasciatus|AUSTR424-13|AUST-426|MK657200

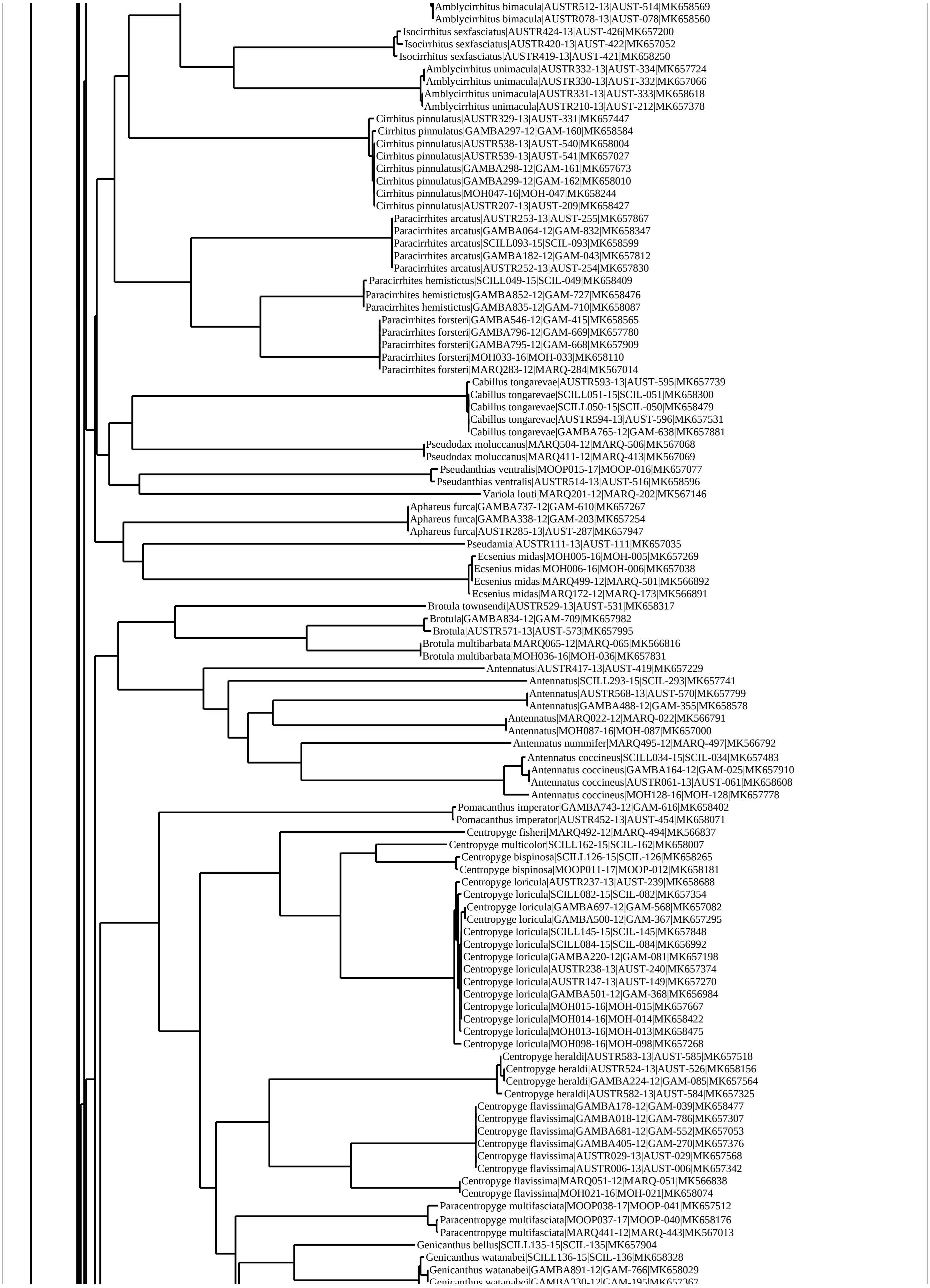
Amblycirrhitus bimacula|AUSTR512-13|AUST-514|MK658569
Amblycirrhitus bimacula|AUSTR078-13|AUST-078|MK658560
Isocirrhitus sexfasciatus|AUSTR424-13|AUST-426|MK657200
Isocirrhitus sexfasciatus|AUSTR420-13|AUST-422|MK657052
Isocirrhitus sexfasciatus|AUSTR419-13|AUST-421|MK658250
Amblycirrhitus unimacula|AUSTR332-13|AUST-334|MK657724
Amblycirrhitus unimacula|AUSTR330-13|AUST-332|MK657066
Amblycirrhitus unimacula|AUSTR331-13|AUST-333|MK658618
Amblycirrhitus unimacula|AUSTR210-13|AUST-212|MK657378
Cirrhitus pinnulatus|AUSTR329-13|AUST-331|MK657447
Cirrhitus pinnulatus|GAMBA297-12|GAM-160|MK658584
Cirrhitus pinnulatus|AUSTR538-13|AUST-540|MK658004
Cirrhitus pinnulatus|AUSTR539-13|AUST-541|MK657027
Cirrhitus pinnulatus|GAMBA298-12|GAM-161|MK657673
Cirrhitus pinnulatus|GAMBA299-12|GAM-162|MK658010
Cirrhitus pinnulatus|MOH047-16|MOH-047|MK658244
Cirrhitus pinnulatus|AUSTR207-13|AUST-209|MK658427
Paracirrhites arcatus|AUSTR253-13|AUST-255|MK657867
Paracirrhites arcatus|GAMBA064-12|GAM-832|MK658347
Paracirrhites arcatus|SCILL093-15|SCIL-093|MK658599
Paracirrhites arcatus|GAMBA182-12|GAM-043|MK657812
Paracirrhites arcatus|AUSTR252-13|AUST-254|MK657830
Paracirrhites hemistictus|SCILL049-15|SCIL-049|MK658409
Paracirrhites hemistictus|GAMBA852-12|GAM-727|MK658476
Paracirrhites hemistictus|GAMBA835-12|GAM-710|MK658087
Paracirrhites forsteri|GAMBA546-12|GAM-415|MK658565
Paracirrhites forsteri|GAMBA796-12|GAM-669|MK657780
Paracirrhites forsteri|GAMBA795-12|GAM-668|MK657909
Paracirrhites forsteri|MOH033-16|MOH-033|MK658110
Paracirrhites forsteri|MARQ283-12|MARQ-284|MK567014
Cabillus tongarevae|AUSTR593-13|AUST-595|MK657739
Cabillus tongarevae|SCILL051-15|SCIL-051|MK658300
Cabillus tongarevae|SCILL050-15|SCIL-050|MK658479
Cabillus tongarevae|AUSTR594-13|AUST-596|MK657531
Cabillus tongarevae|GAMBA765-12|GAM-638|MK657881
Pseudodax moluccanus|MARQ504-12|MARQ-506|MK567068
Pseudodax moluccanus|MARQ411-12|MARQ-413|MK567069
Pseudanthias ventralis|MOOP015-17|MOOP-016|MK657077
Pseudanthias ventralis|AUSTR514-13|AUST-516|MK658596
Variola louti|MARQ201-12|MARQ-202|MK567146
Aphareus furca|GAMBA737-12|GAM-610|MK657267
Aphareus furca|GAMBA338-12|GAM-203|MK657254
Aphareus furca|AUSTR285-13|AUST-287|MK657947
Pseudamia|AUSTR111-13|AUST-111|MK657035
Ecsenius midas|MOH005-16|MOH-005|MK657269
Ecsenius midas|MOH006-16|MOH-006|MK657038
Ecsenius midas|MARQ499-12|MARQ-501|MK566892
Ecsenius midas|MARQ172-12|MARQ-173|MK566891
Brotula townsendi|AUSTR529-13|AUST-531|MK658317
Brotula|GAMBA834-12|GAM-709|MK657982
Brotula|AUSTR571-13|AUST-573|MK657995
Brotula multibarbata|MARQ065-12|MARQ-065|MK566816
Brotula multibarbata|MOH036-16|MOH-036|MK657831
Antennatus|AUSTR417-13|AUST-419|MK657229
Antennatus|SCILL293-15|SCIL-293|MK657741
Antennatus|AUSTR568-13|AUST-570|MK657799
Antennatus|GAMBA488-12|GAM-355|MK658578
Antennatus|MARQ022-12|MARQ-022|MK566791
Antennatus|MOH087-16|MOH-087|MK657000
Antennatus nummifer|MARQ495-12|MARQ-497|MK566792
Antennatus coccineus|SCILL034-15|SCIL-034|MK657483
Antennatus coccineus|GAMBA164-12|GAM-025|MK657910
Antennatus coccineus|AUSTR061-13|AUST-061|MK658608
Antennatus coccineus|MOH128-16|MOH-128|MK657778
Pomacanthus imperator|GAMBA743-12|GAM-616|MK658402
Pomacanthus imperator|AUSTR452-13|AUST-454|MK658071
Centropyge fisheri|MARQ492-12|MARQ-494|MK566837
Centropyge multicolor|SCILL162-15|SCIL-162|MK658007
Centropyge bispinosa|SCILL126-15|SCIL-126|MK658265
Centropyge bispinosa|MOOP011-17|MOOP-012|MK658181
Centropyge loricula|AUSTR237-13|AUST-239|MK658688
Centropyge loricula|SCILL082-15|SCIL-082|MK657354
Centropyge loricula|GAMBA697-12|GAM-568|MK657082
Centropyge loricula|GAMBA500-12|GAM-367|MK657295
Centropyge loricula|SCILL145-15|SCIL-145|MK657848
Centropyge loricula|SCILL084-15|SCIL-084|MK656992
Centropyge loricula|GAMBA220-12|GAM-081|MK657198
Centropyge loricula|AUSTR238-13|AUST-240|MK657374
Centropyge loricula|AUSTR147-13|AUST-149|MK657270
Centropyge loricula|GAMBA501-12|GAM-368|MK656984
Centropyge loricula|MOH015-16|MOH-015|MK657667
Centropyge loricula|MOH014-16|MOH-014|MK658422
Centropyge loricula|MOH013-16|MOH-013|MK658475
Centropyge loricula|MOH098-16|MOH-098|MK657268
Centropyge heraldi|AUSTR583-13|AUST-585|MK657518
Centropyge heraldi|AUSTR524-13|AUST-526|MK658156
Centropyge heraldi|GAMBA224-12|GAM-085|MK657564
Centropyge heraldi|AUSTR582-13|AUST-584|MK657325
Centropyge flavissima|GAMBA178-12|GAM-039|MK658477
Centropyge flavissima|GAMBA018-12|GAM-786|MK657307
Centropyge flavissima|GAMBA681-12|GAM-552|MK657053
Centropyge flavissima|GAMBA405-12|GAM-270|MK657376
Centropyge flavissima|AUSTR029-13|AUST-029|MK657568
Centropyge flavissima|AUSTR006-13|AUST-006|MK657342
Centropyge flavissima|MARQ051-12|MARQ-051|MK566838
Centropyge flavissima|MOH021-16|MOH-021|MK658074
Paracentropyge multifasciata|MOOP038-17|MOOP-041|MK657512
Paracentropyge multifasciata|MOOP037-17|MOOP-040|MK658176
Paracentropyge multifasciata|MARQ441-12|MARQ-443|MK567013
Genicanthus bellus|SCILL135-15|SCIL-135|MK657904
Genicanthus watanabei|SCILL136-15|SCIL-136|MK658328
Genicanthus watanabei|GAMBA891-12|GAM-766|MK658029
Genicanthus watanabei|GAMBA330-12|GAM-195|MK657367

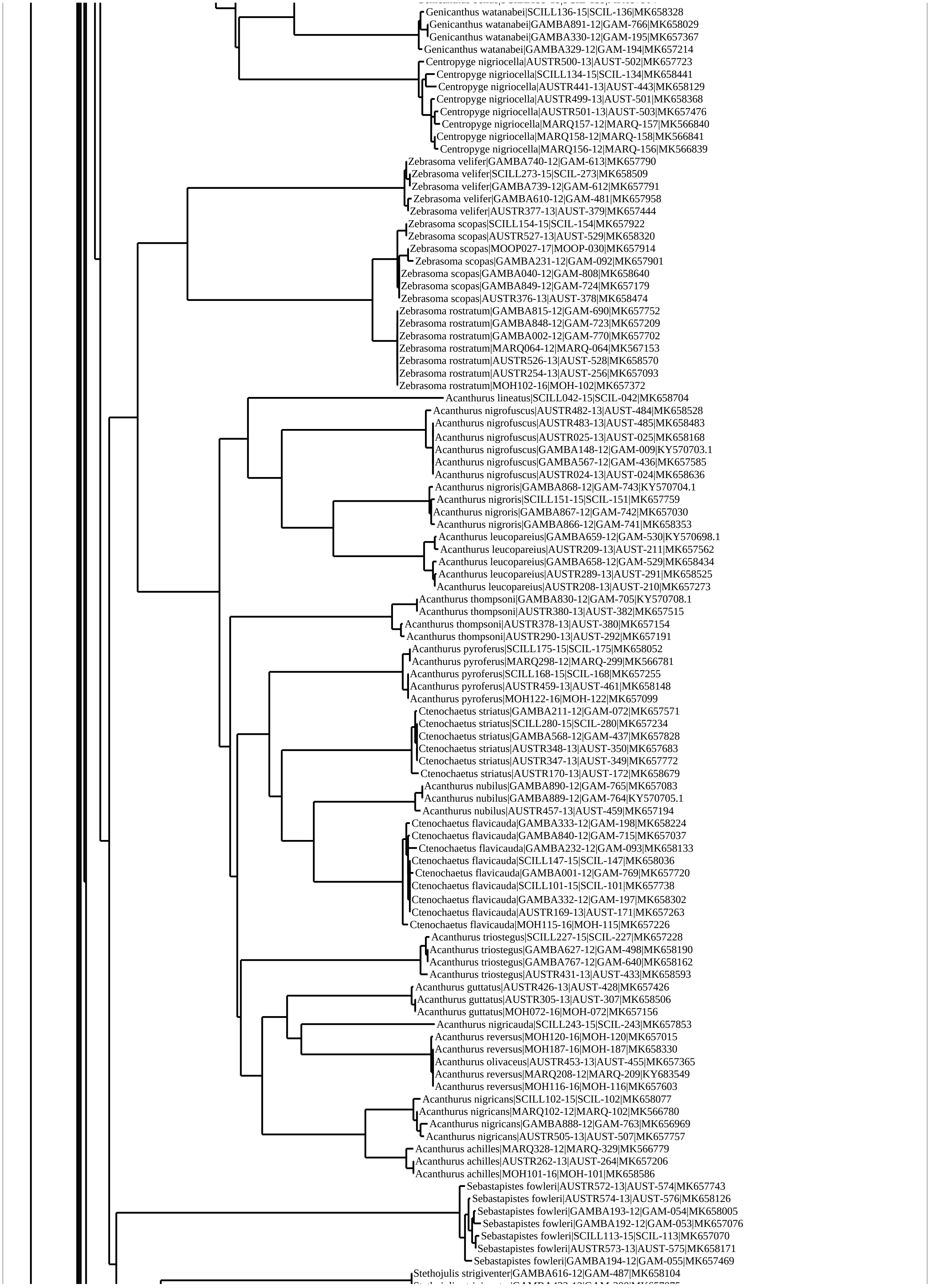
Genicanthus watanabei|SCILL136-15|SCIL-136|MK658328
Genicanthus watanabei|GAMBA891-12|GAM-766|MK658029
Genicanthus watanabei|GAMBA330-12|GAM-195|MK657367
Genicanthus watanabei|GAMBA329-12|GAM-194|MK657214
Centropyge nigriocella|AUSTR500-13|AUST-502|MK657723
Centropyge nigriocella|SCILL134-15|SCIL-134|MK658441
Centropyge nigriocella|AUSTR441-13|AUST-443|MK658129
Centropyge nigriocella|AUSTR499-13|AUST-501|MK658368
Centropyge nigriocella|AUSTR501-13|AUST-503|MK657476
Centropyge nigriocella|MARQ157-12|MARQ-157|MK566840
Centropyge nigriocella|MARQ158-12|MARQ-158|MK566841
Centropyge nigriocella|MARQ156-12|MARQ-156|MK566839
Zebrasoma velifer|GAMBA740-12|GAM-613|MK657790
Zebrasoma velifer|SCILL273-15|SCIL-273|MK658509
Zebrasoma velifer|GAMBA739-12|GAM-612|MK657791
Zebrasoma velifer|GAMBA610-12|GAM-481|MK657958
Zebrasoma velifer|AUSTR377-13|AUST-379|MK657444
Zebrasoma scopas|SCILL154-15|SCIL-154|MK657922
Zebrasoma scopas|AUSTR527-13|AUST-529|MK658320
Zebrasoma scopas|MOOP027-17|MOOP-030|MK657914
Zebrasoma scopas|GAMBA231-12|GAM-092|MK657901
Zebrasoma scopas|GAMBA040-12|GAM-808|MK658640
Zebrasoma scopas|GAMBA849-12|GAM-724|MK657179
Zebrasoma scopas|AUSTR376-13|AUST-378|MK658474
Zebrasoma rostratum|GAMBA815-12|GAM-690|MK657752
Zebrasoma rostratum|GAMBA848-12|GAM-723|MK657209
Zebrasoma rostratum|GAMBA002-12|GAM-770|MK657702
Zebrasoma rostratum|MARQ064-12|MARQ-064|MK567153
Zebrasoma rostratum|AUSTR526-13|AUST-528|MK658570
Zebrasoma rostratum|AUSTR254-13|AUST-256|MK657093
Zebrasoma rostratum|MOH102-16|MOH-102|MK657372
Acanthurus lineatus|SCILL042-15|SCIL-042|MK658704
Acanthurus nigrofuscus|AUSTR482-13|AUST-484|MK658528
Acanthurus nigrofuscus|AUSTR483-13|AUST-485|MK658483
Acanthurus nigrofuscus|AUSTR025-13|AUST-025|MK658168
Acanthurus nigrofuscus|GAMBA148-12|GAM-009|KY570703.1
Acanthurus nigrofuscus|GAMBA567-12|GAM-436|MK657585
Acanthurus nigrofuscus|AUSTR024-13|AUST-024|MK658636
Acanthurus nigroris|GAMBA868-12|GAM-743|KY570704.1
Acanthurus nigroris|SCILL151-15|SCIL-151|MK657759
Acanthurus nigroris|GAMBA867-12|GAM-742|MK657030
Acanthurus nigroris|GAMBA866-12|GAM-741|MK658353
Acanthurus leucopareius|GAMBA659-12|GAM-530|KY570698.1
Acanthurus leucopareius|AUSTR209-13|AUST-211|MK657562
Acanthurus leucopareius|GAMBA658-12|GAM-529|MK658434
Acanthurus leucopareius|AUSTR289-13|AUST-291|MK658525
Acanthurus leucopareius|AUSTR208-13|AUST-210|MK657273
Acanthurus thompsoni|GAMBA830-12|GAM-705|KY570708.1
Acanthurus thompsoni|AUSTR380-13|AUST-382|MK657515
Acanthurus thompsoni|AUSTR378-13|AUST-380|MK657154
Acanthurus thompsoni|AUSTR290-13|AUST-292|MK657191
Acanthurus pyroferus|SCILL175-15|SCIL-175|MK658052
Acanthurus pyroferus|MARQ298-12|MARQ-299|MK566781
Acanthurus pyroferus|SCILL168-15|SCIL-168|MK657255
Acanthurus pyroferus|AUSTR459-13|AUST-461|MK658148
Acanthurus pyroferus|MOH122-16|MOH-122|MK657099
Ctenochaetus striatus|GAMBA211-12|GAM-072|MK657571
Ctenochaetus striatus|SCILL280-15|SCIL-280|MK657234
Ctenochaetus striatus|GAMBA568-12|GAM-437|MK657828
Ctenochaetus striatus|AUSTR348-13|AUST-350|MK657683
Ctenochaetus striatus|AUSTR347-13|AUST-349|MK657772
Ctenochaetus striatus|AUSTR170-13|AUST-172|MK658679
Acanthurus nubilus|GAMBA890-12|GAM-765|MK657083
Acanthurus nubilus|GAMBA889-12|GAM-764|KY570705.1
Acanthurus nubilus|AUSTR457-13|AUST-459|MK657194
Ctenochaetus flavicauda|GAMBA333-12|GAM-198|MK658224
Ctenochaetus flavicauda|GAMBA840-12|GAM-715|MK657037
Ctenochaetus flavicauda|GAMBA232-12|GAM-093|MK658133
Ctenochaetus flavicauda|SCILL147-15|SCIL-147|MK658036
Ctenochaetus flavicauda|GAMBA001-12|GAM-769|MK657720
Ctenochaetus flavicauda|SCILL101-15|SCIL-101|MK657738
Ctenochaetus flavicauda|GAMBA332-12|GAM-197|MK658302
Ctenochaetus flavicauda|AUSTR169-13|AUST-171|MK657263
Ctenochaetus flavicauda|MOH115-16|MOH-115|MK657226
Acanthurus triostegus|SCILL227-15|SCIL-227|MK657228
Acanthurus triostegus|GAMBA627-12|GAM-498|MK658190
Acanthurus triostegus|GAMBA767-12|GAM-640|MK658162
Acanthurus triostegus|AUSTR431-13|AUST-433|MK658593
Acanthurus guttatus|AUSTR426-13|AUST-428|MK657426
Acanthurus guttatus|AUSTR305-13|AUST-307|MK658506
Acanthurus guttatus|MOH072-16|MOH-072|MK657156
Acanthurus nigricauda|SCILL243-15|SCIL-243|MK657853
Acanthurus reversus|MOH120-16|MOH-120|MK657015
Acanthurus reversus|MOH187-16|MOH-187|MK658330
Acanthurus olivaceus|AUSTR453-13|AUST-455|MK657365
Acanthurus reversus|MARQ208-12|MARQ-209|KY683549
Acanthurus reversus|MOH116-16|MOH-116|MK657603
Acanthurus nigricans|SCILL102-15|SCIL-102|MK658077
Acanthurus nigricans|MARQ102-12|MARQ-102|MK566780
Acanthurus nigricans|GAMBA888-12|GAM-763|MK656969
Acanthurus nigricans|AUSTR505-13|AUST-507|MK657757
Acanthurus achilles|MARQ328-12|MARQ-329|MK566779
Acanthurus achilles|AUSTR262-13|AUST-264|MK657206
Acanthurus achilles|MOH101-16|MOH-101|MK658586
Sebastapistes fowleri|AUSTR572-13|AUST-574|MK657743
Sebastapistes fowleri|AUSTR574-13|AUST-576|MK658126
Sebastapistes fowleri|GAMBA193-12|GAM-054|MK658005
Sebastapistes fowleri|GAMBA192-12|GAM-053|MK657076
Sebastapistes fowleri|SCILL113-15|SCIL-113|MK657070
Sebastapistes fowleri|AUSTR573-13|AUST-575|MK658171
Sebastapistes fowleri|GAMBA194-12|GAM-055|MK657469
Stethojulis strigiventer|GAMBA616-12|GAM-487|MK658104

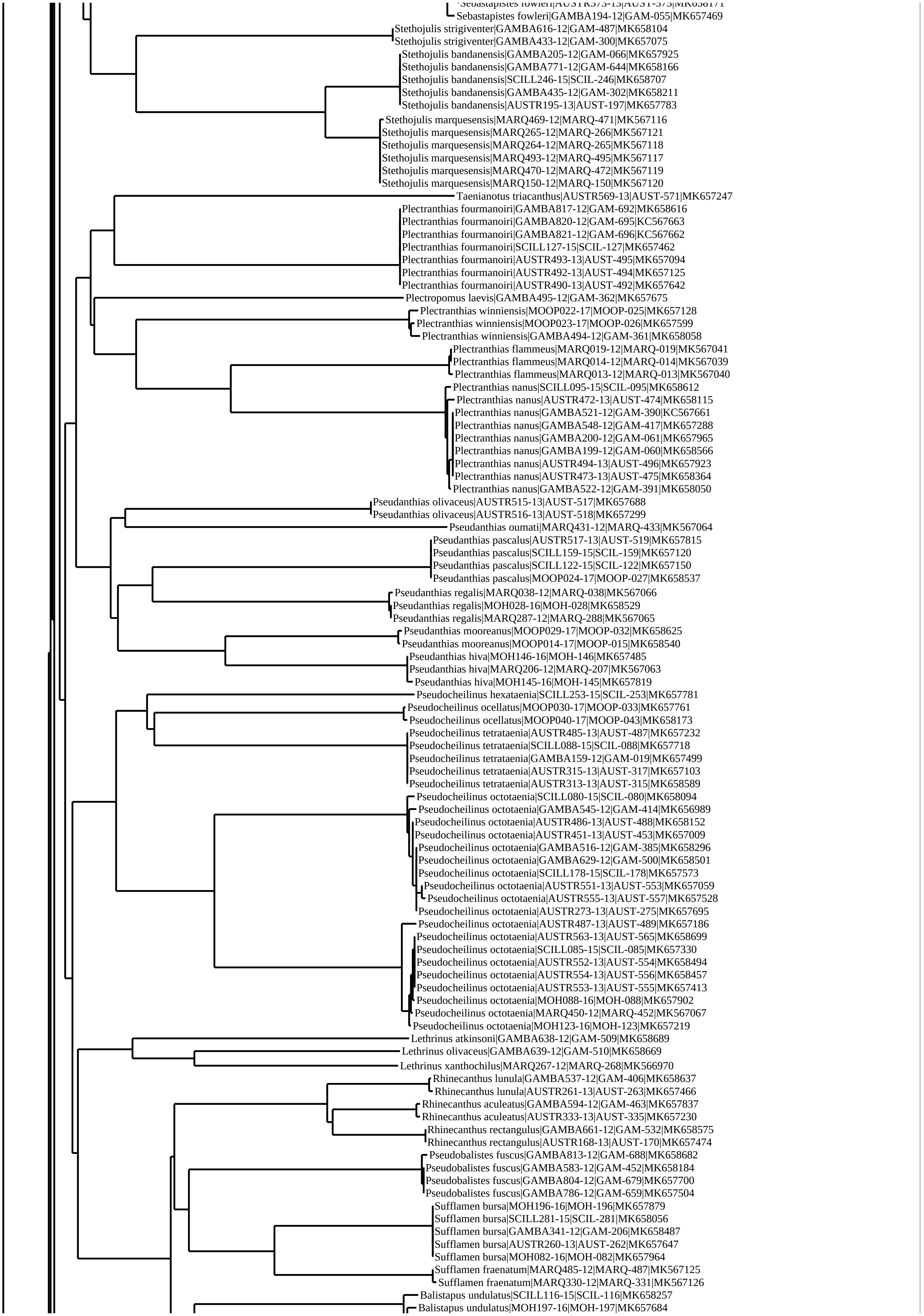
Sebastapistes fowleri|AUSTR573-13|AUST-575|MK658171
Sebastapistes fowleri|GAMBA194-12|GAM-055|MK657469
Stethojulis strigiventer|GAMBA616-12|GAM-487|MK658104
Stethojulis strigiventer|GAMBA433-12|GAM-300|MK657075
Stethojulis bandanensis|GAMBA205-12|GAM-066|MK657925
Stethojulis bandanensis|GAMBA771-12|GAM-644|MK658166
Stethojulis bandanensis|SCILL246-15|SCIL-246|MK658707
Stethojulis bandanensis|GAMBA435-12|GAM-302|MK658211
Stethojulis bandanensis|AUSTR195-13|AUST-197|MK657783
Stethojulis marquesensis|MARQ469-12|MARQ-471|MK567116
Stethojulis marquesensis|MARQ265-12|MARQ-266|MK567121
Stethojulis marquesensis|MARQ264-12|MARQ-265|MK567118
Stethojulis marquesensis|MARQ493-12|MARQ-495|MK567117
Stethojulis marquesensis|MARQ470-12|MARQ-472|MK567119
Stethojulis marquesensis|MARQ150-12|MARQ-150|MK567120
Taenianotus triacanthus|AUSTR569-13|AUST-571|MK657247
Plectranthias fourmanoiri|GAMBA817-12|GAM-692|MK658616
Plectranthias fourmanoiri|GAMBA820-12|GAM-695|KC567663
Plectranthias fourmanoiri|GAMBA821-12|GAM-696|KC567662
Plectranthias fourmanoiri|SCILL127-15|SCIL-127|MK657462
Plectranthias fourmanoiri|AUSTR493-13|AUST-495|MK657094
Plectranthias fourmanoiri|AUSTR492-13|AUST-494|MK657125
Plectranthias fourmanoiri|AUSTR490-13|AUST-492|MK657642
Plectropomus laevis|GAMBA495-12|GAM-362|MK657675
Plectranthias winniensis|MOOP022-17|MOOP-025|MK657128
Plectranthias winniensis|MOOP023-17|MOOP-026|MK657599
Plectranthias winniensis|GAMBA494-12|GAM-361|MK658058
Plectranthias flammeus|MARQ019-12|MARQ-019|MK567041
Plectranthias flammeus|MARQ014-12|MARQ-014|MK567039
Plectranthias flammeus|MARQ013-12|MARQ-013|MK567040
Plectranthias nanus|SCILL095-15|SCIL-095|MK658612
Plectranthias nanus|AUSTR472-13|AUST-474|MK658115
Plectranthias nanus|GAMBA521-12|GAM-390|KC567661
Plectranthias nanus|GAMBA548-12|GAM-417|MK657288
Plectranthias nanus|GAMBA200-12|GAM-061|MK657965
Plectranthias nanus|GAMBA199-12|GAM-060|MK658566
Plectranthias nanus|AUSTR494-13|AUST-496|MK657923
Plectranthias nanus|AUSTR473-13|AUST-475|MK658364
Plectranthias nanus|GAMBA522-12|GAM-391|MK658050
Pseudanthias olivaceus|AUSTR515-13|AUST-517|MK657688
Pseudanthias olivaceus|AUSTR516-13|AUST-518|MK657299
Pseudanthias oumati|MARQ431-12|MARQ-433|MK567064
Pseudanthias pascalus|AUSTR517-13|AUST-519|MK657815
Pseudanthias pascalus|SCILL159-15|SCIL-159|MK657120
Pseudanthias pascalus|SCILL122-15|SCIL-122|MK657150
Pseudanthias pascalus|MOOP024-17|MOOP-027|MK658537
Pseudanthias regalis|MARQ038-12|MARQ-038|MK567066
Pseudanthias regalis|MOH028-16|MOH-028|MK658529
Pseudanthias regalis|MARQ287-12|MARQ-288|MK567065
Pseudanthias mooreanus|MOOP029-17|MOOP-032|MK658625
Pseudanthias mooreanus|MOOP014-17|MOOP-015|MK658540
Pseudanthias hiva|MOH146-16|MOH-146|MK657485
Pseudanthias hiva|MARQ206-12|MARQ-207|MK567063
Pseudanthias hiva|MOH145-16|MOH-145|MK657819
Pseudocheilinus hexataenia|SCILL253-15|SCIL-253|MK657781
Pseudocheilinus ocellatus|MOOP030-17|MOOP-033|MK657761
Pseudocheilinus ocellatus|MOOP040-17|MOOP-043|MK658173
Pseudocheilinus tetrataenia|AUSTR485-13|AUST-487|MK657232
Pseudocheilinus tetrataenia|SCILL088-15|SCIL-088|MK657718
Pseudocheilinus tetrataenia|GAMBA159-12|GAM-019|MK657499
Pseudocheilinus tetrataenia|AUSTR315-13|AUST-317|MK657103
Pseudocheilinus tetrataenia|AUSTR313-13|AUST-315|MK658589
Pseudocheilinus octotaenia|SCILL080-15|SCIL-080|MK658094
Pseudocheilinus octotaenia|GAMBA545-12|GAM-414|MK656989
Pseudocheilinus octotaenia|AUSTR486-13|AUST-488|MK658152
Pseudocheilinus octotaenia|AUSTR451-13|AUST-453|MK657009
Pseudocheilinus octotaenia|GAMBA516-12|GAM-385|MK658296
Pseudocheilinus octotaenia|GAMBA629-12|GAM-500|MK658501
Pseudocheilinus octotaenia|SCILL178-15|SCIL-178|MK657573
Pseudocheilinus octotaenia|AUSTR551-13|AUST-553|MK657059
Pseudocheilinus octotaenia|AUSTR555-13|AUST-557|MK657528
Pseudocheilinus octotaenia|AUSTR273-13|AUST-275|MK657695
Pseudocheilinus octotaenia|AUSTR487-13|AUST-489|MK657186
Pseudocheilinus octotaenia|AUSTR563-13|AUST-565|MK658699
Pseudocheilinus octotaenia|SCILL085-15|SCIL-085|MK657330
Pseudocheilinus octotaenia|AUSTR552-13|AUST-554|MK658494
Pseudocheilinus octotaenia|AUSTR554-13|AUST-556|MK658457
Pseudocheilinus octotaenia|AUSTR553-13|AUST-555|MK657413
Pseudocheilinus octotaenia|MOH088-16|MOH-088|MK657902
Pseudocheilinus octotaenia|MARQ450-12|MARQ-452|MK567067
Pseudocheilinus octotaenia|MOH123-16|MOH-123|MK657219
Lethrinus atkinsoni|GAMBA638-12|GAM-509|MK658689
Lethrinus olivaceus|GAMBA639-12|GAM-510|MK658669
Lethrinus xanthochilus|MARQ267-12|MARQ-268|MK566970
Rhinecanthus lunula|GAMBA537-12|GAM-406|MK658637
Rhinecanthus lunula|AUSTR261-13|AUST-263|MK657466
Rhinecanthus aculeatus|GAMBA594-12|GAM-463|MK657837
Rhinecanthus aculeatus|AUSTR333-13|AUST-335|MK657230
Rhinecanthus rectangulus|GAMBA661-12|GAM-532|MK658575
Rhinecanthus rectangulus|AUSTR168-13|AUST-170|MK657474
Pseudobalistes fuscus|GAMBA813-12|GAM-688|MK658682
Pseudobalistes fuscus|GAMBA583-12|GAM-452|MK658184
Pseudobalistes fuscus|GAMBA804-12|GAM-679|MK657700
Pseudobalistes fuscus|GAMBA786-12|GAM-659|MK657504
Sufflamen bursa|MOH196-16|MOH-196|MK657879
Sufflamen bursa|SCILL281-15|SCIL-281|MK658056
Sufflamen bursa|GAMBA341-12|GAM-206|MK658487
Sufflamen bursa|AUSTR260-13|AUST-262|MK657647
Sufflamen bursa|MOH082-16|MOH-082|MK657964
Sufflamen fraenatum|MARQ485-12|MARQ-487|MK567125
Sufflamen fraenatum|MARQ330-12|MARQ-331|MK567126
Balistapus undulatus|SCILL116-15|SCIL-116|MK658257
Balistapus undulatus|MOH197-16|MOH-197|MK657684

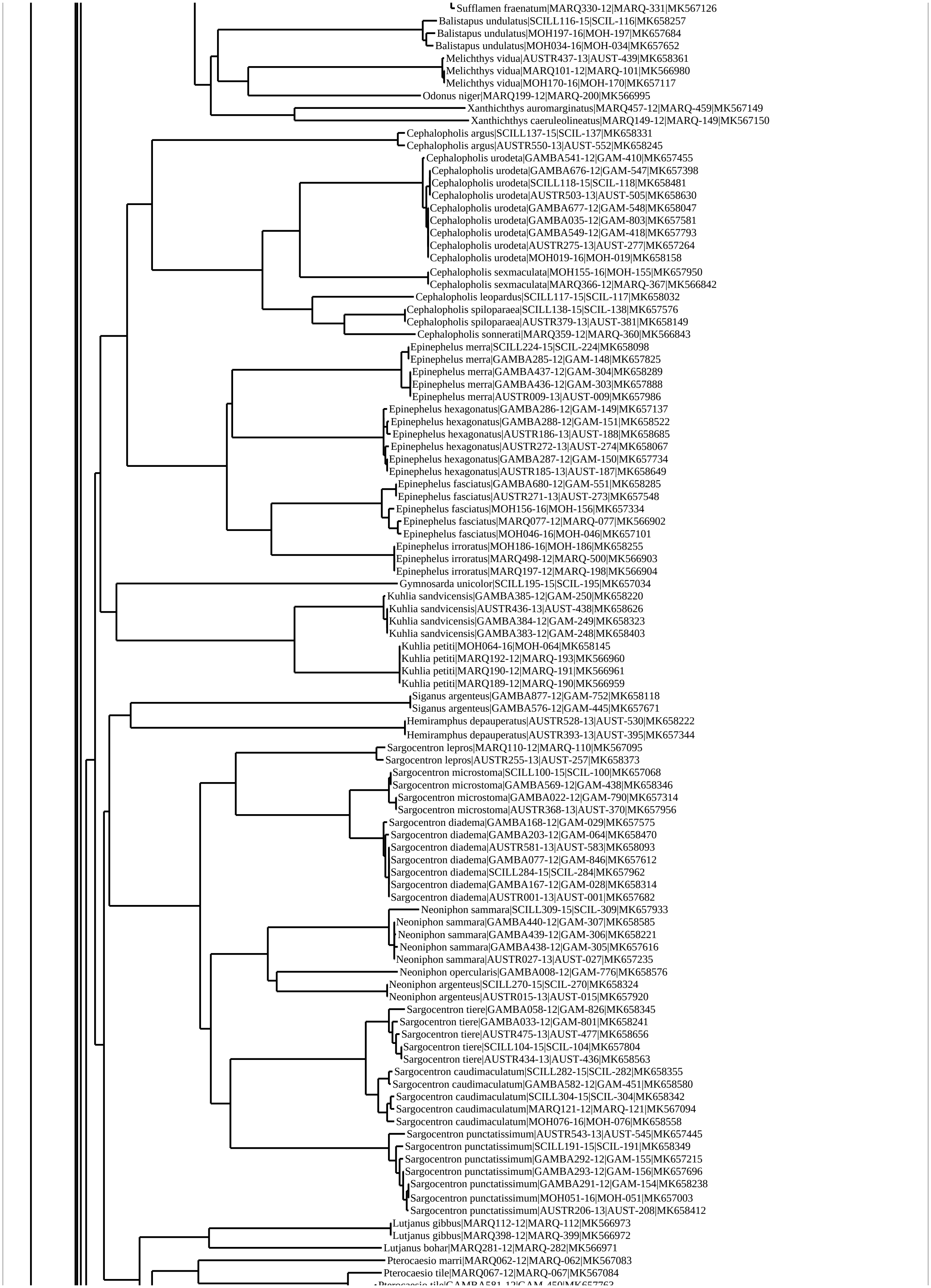
Sufflamen fraenatum|MARQ330-12|MARQ-331|MK567126
Balistapus undulatus|SCILL116-15|SCIL-116|MK658257
Balistapus undulatus|MOH197-16|MOH-197|MK657684
Balistapus undulatus|MOH034-16|MOH-034|MK657652
Melichthys vidua|AUSTR437-13|AUST-439|MK658361
Melichthys vidua|MARQ101-12|MARQ-101|MK566980
Melichthys vidua|MOH170-16|MOH-170|MK657117
Odonus niger|MARQ199-12|MARQ-200|MK566995
Xanthichthys auromarginatus|MARQ457-12|MARQ-459|MK567149
Xanthichthys caeruleolineatus|MARQ149-12|MARQ-149|MK567150
Cephalopholis argus|SCILL137-15|SCIL-137|MK658331
Cephalopholis argus|AUSTR550-13|AUST-552|MK658245
Cephalopholis urodeta|GAMBA541-12|GAM-410|MK657455
Cephalopholis urodeta|GAMBA676-12|GAM-547|MK657398
Cephalopholis urodeta|SCILL118-15|SCIL-118|MK658481
Cephalopholis urodeta|AUSTR503-13|AUST-505|MK658630
Cephalopholis urodeta|GAMBA677-12|GAM-548|MK658047
Cephalopholis urodeta|GAMBA035-12|GAM-803|MK657581
Cephalopholis urodeta|GAMBA549-12|GAM-418|MK657793
Cephalopholis urodeta|AUSTR275-13|AUST-277|MK657264
Cephalopholis urodeta|MOH019-16|MOH-019|MK658158
Cephalopholis sexmaculata|MOH155-16|MOH-155|MK657950
Cephalopholis sexmaculata|MARQ366-12|MARQ-367|MK566842
Cephalopholis leopardus|SCILL117-15|SCIL-117|MK658032
Cephalopholis spiloparaea|SCILL138-15|SCIL-138|MK657576
Cephalopholis spiloparaea|AUSTR379-13|AUST-381|MK658149
Cephalopholis sonnerati|MARQ359-12|MARQ-360|MK566843
Epinephelus merra|SCILL224-15|SCIL-224|MK658098
Epinephelus merra|GAMBA285-12|GAM-148|MK657825
Epinephelus merra|GAMBA437-12|GAM-304|MK658289
Epinephelus merra|GAMBA436-12|GAM-303|MK657888
Epinephelus merra|AUSTR009-13|AUST-009|MK657986
Epinephelus hexagonatus|GAMBA286-12|GAM-149|MK657137
Epinephelus hexagonatus|GAMBA288-12|GAM-151|MK658522
Epinephelus hexagonatus|AUSTR186-13|AUST-188|MK658685
Epinephelus hexagonatus|AUSTR272-13|AUST-274|MK658067
Epinephelus hexagonatus|GAMBA287-12|GAM-150|MK657734
Epinephelus hexagonatus|AUSTR185-13|AUST-187|MK658649
Epinephelus fasciatus|GAMBA680-12|GAM-551|MK658285
Epinephelus fasciatus|AUSTR271-13|AUST-273|MK657548
Epinephelus fasciatus|MOH156-16|MOH-156|MK657334
Epinephelus fasciatus|MARQ077-12|MARQ-077|MK566902
Epinephelus fasciatus|MOH046-16|MOH-046|MK657101
Epinephelus irroratus|MOH186-16|MOH-186|MK658255
Epinephelus irroratus|MARQ498-12|MARQ-500|MK566903
Epinephelus irroratus|MARQ197-12|MARQ-198|MK566904
Gymnosarda unicolor|SCILL195-15|SCIL-195|MK657034
Kuhlia sandvicensis|GAMBA385-12|GAM-250|MK658220
Kuhlia sandvicensis|AUSTR436-13|AUST-438|MK658626
Kuhlia sandvicensis|GAMBA384-12|GAM-249|MK658323
Kuhlia sandvicensis|GAMBA383-12|GAM-248|MK658403
Kuhlia petiti|MOH064-16|MOH-064|MK658145
Kuhlia petiti|MARQ192-12|MARQ-193|MK566960
Kuhlia petiti|MARQ190-12|MARQ-191|MK566961
Kuhlia petiti|MARQ189-12|MARQ-190|MK566959
Siganus argenteus|GAMBA877-12|GAM-752|MK658118
Siganus argenteus|GAMBA576-12|GAM-445|MK657671
Hemiramphus depauperatus|AUSTR528-13|AUST-530|MK658222
Hemiramphus depauperatus|AUSTR393-13|AUST-395|MK657344
Sargocentron lepros|MARQ110-12|MARQ-110|MK567095
Sargocentron lepros|AUSTR255-13|AUST-257|MK658373
Sargocentron microstoma|SCILL100-15|SCIL-100|MK657068
Sargocentron microstoma|GAMBA569-12|GAM-438|MK658346
Sargocentron microstoma|GAMBA022-12|GAM-790|MK657314
Sargocentron microstoma|AUSTR368-13|AUST-370|MK657956
Sargocentron diadema|GAMBA168-12|GAM-029|MK657575
Sargocentron diadema|GAMBA203-12|GAM-064|MK658470
Sargocentron diadema|AUSTR581-13|AUST-583|MK658093
Sargocentron diadema|GAMBA077-12|GAM-846|MK657612
Sargocentron diadema|SCILL284-15|SCIL-284|MK657962
Sargocentron diadema|GAMBA167-12|GAM-028|MK658314
Sargocentron diadema|AUSTR001-13|AUST-001|MK657682
Neoniphon sammara|SCILL309-15|SCIL-309|MK657933
Neoniphon sammara|GAMBA440-12|GAM-307|MK658585
Neoniphon sammara|GAMBA439-12|GAM-306|MK658221
Neoniphon sammara|GAMBA438-12|GAM-305|MK657616
Neoniphon sammara|AUSTR027-13|AUST-027|MK657235
Neoniphon opercularis|GAMBA008-12|GAM-776|MK658576
Neoniphon argenteus|SCILL270-15|SCIL-270|MK658324
Neoniphon argenteus|AUSTR015-13|AUST-015|MK657920
Sargocentron tiere|GAMBA058-12|GAM-826|MK658345
Sargocentron tiere|GAMBA033-12|GAM-801|MK658241
Sargocentron tiere|AUSTR475-13|AUST-477|MK658656
Sargocentron tiere|SCILL104-15|SCIL-104|MK657804
Sargocentron tiere|AUSTR434-13|AUST-436|MK658563
Sargocentron caudimaculatum|SCILL282-15|SCIL-282|MK658355
Sargocentron caudimaculatum|GAMBA582-12|GAM-451|MK658580
Sargocentron caudimaculatum|SCILL304-15|SCIL-304|MK658342
Sargocentron caudimaculatum|MARQ121-12|MARQ-121|MK567094
Sargocentron caudimaculatum|MOH076-16|MOH-076|MK658558
Sargocentron punctatissimum|AUSTR543-13|AUST-545|MK657445
Sargocentron punctatissimum|SCILL191-15|SCIL-191|MK658349
Sargocentron punctatissimum|GAMBA292-12|GAM-155|MK657215
Sargocentron punctatissimum|GAMBA293-12|GAM-156|MK657696
Sargocentron punctatissimum|GAMBA291-12|GAM-154|MK658238
Sargocentron punctatissimum|MOH051-16|MOH-051|MK657003
Sargocentron punctatissimum|AUSTR206-13|AUST-208|MK658412
Lutjanus gibbus|MARQ112-12|MARQ-112|MK566973
Lutjanus gibbus|MARQ398-12|MARQ-399|MK566972
Lutjanus bohar|MARQ281-12|MARQ-282|MK566971
Pterocaesio marri|MARQ062-12|MARQ-062|MK567083
Pterocaesio tile|MARQ067-12|MARQ-067|MK567084
Pterocaesio tile|GAMBA581-12|GAM-450|MK657763

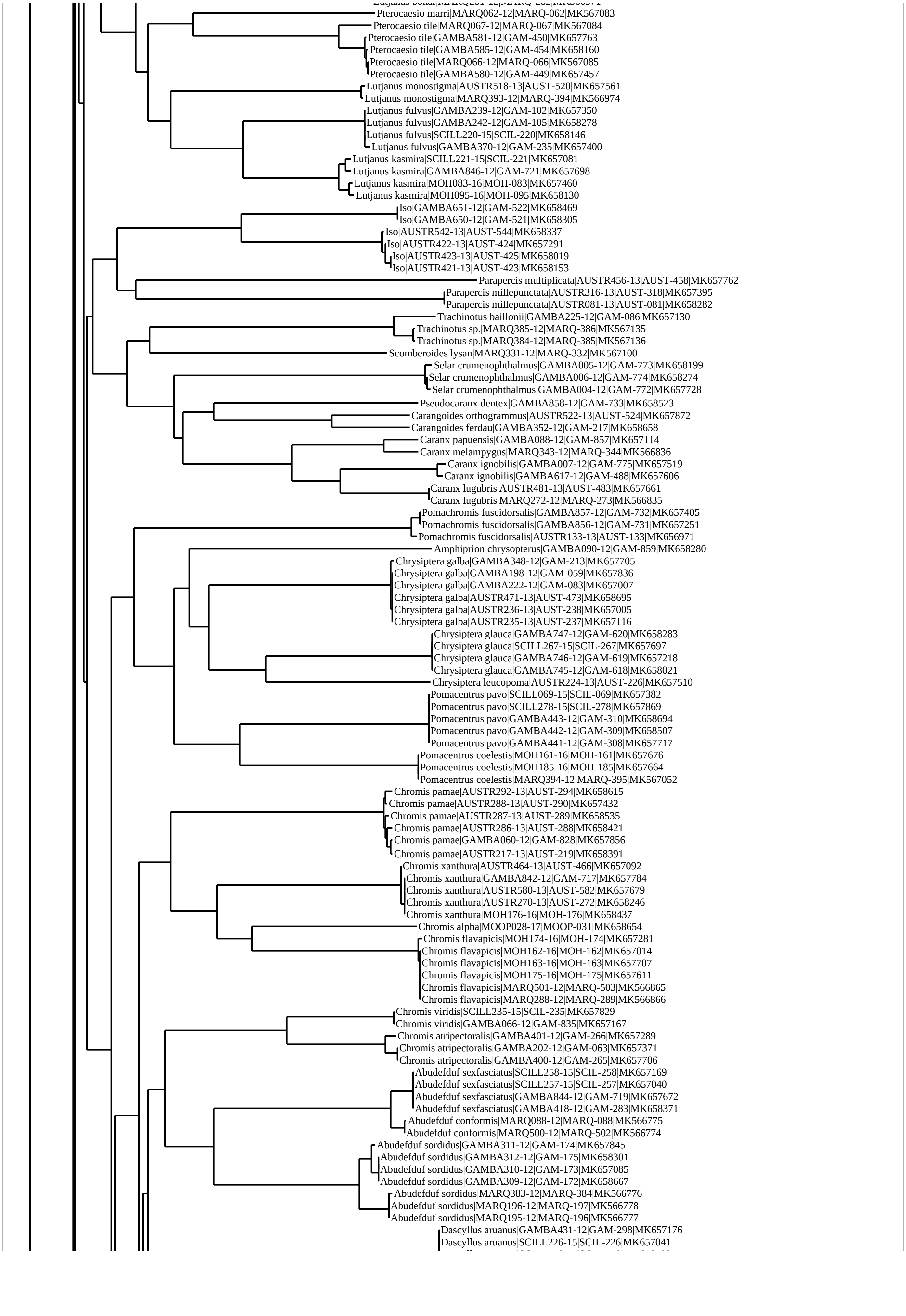
Pterocaesio marri|MARQ062-12|MARQ-062|MK567083
Pterocaesio tile|MARQ067-12|MARQ-067|MK567084
Pterocaesio tile|GAMBA581-12|GAM-450|MK657763
Pterocaesio tile|GAMBA585-12|GAM-454|MK658160
Pterocaesio tile|MARQ066-12|MARQ-066|MK567085
Pterocaesio tile|GAMBA580-12|GAM-449|MK657457
Lutjanus monostigma|AUSTR518-13|AUST-520|MK657561
Lutjanus monostigma|MARQ393-12|MARQ-394|MK566974
Lutjanus fulvus|GAMBA239-12|GAM-102|MK657350
Lutjanus fulvus|GAMBA242-12|GAM-105|MK658278
Lutjanus fulvus|SCILL220-15|SCIL-220|MK658146
Lutjanus fulvus|GAMBA370-12|GAM-235|MK657400
Lutjanus kasmira|SCILL221-15|SCIL-221|MK657081
Lutjanus kasmira|GAMBA846-12|GAM-721|MK657698
Lutjanus kasmira|MOH083-16|MOH-083|MK657460
Lutjanus kasmira|MOH095-16|MOH-095|MK658130
Iso|GAMBA651-12|GAM-522|MK658469
Iso|GAMBA650-12|GAM-521|MK658305
Iso|AUSTR542-13|AUST-544|MK658337
Iso|AUSTR422-13|AUST-424|MK657291
Iso|AUSTR423-13|AUST-425|MK658019
Iso|AUSTR421-13|AUST-423|MK658153
Parapercis multiplicata|AUSTR456-13|AUST-458|MK657762
Parapercis millepunctata|AUSTR316-13|AUST-318|MK657395
Parapercis millepunctata|AUSTR081-13|AUST-081|MK658282
Trachinotus baillonii|GAMBA225-12|GAM-086|MK657130
Trachinotus sp.|MARQ385-12|MARQ-386|MK567135
Trachinotus sp.|MARQ384-12|MARQ-385|MK567136
Scomberoides lysan|MARQ331-12|MARQ-332|MK567100
Selar crumenophthalmus|GAMBA005-12|GAM-773|MK658199
Selar crumenophthalmus|GAMBA006-12|GAM-774|MK658274
Selar crumenophthalmus|GAMBA004-12|GAM-772|MK657728
Pseudocaranx dentex|GAMBA858-12|GAM-733|MK658523
Carangoides orthogrammus|AUSTR522-13|AUST-524|MK657872
Carangoides ferdau|GAMBA352-12|GAM-217|MK658658
Caranx papuensis|GAMBA088-12|GAM-857|MK657114
Caranx melampygus|MARQ343-12|MARQ-344|MK566836
Caranx ignobilis|GAMBA007-12|GAM-775|MK657519
Caranx ignobilis|GAMBA617-12|GAM-488|MK657606
Caranx lugubris|AUSTR481-13|AUST-483|MK657661
Caranx lugubris|MARQ272-12|MARQ-273|MK566835
Pomachromis fuscidorsalis|GAMBA857-12|GAM-732|MK657405
Pomachromis fuscidorsalis|GAMBA856-12|GAM-731|MK657251
Pomachromis fuscidorsalis|AUSTR133-13|AUST-133|MK656971
Amphiprion chrysopterus|GAMBA090-12|GAM-859|MK658280
Chrysiptera galba|GAMBA348-12|GAM-213|MK657705
Chrysiptera galba|GAMBA198-12|GAM-059|MK657836
Chrysiptera galba|GAMBA222-12|GAM-083|MK657007
Chrysiptera galba|AUSTR471-13|AUST-473|MK658695
Chrysiptera galba|AUSTR236-13|AUST-238|MK657005
Chrysiptera galba|AUSTR235-13|AUST-237|MK657116
Chrysiptera glauca|GAMBA747-12|GAM-620|MK658283
Chrysiptera glauca|SCILL267-15|SCIL-267|MK657697
Chrysiptera glauca|GAMBA746-12|GAM-619|MK657218
Chrysiptera glauca|GAMBA745-12|GAM-618|MK658021
Chrysiptera leucopoma|AUSTR224-13|AUST-226|MK657510
Pomacentrus pavo|SCILL069-15|SCIL-069|MK657382
Pomacentrus pavo|SCILL278-15|SCIL-278|MK657869
Pomacentrus pavo|GAMBA443-12|GAM-310|MK658694
Pomacentrus pavo|GAMBA442-12|GAM-309|MK658507
Pomacentrus pavo|GAMBA441-12|GAM-308|MK657717
Pomacentrus coelestis|MOH161-16|MOH-161|MK657676
Pomacentrus coelestis|MOH185-16|MOH-185|MK657664
Pomacentrus coelestis|MARQ394-12|MARQ-395|MK567052
Chromis pamae|AUSTR292-13|AUST-294|MK658615
Chromis pamae|AUSTR288-13|AUST-290|MK657432
Chromis pamae|AUSTR287-13|AUST-289|MK658535
Chromis pamae|AUSTR286-13|AUST-288|MK658421
Chromis pamae|GAMBA060-12|GAM-828|MK657856
Chromis pamae|AUSTR217-13|AUST-219|MK658391
Chromis xanthura|AUSTR464-13|AUST-466|MK657092
Chromis xanthura|GAMBA842-12|GAM-717|MK657784
Chromis xanthura|AUSTR580-13|AUST-582|MK657679
Chromis xanthura|AUSTR270-13|AUST-272|MK658246
Chromis xanthura|MOH176-16|MOH-176|MK658437
Chromis alpha|MOOP028-17|MOOP-031|MK658654
Chromis flavapicis|MOH174-16|MOH-174|MK657281
Chromis flavapicis|MOH162-16|MOH-162|MK657014
Chromis flavapicis|MOH163-16|MOH-163|MK657707
Chromis flavapicis|MOH175-16|MOH-175|MK657611
Chromis flavapicis|MARQ501-12|MARQ-503|MK566865
Chromis flavapicis|MARQ288-12|MARQ-289|MK566866
Chromis viridis|SCILL235-15|SCIL-235|MK657829
Chromis viridis|GAMBA066-12|GAM-835|MK657167
Chromis atripectoralis|GAMBA401-12|GAM-266|MK657289
Chromis atripectoralis|GAMBA202-12|GAM-063|MK657371
Chromis atripectoralis|GAMBA400-12|GAM-265|MK657706
Abudefduf sexfasciatus|SCILL258-15|SCIL-258|MK657169
Abudefduf sexfasciatus|SCILL257-15|SCIL-257|MK657040
Abudefduf sexfasciatus|GAMBA844-12|GAM-719|MK657672
Abudefduf sexfasciatus|GAMBA418-12|GAM-283|MK658371
Abudefduf conformis|MARQ088-12|MARQ-088|MK566775
Abudefduf conformis|MARQ500-12|MARQ-502|MK566774
Abudefduf sordidus|GAMBA311-12|GAM-174|MK657845
Abudefduf sordidus|GAMBA312-12|GAM-175|MK658301
Abudefduf sordidus|GAMBA310-12|GAM-173|MK657085
Abudefduf sordidus|GAMBA309-12|GAM-172|MK658667
Abudefduf sordidus|MARQ383-12|MARQ-384|MK566776
Abudefduf sordidus|MARQ196-12|MARQ-197|MK566778
Abudefduf sordidus|MARQ195-12|MARQ-196|MK566777
Dascyllus aruanus|GAMBA431-12|GAM-298|MK657176
Dascyllus aruanus|SCILL226-15|SCIL-226|MK657041

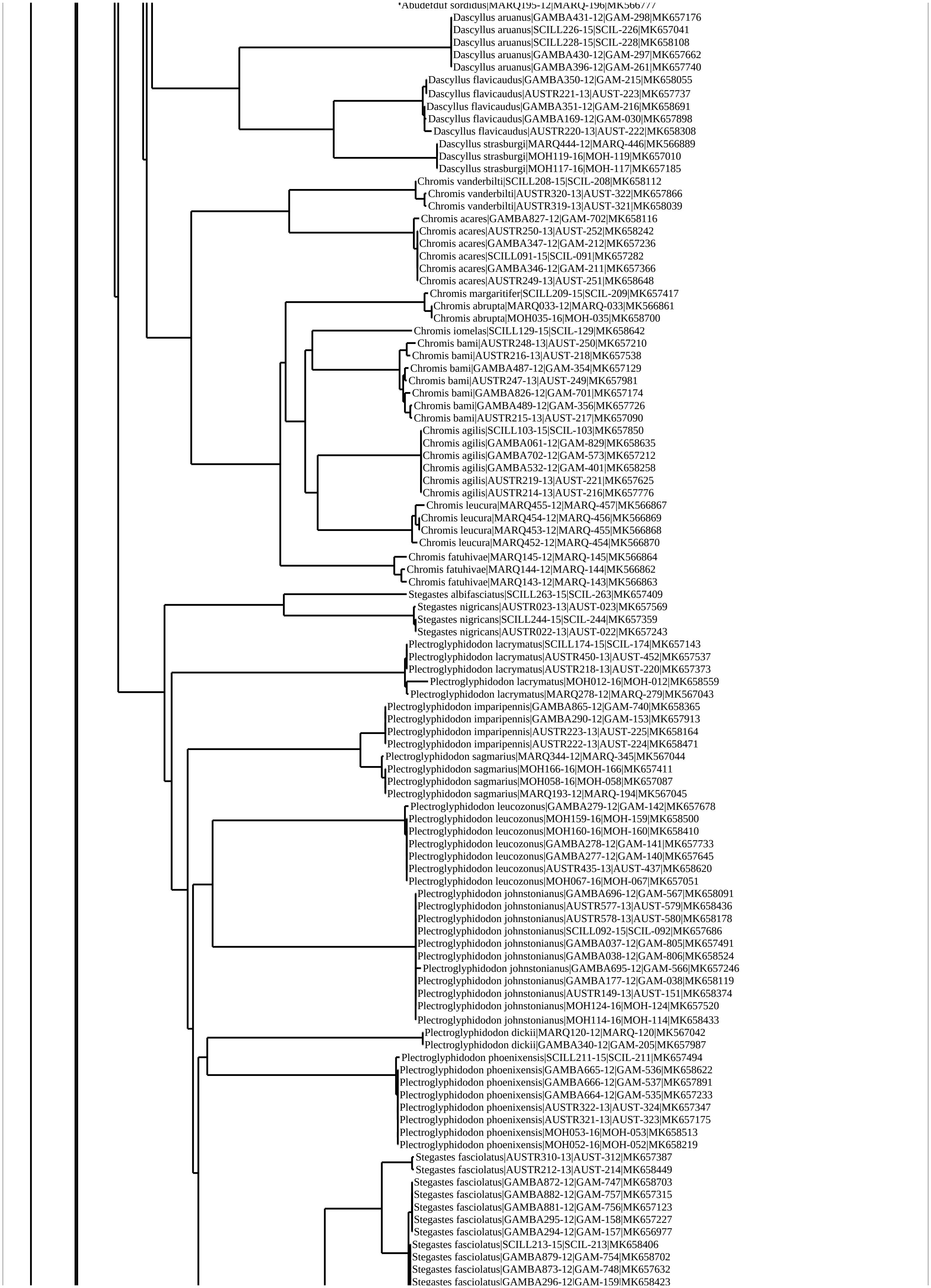
Abudefduf sordidus|MARQ195-12|MARQ-196|MK566777
Dascyllus aruanus|GAMBA431-12|GAM-298|MK657176
Dascyllus aruanus|SCILL226-15|SCIL-226|MK657041
Dascyllus aruanus|SCILL228-15|SCIL-228|MK658108
Dascyllus aruanus|GAMBA430-12|GAM-297|MK657662
Dascyllus aruanus|GAMBA396-12|GAM-261|MK657740
Dascyllus flavicaudus|GAMBA350-12|GAM-215|MK658055
Dascyllus flavicaudus|AUSTR221-13|AUST-223|MK657737
Dascyllus flavicaudus|GAMBA351-12|GAM-216|MK658691
Dascyllus flavicaudus|GAMBA169-12|GAM-030|MK657898
Dascyllus flavicaudus|AUSTR220-13|AUST-222|MK658308
Dascyllus strasburgi|MARQ444-12|MARQ-446|MK566889
Dascyllus strasburgi|MOH119-16|MOH-119|MK657010
Dascyllus strasburgi|MOH117-16|MOH-117|MK657185
Chromis vanderbilti|SCILL208-15|SCIL-208|MK658112
Chromis vanderbilti|AUSTR320-13|AUST-322|MK657866
Chromis vanderbilti|AUSTR319-13|AUST-321|MK658039
Chromis acares|GAMBA827-12|GAM-702|MK658116
Chromis acares|AUSTR250-13|AUST-252|MK658242
Chromis acares|GAMBA347-12|GAM-212|MK657236
Chromis acares|SCILL091-15|SCIL-091|MK657282
Chromis acares|GAMBA346-12|GAM-211|MK657366
Chromis acares|AUSTR249-13|AUST-251|MK658648
Chromis margaritifer|SCILL209-15|SCIL-209|MK657417
Chromis abrupta|MARQ033-12|MARQ-033|MK566861
Chromis abrupta|MOH035-16|MOH-035|MK658700
Chromis iomelas|SCILL129-15|SCIL-129|MK658642
Chromis bami|AUSTR248-13|AUST-250|MK657210
Chromis bami|AUSTR216-13|AUST-218|MK657538
Chromis bami|GAMBA487-12|GAM-354|MK657129
Chromis bami|AUSTR247-13|AUST-249|MK657981
Chromis bami|GAMBA826-12|GAM-701|MK657174
Chromis bami|GAMBA489-12|GAM-356|MK657726
Chromis bami|AUSTR215-13|AUST-217|MK657090
Chromis agilis|SCILL103-15|SCIL-103|MK657850
Chromis agilis|GAMBA061-12|GAM-829|MK658635
Chromis agilis|GAMBA702-12|GAM-573|MK657212
Chromis agilis|GAMBA532-12|GAM-401|MK658258
Chromis agilis|AUSTR219-13|AUST-221|MK657625
Chromis agilis|AUSTR214-13|AUST-216|MK657776
Chromis leucura|MARQ455-12|MARQ-457|MK566867
Chromis leucura|MARQ454-12|MARQ-456|MK566869
Chromis leucura|MARQ453-12|MARQ-455|MK566868
Chromis leucura|MARQ452-12|MARQ-454|MK566870
Chromis fatuhivae|MARQ145-12|MARQ-145|MK566864
Chromis fatuhivae|MARQ144-12|MARQ-144|MK566862
Chromis fatuhivae|MARQ143-12|MARQ-143|MK566863
Stegastes albifasciatus|SCILL263-15|SCIL-263|MK657409
Stegastes nigricans|AUSTR023-13|AUST-023|MK657569
Stegastes nigricans|SCILL244-15|SCIL-244|MK657359
Stegastes nigricans|AUSTR022-13|AUST-022|MK657243
Plectroglyphidodon lacrymatus|SCILL174-15|SCIL-174|MK657143
Plectroglyphidodon lacrymatus|AUSTR450-13|AUST-452|MK657537
Plectroglyphidodon lacrymatus|AUSTR218-13|AUST-220|MK657373
Plectroglyphidodon lacrymatus|MOH012-16|MOH-012|MK658559
Plectroglyphidodon lacrymatus|MARQ278-12|MARQ-279|MK567043
Plectroglyphidodon imparipennis|GAMBA865-12|GAM-740|MK658365
Plectroglyphidodon imparipennis|GAMBA290-12|GAM-153|MK657913
Plectroglyphidodon imparipennis|AUSTR223-13|AUST-225|MK658164
Plectroglyphidodon imparipennis|AUSTR222-13|AUST-224|MK658471
Plectroglyphidodon sagmarius|MARQ344-12|MARQ-345|MK567044
Plectroglyphidodon sagmarius|MOH166-16|MOH-166|MK657411
Plectroglyphidodon sagmarius|MOH058-16|MOH-058|MK657087
Plectroglyphidodon sagmarius|MARQ193-12|MARQ-194|MK567045
Plectroglyphidodon leucozonus|GAMBA279-12|GAM-142|MK657678
Plectroglyphidodon leucozonus|MOH159-16|MOH-159|MK658500
Plectroglyphidodon leucozonus|MOH160-16|MOH-160|MK658410
Plectroglyphidodon leucozonus|GAMBA278-12|GAM-141|MK657733
Plectroglyphidodon leucozonus|GAMBA277-12|GAM-140|MK657645
Plectroglyphidodon leucozonus|AUSTR435-13|AUST-437|MK658620
Plectroglyphidodon leucozonus|MOH067-16|MOH-067|MK657051
Plectroglyphidodon johnstonianus|GAMBA696-12|GAM-567|MK658091
Plectroglyphidodon johnstonianus|AUSTR577-13|AUST-579|MK658436
Plectroglyphidodon johnstonianus|AUSTR578-13|AUST-580|MK658178
Plectroglyphidodon johnstonianus|SCILL092-15|SCIL-092|MK657686
Plectroglyphidodon johnstonianus|GAMBA037-12|GAM-805|MK657491
Plectroglyphidodon johnstonianus|GAMBA038-12|GAM-806|MK658524
Plectroglyphidodon johnstonianus|GAMBA695-12|GAM-566|MK657246
Plectroglyphidodon johnstonianus|GAMBA177-12|GAM-038|MK658119
Plectroglyphidodon johnstonianus|AUSTR149-13|AUST-151|MK658374
Plectroglyphidodon johnstonianus|MOH124-16|MOH-124|MK657520
Plectroglyphidodon johnstonianus|MOH114-16|MOH-114|MK658433
Plectroglyphidodon dickii|MARQ120-12|MARQ-120|MK567042
Plectroglyphidodon dickii|GAMBA340-12|GAM-205|MK657987
Plectroglyphidodon phoenixensis|SCILL211-15|SCIL-211|MK657494
Plectroglyphidodon phoenixensis|GAMBA665-12|GAM-536|MK658622
Plectroglyphidodon phoenixensis|GAMBA666-12|GAM-537|MK657891
Plectroglyphidodon phoenixensis|GAMBA664-12|GAM-535|MK657233
Plectroglyphidodon phoenixensis|AUSTR322-13|AUST-324|MK657347
Plectroglyphidodon phoenixensis|AUSTR321-13|AUST-323|MK657175
Plectroglyphidodon phoenixensis|MOH053-16|MOH-053|MK658513
Plectroglyphidodon phoenixensis|MOH052-16|MOH-052|MK658219
Stegastes fasciolatus|AUSTR310-13|AUST-312|MK657387
Stegastes fasciolatus|AUSTR212-13|AUST-214|MK658449
Stegastes fasciolatus|GAMBA872-12|GAM-747|MK658703
Stegastes fasciolatus|GAMBA882-12|GAM-757|MK657315
Stegastes fasciolatus|GAMBA881-12|GAM-756|MK657123
Stegastes fasciolatus|GAMBA295-12|GAM-158|MK657227
Stegastes fasciolatus|GAMBA294-12|GAM-157|MK656977
Stegastes fasciolatus|SCILL213-15|SCIL-213|MK658406
Stegastes fasciolatus|GAMBA879-12|GAM-754|MK658702
Stegastes fasciolatus|GAMBA873-12|GAM-748|MK657632
Stegastes fasciolatus|GAMBA296-12|GAM-159|MK658423

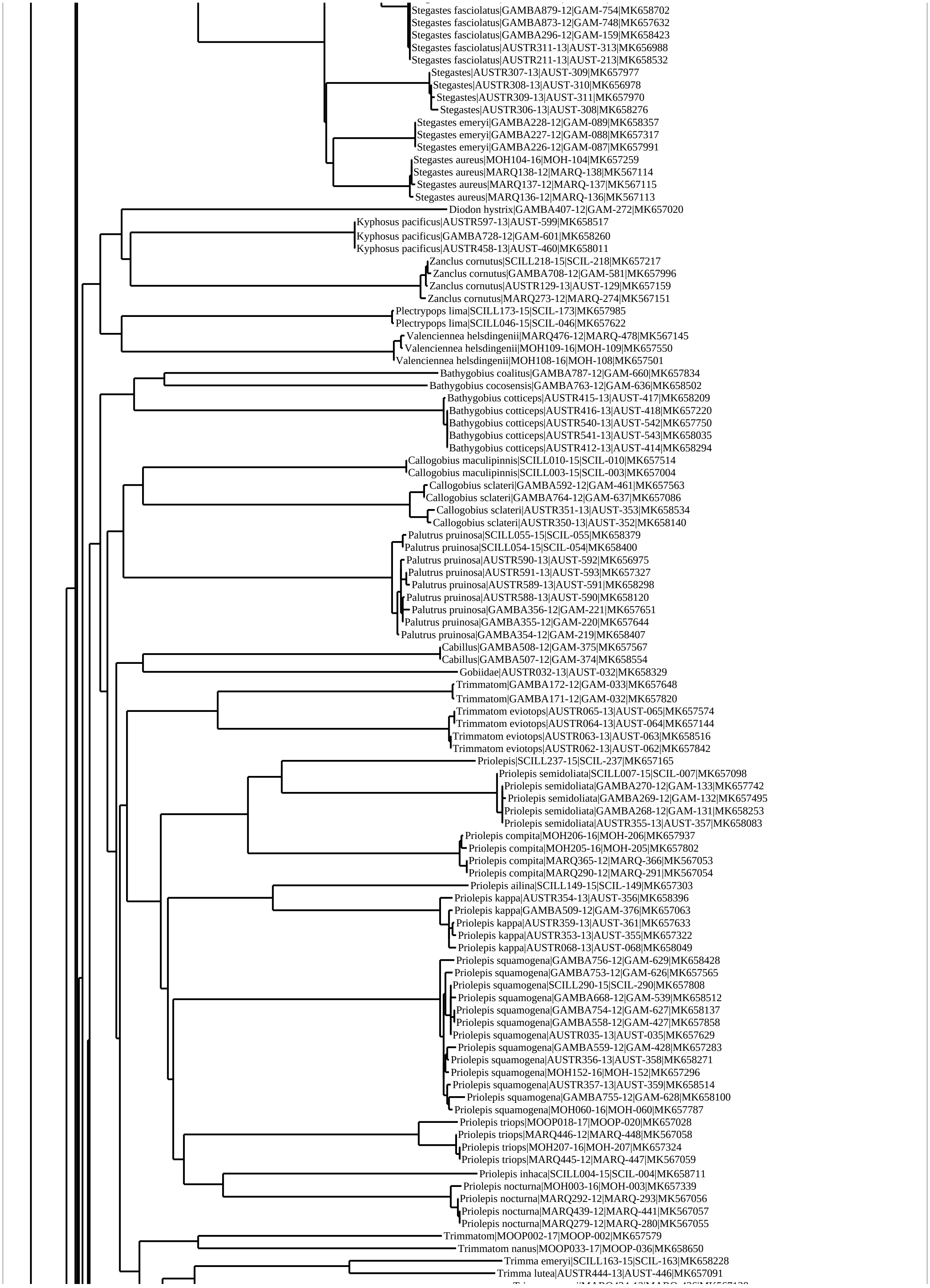
Stegastes fasciolatus|GAMBA879-12|GAM-754|MK658702
Stegastes fasciolatus|GAMBA873-12|GAM-748|MK657632
Stegastes fasciolatus|GAMBA296-12|GAM-159|MK658423
Stegastes fasciolatus|AUSTR311-13|AUST-313|MK656988
Stegastes fasciolatus|AUSTR211-13|AUST-213|MK658532
Stegastes|AUSTR307-13|AUST-309|MK657977
Stegastes|AUSTR308-13|AUST-310|MK656978
Stegastes|AUSTR309-13|AUST-311|MK657970
Stegastes|AUSTR306-13|AUST-308|MK658276
Stegastes emeryi|GAMBA228-12|GAM-089|MK658357
Stegastes emeryi|GAMBA227-12|GAM-088|MK657317
Stegastes emeryi|GAMBA226-12|GAM-087|MK657991
Stegastes aureus|MOH104-16|MOH-104|MK657259
Stegastes aureus|MARQ138-12|MARQ-138|MK567114
Stegastes aureus|MARQ137-12|MARQ-137|MK567115
Stegastes aureus|MARQ136-12|MARQ-136|MK567113
Diodon hystrix|GAMBA407-12|GAM-272|MK657020
Kyphosus pacificus|AUSTR597-13|AUST-599|MK658517
Kyphosus pacificus|GAMBA728-12|GAM-601|MK658260
Kyphosus pacificus|AUSTR458-13|AUST-460|MK658011
Zanclus cornutus|SCILL218-15|SCIL-218|MK657217
Zanclus cornutus|GAMBA708-12|GAM-581|MK657996
Zanclus cornutus|AUSTR129-13|AUST-129|MK657159
Zanclus cornutus|MARQ273-12|MARQ-274|MK567151
Plectrypops lima|SCILL173-15|SCIL-173|MK657985
Plectrypops lima|SCILL046-15|SCIL-046|MK657622
Valenciennea helsdingenii|MARQ476-12|MARQ-478|MK567145
Valenciennea helsdingenii|MOH109-16|MOH-109|MK657550
Valenciennea helsdingenii|MOH108-16|MOH-108|MK657501
Bathygobius coalitus|GAMBA787-12|GAM-660|MK657834
Bathygobius cocosensis|GAMBA763-12|GAM-636|MK658502
Bathygobius cotticeps|AUSTR415-13|AUST-417|MK658209
Bathygobius cotticeps|AUSTR416-13|AUST-418|MK657220
Bathygobius cotticeps|AUSTR540-13|AUST-542|MK657750
Bathygobius cotticeps|AUSTR541-13|AUST-543|MK658035
Bathygobius cotticeps|AUSTR412-13|AUST-414|MK658294
Callogobius maculipinnis|SCILL010-15|SCIL-010|MK657514
Callogobius maculipinnis|SCILL003-15|SCIL-003|MK657004
Callogobius sclateri|GAMBA592-12|GAM-461|MK657563
Callogobius sclateri|GAMBA764-12|GAM-637|MK657086
Callogobius sclateri|AUSTR351-13|AUST-353|MK658534
Callogobius sclateri|AUSTR350-13|AUST-352|MK658140
Palutrus pruinosa|SCILL055-15|SCIL-055|MK658379
Palutrus pruinosa|SCILL054-15|SCIL-054|MK658400
Palutrus pruinosa|AUSTR590-13|AUST-592|MK656975
Palutrus pruinosa|AUSTR591-13|AUST-593|MK657327
Palutrus pruinosa|AUSTR589-13|AUST-591|MK658298
Palutrus pruinosa|AUSTR588-13|AUST-590|MK658120
Palutrus pruinosa|GAMBA356-12|GAM-221|MK657651
Palutrus pruinosa|GAMBA355-12|GAM-220|MK657644
Palutrus pruinosa|GAMBA354-12|GAM-219|MK658407
Cabillus|GAMBA508-12|GAM-375|MK657567
Cabillus|GAMBA507-12|GAM-374|MK658554
Gobiidae|AUSTR032-13|AUST-032|MK658329
Trimmatom|GAMBA172-12|GAM-033|MK657648
Trimmatom|GAMBA171-12|GAM-032|MK657820
Trimmatom eviotops|AUSTR065-13|AUST-065|MK657574
Trimmatom eviotops|AUSTR064-13|AUST-064|MK657144
Trimmatom eviotops|AUSTR063-13|AUST-063|MK658516
Trimmatom eviotops|AUSTR062-13|AUST-062|MK657842
Priolepis|SCILL237-15|SCIL-237|MK657165
Priolepis semidoliata|SCILL007-15|SCIL-007|MK657098
Priolepis semidoliata|GAMBA270-12|GAM-133|MK657742
Priolepis semidoliata|GAMBA269-12|GAM-132|MK657495
Priolepis semidoliata|GAMBA268-12|GAM-131|MK658253
Priolepis semidoliata|AUSTR355-13|AUST-357|MK658083
Priolepis compita|MOH206-16|MOH-206|MK657937
Priolepis compita|MOH205-16|MOH-205|MK657802
Priolepis compita|MARQ365-12|MARQ-366|MK567053
Priolepis compita|MARQ290-12|MARQ-291|MK567054
Priolepis ailina|SCILL149-15|SCIL-149|MK657303
Priolepis kappa|AUSTR354-13|AUST-356|MK658396
Priolepis kappa|GAMBA509-12|GAM-376|MK657063
Priolepis kappa|AUSTR359-13|AUST-361|MK657633
Priolepis kappa|AUSTR353-13|AUST-355|MK657322
Priolepis kappa|AUSTR068-13|AUST-068|MK658049
Priolepis squamogena|GAMBA756-12|GAM-629|MK658428
Priolepis squamogena|GAMBA753-12|GAM-626|MK657565
Priolepis squamogena|SCILL290-15|SCIL-290|MK657808
Priolepis squamogena|GAMBA668-12|GAM-539|MK658512
Priolepis squamogena|GAMBA754-12|GAM-627|MK658137
Priolepis squamogena|GAMBA558-12|GAM-427|MK657858
Priolepis squamogena|AUSTR035-13|AUST-035|MK657629
Priolepis squamogena|GAMBA559-12|GAM-428|MK657283
Priolepis squamogena|AUSTR356-13|AUST-358|MK658271
Priolepis squamogena|MOH152-16|MOH-152|MK657296
Priolepis squamogena|AUSTR357-13|AUST-359|MK658514
Priolepis squamogena|GAMBA755-12|GAM-628|MK658100
Priolepis squamogena|MOH060-16|MOH-060|MK657787
Priolepis triops|MOOP018-17|MOOP-020|MK657028
Priolepis triops|MARQ446-12|MARQ-448|MK567058
Priolepis triops|MOH207-16|MOH-207|MK657324
Priolepis triops|MARQ445-12|MARQ-447|MK567059
Priolepis inhaca|SCILL004-15|SCIL-004|MK658711
Priolepis nocturna|MOH003-16|MOH-003|MK657339
Priolepis nocturna|MARQ292-12|MARQ-293|MK567056
Priolepis nocturna|MARQ439-12|MARQ-441|MK567057
Priolepis nocturna|MARQ279-12|MARQ-280|MK567055
Trimmatom|MOOP002-17|MOOP-002|MK657579
Trimmatom nanus|MOOP033-17|MOOP-036|MK658650
Trimma emeryi|SCILL163-15|SCIL-163|MK658228
Trimma lutea|AUSTR444-13|AUST-446|MK657091

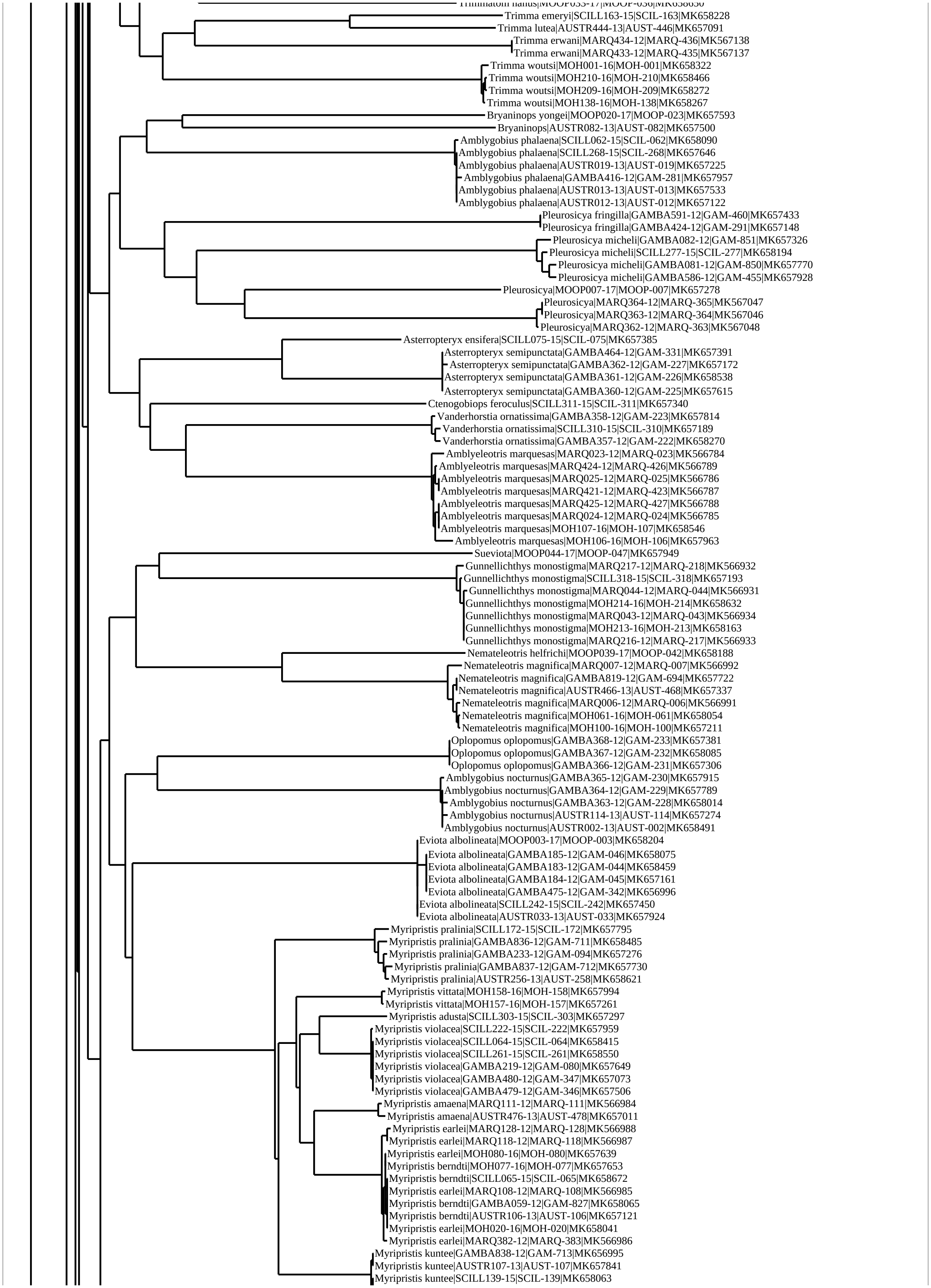
Trimma emeryi|SCILL163-15|SCIL-163|MK658228
Trimma lutea|AUSTR444-13|AUST-446|MK657091
Trimma erwani|MARQ434-12|MARQ-436|MK567138
Trimma erwani|MARQ433-12|MARQ-435|MK567137
Trimma woutsi|MOH001-16|MOH-001|MK658322
Trimma woutsi|MOH210-16|MOH-210|MK658466
Trimma woutsi|MOH209-16|MOH-209|MK658272
Trimma woutsi|MOH138-16|MOH-138|MK658267
Bryaninops yongei|MOOP020-17|MOOP-023|MK657593
Bryaninops|AUSTR082-13|AUST-082|MK657500
Amblygobius phalaena|SCILL062-15|SCIL-062|MK658090
Amblygobius phalaena|SCILL268-15|SCIL-268|MK657646
Amblygobius phalaena|AUSTR019-13|AUST-019|MK657225
Amblygobius phalaena|GAMBA416-12|GAM-281|MK657957
Amblygobius phalaena|AUSTR013-13|AUST-013|MK657533
Amblygobius phalaena|AUSTR012-13|AUST-012|MK657122
Pleurosicya fringilla|GAMBA591-12|GAM-460|MK657433
Pleurosicya fringilla|GAMBA424-12|GAM-291|MK657148
Pleurosicya micheli|GAMBA082-12|GAM-851|MK657326
Pleurosicya micheli|SCILL277-15|SCIL-277|MK658194
Pleurosicya micheli|GAMBA081-12|GAM-850|MK657770
Pleurosicya micheli|GAMBA586-12|GAM-455|MK657928
Pleurosicya|MOOP007-17|MOOP-007|MK657278
Pleurosicya|MARQ364-12|MARQ-365|MK567047
Pleurosicya|MARQ363-12|MARQ-364|MK567046
Pleurosicya|MARQ362-12|MARQ-363|MK567048
Asterropteryx ensifera|SCILL075-15|SCIL-075|MK657385
Asterropteryx semipunctata|GAMBA464-12|GAM-331|MK657391
Asterropteryx semipunctata|GAMBA362-12|GAM-227|MK657172
Asterropteryx semipunctata|GAMBA361-12|GAM-226|MK658538
Asterropteryx semipunctata|GAMBA360-12|GAM-225|MK657615
Ctenogobiops feroculus|SCILL311-15|SCIL-311|MK657340
Vanderhorstia ornatissima|GAMBA358-12|GAM-223|MK657814
Vanderhorstia ornatissima|SCILL310-15|SCIL-310|MK657189
Vanderhorstia ornatissima|GAMBA357-12|GAM-222|MK658270
Amblyeleotris marquesas|MARQ023-12|MARQ-023|MK566784
Amblyeleotris marquesas|MARQ424-12|MARQ-426|MK566789
Amblyeleotris marquesas|MARQ025-12|MARQ-025|MK566786
Amblyeleotris marquesas|MARQ421-12|MARQ-423|MK566787
Amblyeleotris marquesas|MARQ425-12|MARQ-427|MK566788
Amblyeleotris marquesas|MARQ024-12|MARQ-024|MK566785
Amblyeleotris marquesas|MOH107-16|MOH-107|MK658546
Amblyeleotris marquesas|MOH106-16|MOH-106|MK657963
Sueviota|MOOP044-17|MOOP-047|MK657949
Gunnellichthys monostigma|MARQ217-12|MARQ-218|MK566932
Gunnellichthys monostigma|SCILL318-15|SCIL-318|MK657193
Gunnellichthys monostigma|MARQ044-12|MARQ-044|MK566931
Gunnellichthys monostigma|MOH214-16|MOH-214|MK658632
Gunnellichthys monostigma|MARQ043-12|MARQ-043|MK566934
Gunnellichthys monostigma|MOH213-16|MOH-213|MK658163
Gunnellichthys monostigma|MARQ216-12|MARQ-217|MK566933
Nemateleotris helfrichi|MOOP039-17|MOOP-042|MK658188
Nemateleotris magnifica|MARQ007-12|MARQ-007|MK566992
Nemateleotris magnifica|GAMBA819-12|GAM-694|MK657722
Nemateleotris magnifica|AUSTR466-13|AUST-468|MK657337
Nemateleotris magnifica|MARQ006-12|MARQ-006|MK566991
Nemateleotris magnifica|MOH061-16|MOH-061|MK658054
Nemateleotris magnifica|MOH100-16|MOH-100|MK657211
Oplopomus oplopomus|GAMBA368-12|GAM-233|MK657381
Oplopomus oplopomus|GAMBA367-12|GAM-232|MK658085
Oplopomus oplopomus|GAMBA366-12|GAM-231|MK657306
Amblygobius nocturnus|GAMBA365-12|GAM-230|MK657915
Amblygobius nocturnus|GAMBA364-12|GAM-229|MK657789
Amblygobius nocturnus|GAMBA363-12|GAM-228|MK658014
Amblygobius nocturnus|AUSTR114-13|AUST-114|MK657274
Amblygobius nocturnus|AUSTR002-13|AUST-002|MK658491
Eviota albolineata|MOOP003-17|MOOP-003|MK658204
Eviota albolineata|GAMBA185-12|GAM-046|MK658075
Eviota albolineata|GAMBA183-12|GAM-044|MK658459
Eviota albolineata|GAMBA184-12|GAM-045|MK657161
Eviota albolineata|GAMBA475-12|GAM-342|MK656996
Eviota albolineata|SCILL242-15|SCIL-242|MK657450
Eviota albolineata|AUSTR033-13|AUST-033|MK657924
Myripristis pralinia|SCILL172-15|SCIL-172|MK657795
Myripristis pralinia|GAMBA836-12|GAM-711|MK658485
Myripristis pralinia|GAMBA233-12|GAM-094|MK657276
Myripristis pralinia|GAMBA837-12|GAM-712|MK657730
Myripristis pralinia|AUSTR256-13|AUST-258|MK658621
Myripristis vittata|MOH158-16|MOH-158|MK657994
Myripristis vittata|MOH157-16|MOH-157|MK657261
Myripristis adusta|SCILL303-15|SCIL-303|MK657297
Myripristis violacea|SCILL222-15|SCIL-222|MK657959
Myripristis violacea|SCILL064-15|SCIL-064|MK658415
Myripristis violacea|SCILL261-15|SCIL-261|MK658550
Myripristis violacea|GAMBA219-12|GAM-080|MK657649
Myripristis violacea|GAMBA480-12|GAM-347|MK657073
Myripristis violacea|GAMBA479-12|GAM-346|MK657506
Myripristis amaena|MARQ111-12|MARQ-111|MK566984
Myripristis amaena|AUSTR476-13|AUST-478|MK657011
Myripristis earlei|MARQ128-12|MARQ-128|MK566988
Myripristis earlei|MARQ118-12|MARQ-118|MK566987
Myripristis earlei|MOH080-16|MOH-080|MK657639
Myripristis berndti|MOH077-16|MOH-077|MK657653
Myripristis berndti|SCILL065-15|SCIL-065|MK658672
Myripristis earlei|MARQ108-12|MARQ-108|MK566985
Myripristis berndti|GAMBA059-12|GAM-827|MK658065
Myripristis berndti|AUSTR106-13|AUST-106|MK657121
Myripristis earlei|MOH020-16|MOH-020|MK658041
Myripristis earlei|MARQ382-12|MARQ-383|MK566986
Myripristis kuntee|GAMBA838-12|GAM-713|MK656995
Myripristis kuntee|AUSTR107-13|AUST-107|MK657841
Myripristis kuntee|SCILL139-15|SCIL-139|MK658063

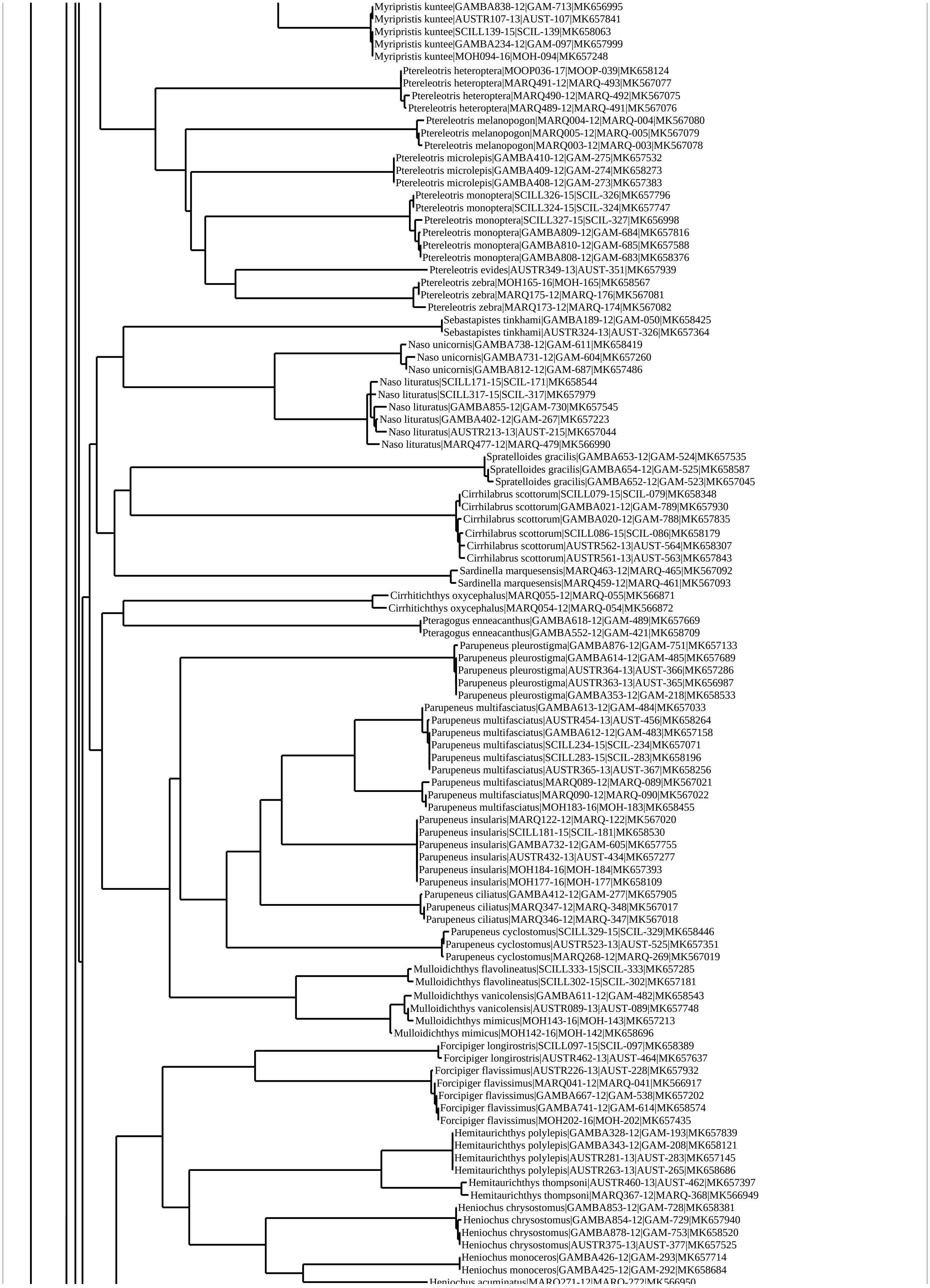
Myripristis kuntee|GAMBA838-12|GAM-713|MK656995
Myripristis kuntee|AUSTR107-13|AUST-107|MK657841
Myripristis kuntee|SCILL139-15|SCIL-139|MK658063
Myripristis kuntee|GAMBA234-12|GAM-097|MK657999
Myripristis kuntee|MOH094-16|MOH-094|MK657248
Ptereleotris heteroptera|MOOP036-17|MOOP-039|MK658124
Ptereleotris heteroptera|MARQ491-12|MARQ-493|MK567077
Ptereleotris heteroptera|MARQ490-12|MARQ-492|MK567075
Ptereleotris heteroptera|MARQ489-12|MARQ-491|MK567076
Ptereleotris melanopogon|MARQ004-12|MARQ-004|MK567080
Ptereleotris melanopogon|MARQ005-12|MARQ-005|MK567079
Ptereleotris melanopogon|MARQ003-12|MARQ-003|MK567078
Ptereleotris microlepis|GAMBA410-12|GAM-275|MK657532
Ptereleotris microlepis|GAMBA409-12|GAM-274|MK658273
Ptereleotris microlepis|GAMBA408-12|GAM-273|MK657383
Ptereleotris monoptera|SCILL326-15|SCIL-326|MK657796
Ptereleotris monoptera|SCILL324-15|SCIL-324|MK657747
Ptereleotris monoptera|SCILL327-15|SCIL-327|MK656998
Ptereleotris monoptera|GAMBA809-12|GAM-684|MK657816
Ptereleotris monoptera|GAMBA810-12|GAM-685|MK657588
Ptereleotris monoptera|GAMBA808-12|GAM-683|MK658376
Ptereleotris evides|AUSTR349-13|AUST-351|MK657939
Ptereleotris zebra|MOH165-16|MOH-165|MK658567
Ptereleotris zebra|MARQ175-12|MARQ-176|MK567081
Ptereleotris zebra|MARQ173-12|MARQ-174|MK567082
Sebastapistes tinkhami|GAMBA189-12|GAM-050|MK658425
Sebastapistes tinkhami|AUSTR324-13|AUST-326|MK657364
Naso unicornis|GAMBA738-12|GAM-611|MK658419
Naso unicornis|GAMBA731-12|GAM-604|MK657260
Naso unicornis|GAMBA812-12|GAM-687|MK657486
Naso lituratus|SCILL171-15|SCIL-171|MK658544
Naso lituratus|SCILL317-15|SCIL-317|MK657979
Naso lituratus|GAMBA855-12|GAM-730|MK657545
Naso lituratus|GAMBA402-12|GAM-267|MK657223
Naso lituratus|AUSTR213-13|AUST-215|MK657044
Naso lituratus|MARQ477-12|MARQ-479|MK566990
Spratelloides gracilis|GAMBA653-12|GAM-524|MK657535
Spratelloides gracilis|GAMBA654-12|GAM-525|MK658587
Spratelloides gracilis|GAMBA652-12|GAM-523|MK657045
Cirrhilabrus scottorum|SCILL079-15|SCIL-079|MK658348
Cirrhilabrus scottorum|GAMBA021-12|GAM-789|MK657930
Cirrhilabrus scottorum|GAMBA020-12|GAM-788|MK657835
Cirrhilabrus scottorum|SCILL086-15|SCIL-086|MK658179
Cirrhilabrus scottorum|AUSTR562-13|AUST-564|MK658307
Cirrhilabrus scottorum|AUSTR561-13|AUST-563|MK657843
Sardinella marquesensis|MARQ463-12|MARQ-465|MK567092
Sardinella marquesensis|MARQ459-12|MARQ-461|MK567093
Cirrhitichthys oxycephalus|MARQ055-12|MARQ-055|MK566871
Cirrhitichthys oxycephalus|MARQ054-12|MARQ-054|MK566872
Pteragogus enneacanthus|GAMBA618-12|GAM-489|MK657669
Pteragogus enneacanthus|GAMBA552-12|GAM-421|MK658709
Parupeneus pleurostigma|GAMBA876-12|GAM-751|MK657133
Parupeneus pleurostigma|GAMBA614-12|GAM-485|MK657689
Parupeneus pleurostigma|AUSTR364-13|AUST-366|MK657286
Parupeneus pleurostigma|AUSTR363-13|AUST-365|MK656987
Parupeneus pleurostigma|GAMBA353-12|GAM-218|MK658533
Parupeneus multifasciatus|GAMBA613-12|GAM-484|MK657033
Parupeneus multifasciatus|AUSTR454-13|AUST-456|MK658264
Parupeneus multifasciatus|GAMBA612-12|GAM-483|MK657158
Parupeneus multifasciatus|SCILL234-15|SCIL-234|MK657071
Parupeneus multifasciatus|SCILL283-15|SCIL-283|MK658196
Parupeneus multifasciatus|AUSTR365-13|AUST-367|MK658256
Parupeneus multifasciatus|MARQ089-12|MARQ-089|MK567021
Parupeneus multifasciatus|MARQ090-12|MARQ-090|MK567022
Parupeneus multifasciatus|MOH183-16|MOH-183|MK658455
Parupeneus insularis|MARQ122-12|MARQ-122|MK567020
Parupeneus insularis|SCILL181-15|SCIL-181|MK658530
Parupeneus insularis|GAMBA732-12|GAM-605|MK657755
Parupeneus insularis|AUSTR432-13|AUST-434|MK657277
Parupeneus insularis|MOH184-16|MOH-184|MK657393
Parupeneus insularis|MOH177-16|MOH-177|MK658109
Parupeneus ciliatus|GAMBA412-12|GAM-277|MK657905
Parupeneus ciliatus|MARQ347-12|MARQ-348|MK567017
Parupeneus ciliatus|MARQ346-12|MARQ-347|MK567018
Parupeneus cyclostomus|SCILL329-15|SCIL-329|MK658446
Parupeneus cyclostomus|AUSTR523-13|AUST-525|MK657351
Parupeneus cyclostomus|MARQ268-12|MARQ-269|MK567019
Mulloidichthys flavolineatus|SCILL333-15|SCIL-333|MK657285
Mulloidichthys flavolineatus|SCILL302-15|SCIL-302|MK657181
Mulloidichthys vanicolensis|GAMBA611-12|GAM-482|MK658543
Mulloidichthys vanicolensis|AUSTR089-13|AUST-089|MK657748
Mulloidichthys mimicus|MOH143-16|MOH-143|MK657213
Mulloidichthys mimicus|MOH142-16|MOH-142|MK658696
Forcipiger longirostris|SCILL097-15|SCIL-097|MK658389
Forcipiger longirostris|AUSTR462-13|AUST-464|MK657637
Forcipiger flavissimus|AUSTR226-13|AUST-228|MK657932
Forcipiger flavissimus|MARQ041-12|MARQ-041|MK566917
Forcipiger flavissimus|GAMBA667-12|GAM-538|MK657202
Forcipiger flavissimus|GAMBA741-12|GAM-614|MK658574
Forcipiger flavissimus|MOH202-16|MOH-202|MK657435
Hemitaurichthys polylepis|GAMBA328-12|GAM-193|MK657839
Hemitaurichthys polylepis|GAMBA343-12|GAM-208|MK658121
Hemitaurichthys polylepis|AUSTR281-13|AUST-283|MK657145
Hemitaurichthys polylepis|AUSTR263-13|AUST-265|MK658686
Hemitaurichthys thompsoni|AUSTR460-13|AUST-462|MK657397
Hemitaurichthys thompsoni|MARQ367-12|MARQ-368|MK566949
Heniochus chrysostomus|GAMBA853-12|GAM-728|MK658381
Heniochus chrysostomus|GAMBA854-12|GAM-729|MK657940
Heniochus chrysostomus|GAMBA878-12|GAM-753|MK658520
Heniochus chrysostomus|AUSTR375-13|AUST-377|MK657525
Heniochus monoceros|GAMBA426-12|GAM-293|MK657714
Heniochus monoceros|GAMBA425-12|GAM-292|MK658684
Heniochus acuminatus|MARQ271-12|MARQ-272|MK566950

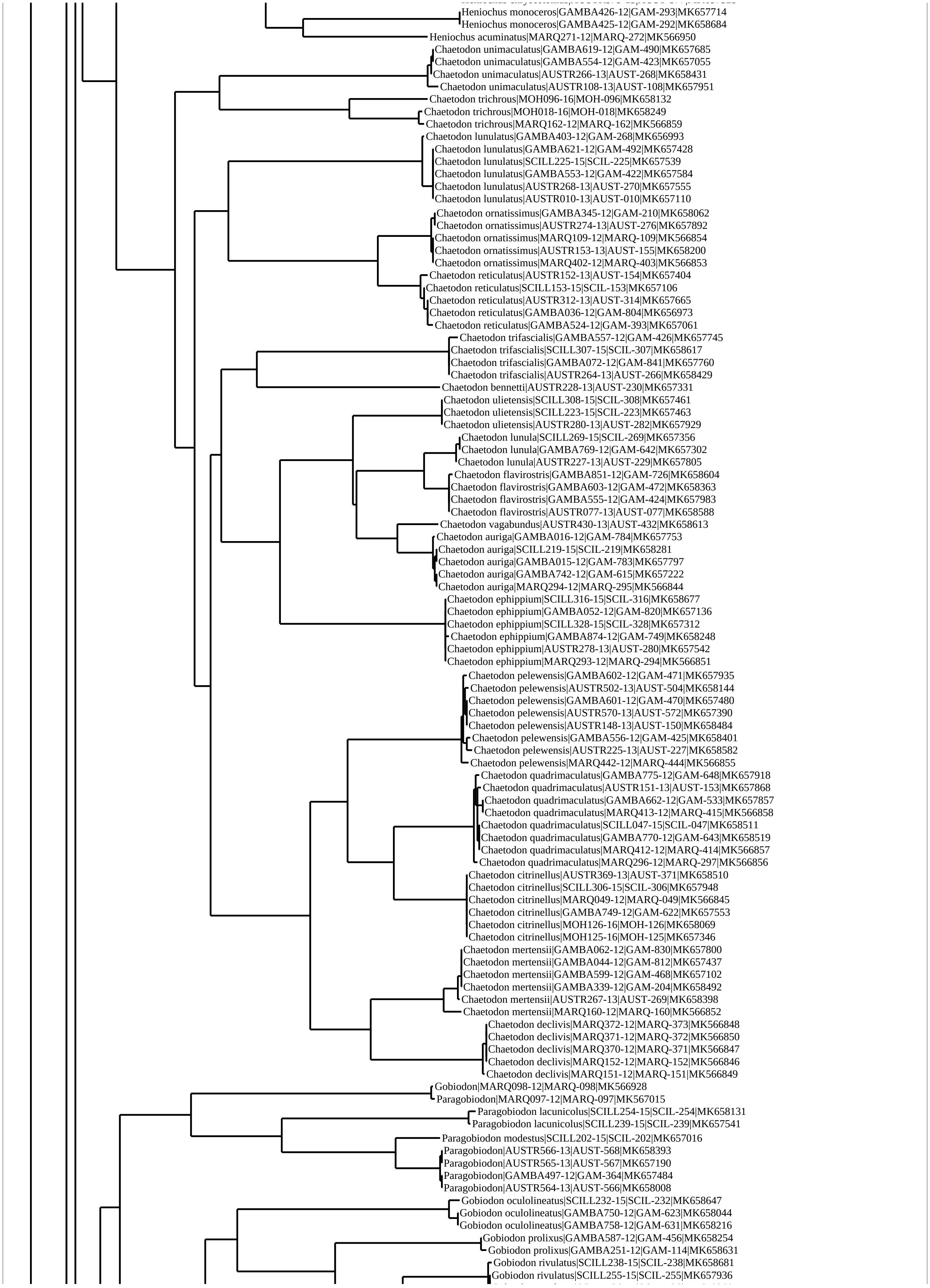
Heniochus monoceros|GAMBA426-12|GAM-293|MK657714
Heniochus monoceros|GAMBA425-12|GAM-292|MK658684
Heniochus acuminatus|MARQ271-12|MARQ-272|MK566950
Chaetodon unimaculatus|GAMBA619-12|GAM-490|MK657685
Chaetodon unimaculatus|GAMBA554-12|GAM-423|MK657055
Chaetodon unimaculatus|AUSTR266-13|AUST-268|MK658431
Chaetodon unimaculatus|AUSTR108-13|AUST-108|MK657951
Chaetodon trichrous|MOH096-16|MOH-096|MK658132
Chaetodon trichrous|MOH018-16|MOH-018|MK658249
Chaetodon trichrous|MARQ162-12|MARQ-162|MK566859
Chaetodon lunulatus|GAMBA403-12|GAM-268|MK656993
Chaetodon lunulatus|GAMBA621-12|GAM-492|MK657428
Chaetodon lunulatus|SCILL225-15|SCIL-225|MK657539
Chaetodon lunulatus|GAMBA553-12|GAM-422|MK657584
Chaetodon lunulatus|AUSTR268-13|AUST-270|MK657555
Chaetodon lunulatus|AUSTR010-13|AUST-010|MK657110
Chaetodon ornatissimus|GAMBA345-12|GAM-210|MK658062
Chaetodon ornatissimus|AUSTR274-13|AUST-276|MK657892
Chaetodon ornatissimus|MARQ109-12|MARQ-109|MK566854
Chaetodon ornatissimus|AUSTR153-13|AUST-155|MK658200
Chaetodon ornatissimus|MARQ402-12|MARQ-403|MK566853
Chaetodon reticulatus|AUSTR152-13|AUST-154|MK657404
Chaetodon reticulatus|SCILL153-15|SCIL-153|MK657106
Chaetodon reticulatus|AUSTR312-13|AUST-314|MK657665
Chaetodon reticulatus|GAMBA036-12|GAM-804|MK656973
Chaetodon reticulatus|GAMBA524-12|GAM-393|MK657061
Chaetodon trifascialis|GAMBA557-12|GAM-426|MK657745
Chaetodon trifascialis|SCILL307-15|SCIL-307|MK658617
Chaetodon trifascialis|GAMBA072-12|GAM-841|MK657760
Chaetodon trifascialis|AUSTR264-13|AUST-266|MK658429
Chaetodon bennetti|AUSTR228-13|AUST-230|MK657331
Chaetodon ulietensis|SCILL308-15|SCIL-308|MK657461
Chaetodon ulietensis|SCILL223-15|SCIL-223|MK657463
Chaetodon ulietensis|AUSTR280-13|AUST-282|MK657929
Chaetodon lunula|SCILL269-15|SCIL-269|MK657356
Chaetodon lunula|GAMBA769-12|GAM-642|MK657302
Chaetodon lunula|AUSTR227-13|AUST-229|MK657805
Chaetodon flavirostris|GAMBA851-12|GAM-726|MK658604
Chaetodon flavirostris|GAMBA603-12|GAM-472|MK658363
Chaetodon flavirostris|GAMBA555-12|GAM-424|MK657983
Chaetodon flavirostris|AUSTR077-13|AUST-077|MK658588
Chaetodon vagabundus|AUSTR430-13|AUST-432|MK658613
Chaetodon auriga|GAMBA016-12|GAM-784|MK657753
Chaetodon auriga|SCILL219-15|SCIL-219|MK658281
Chaetodon auriga|GAMBA015-12|GAM-783|MK657797
Chaetodon auriga|GAMBA742-12|GAM-615|MK657222
Chaetodon auriga|MARQ294-12|MARQ-295|MK566844
Chaetodon ephippium|SCILL316-15|SCIL-316|MK658677
Chaetodon ephippium|GAMBA052-12|GAM-820|MK657136
Chaetodon ephippium|SCILL328-15|SCIL-328|MK657312
Chaetodon ephippium|GAMBA874-12|GAM-749|MK658248
Chaetodon ephippium|AUSTR278-13|AUST-280|MK657542
Chaetodon ephippium|MARQ293-12|MARQ-294|MK566851
Chaetodon pelewensis|GAMBA602-12|GAM-471|MK657935
Chaetodon pelewensis|AUSTR502-13|AUST-504|MK658144
Chaetodon pelewensis|GAMBA601-12|GAM-470|MK657480
Chaetodon pelewensis|AUSTR570-13|AUST-572|MK657390
Chaetodon pelewensis|AUSTR148-13|AUST-150|MK658484
Chaetodon pelewensis|GAMBA556-12|GAM-425|MK658401
Chaetodon pelewensis|AUSTR225-13|AUST-227|MK658582
Chaetodon pelewensis|MARQ442-12|MARQ-444|MK566855
Chaetodon quadrimaculatus|GAMBA775-12|GAM-648|MK657918
Chaetodon quadrimaculatus|AUSTR151-13|AUST-153|MK657868
Chaetodon quadrimaculatus|GAMBA662-12|GAM-533|MK657857
Chaetodon quadrimaculatus|MARQ413-12|MARQ-415|MK566858
Chaetodon quadrimaculatus|SCILL047-15|SCIL-047|MK658511
Chaetodon quadrimaculatus|GAMBA770-12|GAM-643|MK658519
Chaetodon quadrimaculatus|MARQ412-12|MARQ-414|MK566857
Chaetodon quadrimaculatus|MARQ296-12|MARQ-297|MK566856
Chaetodon citrinellus|AUSTR369-13|AUST-371|MK658510
Chaetodon citrinellus|SCILL306-15|SCIL-306|MK657948
Chaetodon citrinellus|MARQ049-12|MARQ-049|MK566845
Chaetodon citrinellus|GAMBA749-12|GAM-622|MK657553
Chaetodon citrinellus|MOH126-16|MOH-126|MK658069
Chaetodon citrinellus|MOH125-16|MOH-125|MK657346
Chaetodon mertensii|GAMBA062-12|GAM-830|MK657800
Chaetodon mertensii|GAMBA044-12|GAM-812|MK657437
Chaetodon mertensii|GAMBA599-12|GAM-468|MK657102
Chaetodon mertensii|GAMBA339-12|GAM-204|MK658492
Chaetodon mertensii|AUSTR267-13|AUST-269|MK658398
Chaetodon mertensii|MARQ160-12|MARQ-160|MK566852
Chaetodon declivis|MARQ372-12|MARQ-373|MK566848
Chaetodon declivis|MARQ371-12|MARQ-372|MK566850
Chaetodon declivis|MARQ370-12|MARQ-371|MK566847
Chaetodon declivis|MARQ152-12|MARQ-152|MK566846
Chaetodon declivis|MARQ151-12|MARQ-151|MK566849
Gobiodon|MARQ098-12|MARQ-098|MK566928
Paragobiodon|MARQ097-12|MARQ-097|MK567015
Paragobiodon lacunicolus|SCILL254-15|SCIL-254|MK658131
Paragobiodon lacunicolus|SCILL239-15|SCIL-239|MK657541
Paragobiodon modestus|SCILL202-15|SCIL-202|MK657016
Paragobiodon|AUSTR566-13|AUST-568|MK658393
Paragobiodon|AUSTR565-13|AUST-567|MK657190
Paragobiodon|GAMBA497-12|GAM-364|MK657484
Paragobiodon|AUSTR564-13|AUST-566|MK658008
Gobiodon oculolineatus|SCILL232-15|SCIL-232|MK658647
Gobiodon oculolineatus|GAMBA750-12|GAM-623|MK658044
Gobiodon oculolineatus|GAMBA758-12|GAM-631|MK658216
Gobiodon prolixus|GAMBA587-12|GAM-456|MK658254
Gobiodon prolixus|GAMBA251-12|GAM-114|MK658631
Gobiodon rivulatus|SCILL238-15|SCIL-238|MK658681
Gobiodon rivulatus|SCILL255-15|SCIL-255|MK657936

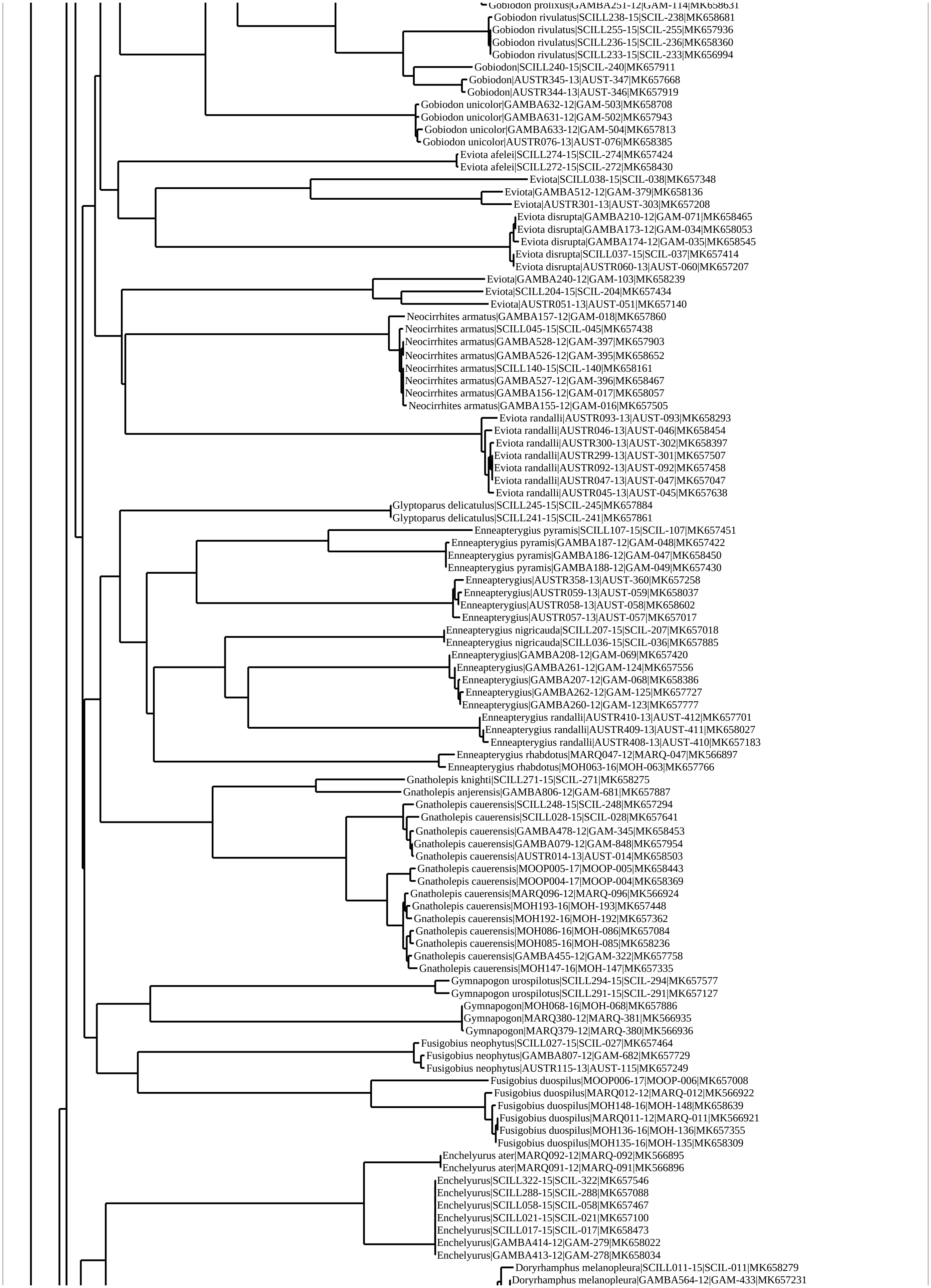
Gobiodon prolixus|GAMBA251-12|GAM-114|MK658631
Gobiodon rivulatus|SCILL238-15|SCIL-238|MK658681
Gobiodon rivulatus|SCILL255-15|SCIL-255|MK657936
Gobiodon rivulatus|SCILL236-15|SCIL-236|MK658360
Gobiodon rivulatus|SCILL233-15|SCIL-233|MK656994
Gobiodon|SCILL240-15|SCIL-240|MK657911
Gobiodon|AUSTR345-13|AUST-347|MK657668
Gobiodon|AUSTR344-13|AUST-346|MK657919
Gobiodon unicolor|GAMBA632-12|GAM-503|MK658708
Gobiodon unicolor|GAMBA631-12|GAM-502|MK657943
Gobiodon unicolor|GAMBA633-12|GAM-504|MK657813
Gobiodon unicolor|AUSTR076-13|AUST-076|MK658385
Eviota afelei|SCILL274-15|SCIL-274|MK657424
Eviota afelei|SCILL272-15|SCIL-272|MK658430
Eviota|SCILL038-15|SCIL-038|MK657348
Eviota|GAMBA512-12|GAM-379|MK658136
Eviota|AUSTR301-13|AUST-303|MK657208
Eviota disrupta|GAMBA210-12|GAM-071|MK658465
Eviota disrupta|GAMBA173-12|GAM-034|MK658053
Eviota disrupta|GAMBA174-12|GAM-035|MK658545
Eviota disrupta|SCILL037-15|SCIL-037|MK657414
Eviota disrupta|AUSTR060-13|AUST-060|MK657207
Eviota|GAMBA240-12|GAM-103|MK658239
Eviota|SCILL204-15|SCIL-204|MK657434
Eviota|AUSTR051-13|AUST-051|MK657140
Neocirrhites armatus|GAMBA157-12|GAM-018|MK657860
Neocirrhites armatus|SCILL045-15|SCIL-045|MK657438
Neocirrhites armatus|GAMBA528-12|GAM-397|MK657903
Neocirrhites armatus|GAMBA526-12|GAM-395|MK658652
Neocirrhites armatus|SCILL140-15|SCIL-140|MK658161
Neocirrhites armatus|GAMBA527-12|GAM-396|MK658467
Neocirrhites armatus|GAMBA156-12|GAM-017|MK658057
Neocirrhites armatus|GAMBA155-12|GAM-016|MK657505
Eviota randalli|AUSTR093-13|AUST-093|MK658293
Eviota randalli|AUSTR046-13|AUST-046|MK658454
Eviota randalli|AUSTR300-13|AUST-302|MK658397
Eviota randalli|AUSTR299-13|AUST-301|MK657507
Eviota randalli|AUSTR092-13|AUST-092|MK657458
Eviota randalli|AUSTR047-13|AUST-047|MK657047
Eviota randalli|AUSTR045-13|AUST-045|MK657638
Glyptoparus delicatulus|SCILL245-15|SCIL-245|MK657884
Glyptoparus delicatulus|SCILL241-15|SCIL-241|MK657861
Enneapterygius pyramis|SCILL107-15|SCIL-107|MK657451
Enneapterygius pyramis|GAMBA187-12|GAM-048|MK657422
Enneapterygius pyramis|GAMBA186-12|GAM-047|MK658450
Enneapterygius pyramis|GAMBA188-12|GAM-049|MK657430
Enneapterygius|AUSTR358-13|AUST-360|MK657258
Enneapterygius|AUSTR059-13|AUST-059|MK658037
Enneapterygius|AUSTR058-13|AUST-058|MK658602
Enneapterygius|AUSTR057-13|AUST-057|MK657017
Enneapterygius nigricauda|SCILL207-15|SCIL-207|MK657018
Enneapterygius nigricauda|SCILL036-15|SCIL-036|MK657885
Enneapterygius|GAMBA208-12|GAM-069|MK657420
Enneapterygius|GAMBA261-12|GAM-124|MK657556
Enneapterygius|GAMBA207-12|GAM-068|MK658386
Enneapterygius|GAMBA262-12|GAM-125|MK657727
Enneapterygius|GAMBA260-12|GAM-123|MK657777
Enneapterygius randalli|AUSTR410-13|AUST-412|MK657701
Enneapterygius randalli|AUSTR409-13|AUST-411|MK658027
Enneapterygius randalli|AUSTR408-13|AUST-410|MK657183
Enneapterygius rhabdotus|MARQ047-12|MARQ-047|MK566897
Enneapterygius rhabdotus|MOH063-16|MOH-063|MK657766
Gnatholepis knighti|SCILL271-15|SCIL-271|MK658275
Gnatholepis anjerensis|GAMBA806-12|GAM-681|MK657887
Gnatholepis cauerensis|SCILL248-15|SCIL-248|MK657294
Gnatholepis cauerensis|SCILL028-15|SCIL-028|MK657641
Gnatholepis cauerensis|GAMBA478-12|GAM-345|MK658453
Gnatholepis cauerensis|GAMBA079-12|GAM-848|MK657954
Gnatholepis cauerensis|AUSTR014-13|AUST-014|MK658503
Gnatholepis cauerensis|MOOP005-17|MOOP-005|MK658443
Gnatholepis cauerensis|MOOP004-17|MOOP-004|MK658369
Gnatholepis cauerensis|MARQ096-12|MARQ-096|MK566924
Gnatholepis cauerensis|MOH193-16|MOH-193|MK657448
Gnatholepis cauerensis|MOH192-16|MOH-192|MK657362
Gnatholepis cauerensis|MOH086-16|MOH-086|MK657084
Gnatholepis cauerensis|MOH085-16|MOH-085|MK658236
Gnatholepis cauerensis|GAMBA455-12|GAM-322|MK657758
Gnatholepis cauerensis|MOH147-16|MOH-147|MK657335
Gymnapogon urospilotus|SCILL294-15|SCIL-294|MK657577
Gymnapogon urospilotus|SCILL291-15|SCIL-291|MK657127
Gymnapogon|MOH068-16|MOH-068|MK657886
Gymnapogon|MARQ380-12|MARQ-381|MK566935
Gymnapogon|MARQ379-12|MARQ-380|MK566936
Fusigobius neophytus|SCILL027-15|SCIL-027|MK657464
Fusigobius neophytus|GAMBA807-12|GAM-682|MK657729
Fusigobius neophytus|AUSTR115-13|AUST-115|MK657249
Fusigobius duospilus|MOOP006-17|MOOP-006|MK657008
Fusigobius duospilus|MARQ012-12|MARQ-012|MK566922
Fusigobius duospilus|MOH148-16|MOH-148|MK658639
Fusigobius duospilus|MARQ011-12|MARQ-011|MK566921
Fusigobius duospilus|MOH136-16|MOH-136|MK657355
Fusigobius duospilus|MOH135-16|MOH-135|MK658309
Enchelyurus ater|MARQ092-12|MARQ-092|MK566895
Enchelyurus ater|MARQ091-12|MARQ-091|MK566896
Enchelyurus|SCILL322-15|SCIL-322|MK657546
Enchelyurus|SCILL288-15|SCIL-288|MK657088
Enchelyurus|SCILL058-15|SCIL-058|MK657467
Enchelyurus|SCILL021-15|SCIL-021|MK657100
Enchelyurus|SCILL017-15|SCIL-017|MK658473
Enchelyurus|GAMBA414-12|GAM-279|MK658022
Enchelyurus|GAMBA413-12|GAM-278|MK658034
Doryrhamphus melanopleura|SCILL011-15|SCIL-011|MK658279
Doryrhamphus melanopleura|GAMBA564-12|GAM-433|MK657231

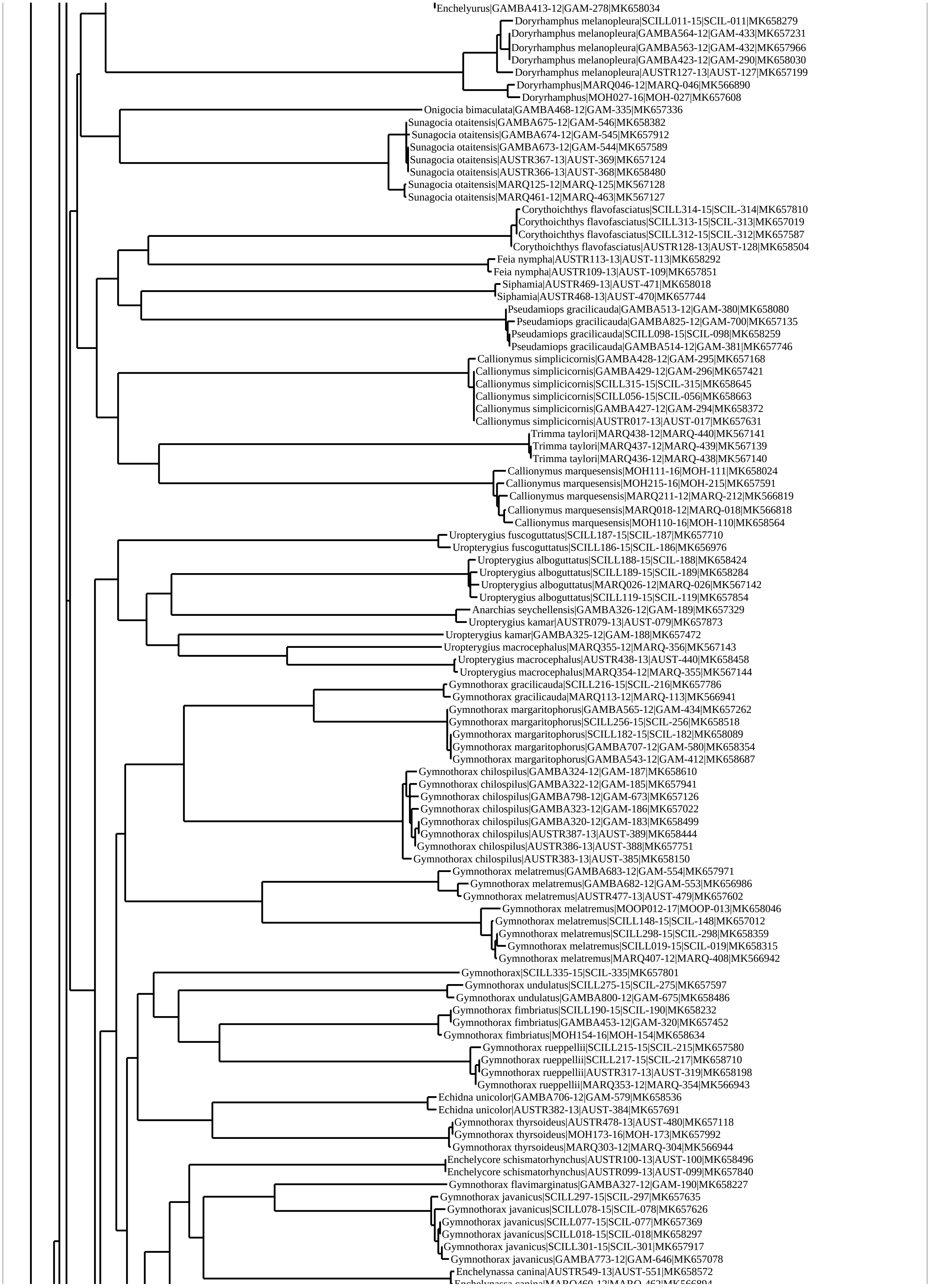
Enchelyurus|GAMBA413-12|GAM-278|MK658034
Doryrhamphus melanopleura|SCILL011-15|SCIL-011|MK658279
Doryrhamphus melanopleura|GAMBA564-12|GAM-433|MK657231
Doryrhamphus melanopleura|GAMBA563-12|GAM-432|MK657966
Doryrhamphus melanopleura|GAMBA423-12|GAM-290|MK658030
Doryrhamphus melanopleura|AUSTR127-13|AUST-127|MK657199
Doryrhamphus|MARQ046-12|MARQ-046|MK566890
Doryrhamphus|MOH027-16|MOH-027|MK657608
Onigocia bimaculata|GAMBA468-12|GAM-335|MK657336
Sunagocia otaitensis|GAMBA675-12|GAM-546|MK658382
Sunagocia otaitensis|GAMBA674-12|GAM-545|MK657912
Sunagocia otaitensis|GAMBA673-12|GAM-544|MK657589
Sunagocia otaitensis|AUSTR367-13|AUST-369|MK657124
Sunagocia otaitensis|AUSTR366-13|AUST-368|MK658480
Sunagocia otaitensis|MARQ125-12|MARQ-125|MK567128
Sunagocia otaitensis|MARQ461-12|MARQ-463|MK567127
Corythoichthys flavofasciatus|SCILL314-15|SCIL-314|MK657810
Corythoichthys flavofasciatus|SCILL313-15|SCIL-313|MK657019
Corythoichthys flavofasciatus|SCILL312-15|SCIL-312|MK657587
Corythoichthys flavofasciatus|AUSTR128-13|AUST-128|MK658504
Feia nympha|AUSTR113-13|AUST-113|MK658292
Feia nympha|AUSTR109-13|AUST-109|MK657851
Siphamia|AUSTR469-13|AUST-471|MK658018
Siphamia|AUSTR468-13|AUST-470|MK657744
Pseudamiops gracilicauda|GAMBA513-12|GAM-380|MK658080
Pseudamiops gracilicauda|GAMBA825-12|GAM-700|MK657135
Pseudamiops gracilicauda|SCILL098-15|SCIL-098|MK658259
Pseudamiops gracilicauda|GAMBA514-12|GAM-381|MK657746
Callionymus simplicicornis|GAMBA428-12|GAM-295|MK657168
Callionymus simplicicornis|GAMBA429-12|GAM-296|MK657421
Callionymus simplicicornis|SCILL315-15|SCIL-315|MK658645
Callionymus simplicicornis|SCILL056-15|SCIL-056|MK658663
Callionymus simplicicornis|GAMBA427-12|GAM-294|MK658372
Callionymus simplicicornis|AUSTR017-13|AUST-017|MK657631
Trimma taylori|MARQ438-12|MARQ-440|MK567141
Trimma taylori|MARQ437-12|MARQ-439|MK567139
Trimma taylori|MARQ436-12|MARQ-438|MK567140
Callionymus marquesensis|MOH111-16|MOH-111|MK658024
Callionymus marquesensis|MOH215-16|MOH-215|MK657591
Callionymus marquesensis|MARQ211-12|MARQ-212|MK566819
Callionymus marquesensis|MARQ018-12|MARQ-018|MK566818
Callionymus marquesensis|MOH110-16|MOH-110|MK658564
Uropterygius fuscoguttatus|SCILL187-15|SCIL-187|MK657710
Uropterygius fuscoguttatus|SCILL186-15|SCIL-186|MK656976
Uropterygius alboguttatus|SCILL188-15|SCIL-188|MK658424
Uropterygius alboguttatus|SCILL189-15|SCIL-189|MK658284
Uropterygius alboguttatus|MARQ026-12|MARQ-026|MK567142
Uropterygius alboguttatus|SCILL119-15|SCIL-119|MK657854
Anarchias seychellensis|GAMBA326-12|GAM-189|MK657329
Uropterygius kamar|AUSTR079-13|AUST-079|MK657873
Uropterygius kamar|GAMBA325-12|GAM-188|MK657472
Uropterygius macrocephalus|MARQ355-12|MARQ-356|MK567143
Uropterygius macrocephalus|AUSTR438-13|AUST-440|MK658458
Uropterygius macrocephalus|MARQ354-12|MARQ-355|MK567144
Gymnothorax gracilicauda|SCILL216-15|SCIL-216|MK657786
Gymnothorax gracilicauda|MARQ113-12|MARQ-113|MK566941
Gymnothorax margaritophorus|GAMBA565-12|GAM-434|MK657262
Gymnothorax margaritophorus|SCILL256-15|SCIL-256|MK658518
Gymnothorax margaritophorus|SCILL182-15|SCIL-182|MK658089
Gymnothorax margaritophorus|GAMBA707-12|GAM-580|MK658354
Gymnothorax margaritophorus|GAMBA543-12|GAM-412|MK658687
Gymnothorax chilospilus|GAMBA324-12|GAM-187|MK658610
Gymnothorax chilospilus|GAMBA322-12|GAM-185|MK657941
Gymnothorax chilospilus|GAMBA798-12|GAM-673|MK657126
Gymnothorax chilospilus|GAMBA323-12|GAM-186|MK657022
Gymnothorax chilospilus|GAMBA320-12|GAM-183|MK658499
Gymnothorax chilospilus|AUSTR387-13|AUST-389|MK658444
Gymnothorax chilospilus|AUSTR386-13|AUST-388|MK657751
Gymnothorax chilospilus|AUSTR383-13|AUST-385|MK658150
Gymnothorax melatremus|GAMBA683-12|GAM-554|MK657971
Gymnothorax melatremus|GAMBA682-12|GAM-553|MK656986
Gymnothorax melatremus|AUSTR477-13|AUST-479|MK657602
Gymnothorax melatremus|MOOP012-17|MOOP-013|MK658046
Gymnothorax melatremus|SCILL148-15|SCIL-148|MK657012
Gymnothorax melatremus|SCILL298-15|SCIL-298|MK658359
Gymnothorax melatremus|SCILL019-15|SCIL-019|MK658315
Gymnothorax melatremus|MARQ407-12|MARQ-408|MK566942
Gymnothorax|SCILL335-15|SCIL-335|MK657801
Gymnothorax undulatus|SCILL275-15|SCIL-275|MK657597
Gymnothorax undulatus|GAMBA800-12|GAM-675|MK658486
Gymnothorax fimbriatus|SCILL190-15|SCIL-190|MK658232
Gymnothorax fimbriatus|GAMBA453-12|GAM-320|MK657452
Gymnothorax fimbriatus|MOH154-16|MOH-154|MK658634
Gymnothorax rueppellii|SCILL215-15|SCIL-215|MK657580
Gymnothorax rueppellii|SCILL217-15|SCIL-217|MK658710
Gymnothorax rueppellii|AUSTR317-13|AUST-319|MK658198
Gymnothorax rueppellii|MARQ353-12|MARQ-354|MK566943
Echidna unicolor|GAMBA706-12|GAM-579|MK658536
Echidna unicolor|AUSTR382-13|AUST-384|MK657691
Gymnothorax thyrsoideus|AUSTR478-13|AUST-480|MK657118
Gymnothorax thyrsoideus|MOH173-16|MOH-173|MK657992
Gymnothorax thyrsoideus|MARQ303-12|MARQ-304|MK566944
Enchelycore schismatorhynchus|AUSTR100-13|AUST-100|MK658496
Enchelycore schismatorhynchus|AUSTR099-13|AUST-099|MK657840
Gymnothorax flavimarginatus|GAMBA327-12|GAM-190|MK658227
Gymnothorax javanicus|SCILL297-15|SCIL-297|MK657635
Gymnothorax javanicus|SCILL078-15|SCIL-078|MK657626
Gymnothorax javanicus|SCILL077-15|SCIL-077|MK657369
Gymnothorax javanicus|SCILL018-15|SCIL-018|MK658297
Gymnothorax javanicus|SCILL301-15|SCIL-301|MK657917
Gymnothorax javanicus|GAMBA773-12|GAM-646|MK657078
Enchelynassa canina|AUSTR549-13|AUST-551|MK658572

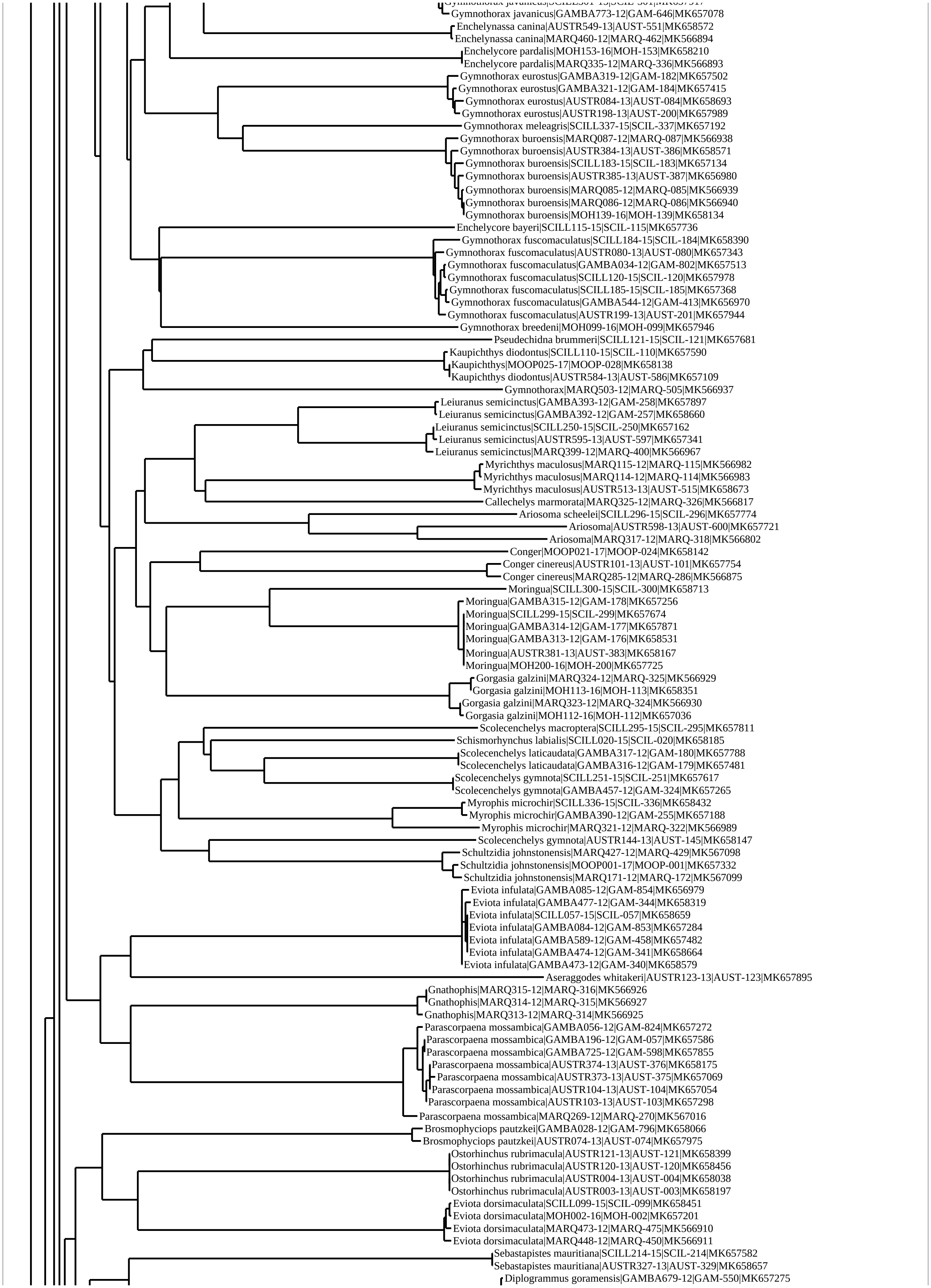
Gymnothorax javanicus|GAMBA773-12|GAM-646|MK657078
Enchelynassa canina|AUSTR549-13|AUST-551|MK658572
Enchelynassa canina|MARQ460-12|MARQ-462|MK566894
Enchelycore pardalis|MOH153-16|MOH-153|MK658210
Enchelycore pardalis|MARQ335-12|MARQ-336|MK566893
Gymnothorax eurostus|GAMBA319-12|GAM-182|MK657502
Gymnothorax eurostus|GAMBA321-12|GAM-184|MK657415
Gymnothorax eurostus|AUSTR084-13|AUST-084|MK658693
Gymnothorax eurostus|AUSTR198-13|AUST-200|MK657989
Gymnothorax meleagris|SCILL337-15|SCIL-337|MK657192
Gymnothorax buroensis|MARQ087-12|MARQ-087|MK566938
Gymnothorax buroensis|AUSTR384-13|AUST-386|MK658571
Gymnothorax buroensis|SCILL183-15|SCIL-183|MK657134
Gymnothorax buroensis|AUSTR385-13|AUST-387|MK656980
Gymnothorax buroensis|MARQ085-12|MARQ-085|MK566939
Gymnothorax buroensis|MARQ086-12|MARQ-086|MK566940
Gymnothorax buroensis|MOH139-16|MOH-139|MK658134
Enchelycore bayeri|SCILL115-15|SCIL-115|MK657736
Gymnothorax fuscomaculatus|SCILL184-15|SCIL-184|MK658390
Gymnothorax fuscomaculatus|AUSTR080-13|AUST-080|MK657343
Gymnothorax fuscomaculatus|GAMBA034-12|GAM-802|MK657513
Gymnothorax fuscomaculatus|SCILL120-15|SCIL-120|MK657978
Gymnothorax fuscomaculatus|SCILL185-15|SCIL-185|MK657368
Gymnothorax fuscomaculatus|GAMBA544-12|GAM-413|MK656970
Gymnothorax fuscomaculatus|AUSTR199-13|AUST-201|MK657944
Gymnothorax breedeni|MOH099-16|MOH-099|MK657946
Pseudechidna brummeri|SCILL121-15|SCIL-121|MK657681
Kaupichthys diodontus|SCILL110-15|SCIL-110|MK657590
Kaupichthys|MOOP025-17|MOOP-028|MK658138
Kaupichthys diodontus|AUSTR584-13|AUST-586|MK657109
Gymnothorax|MARQ503-12|MARQ-505|MK566937
Leiuranus semicinctus|GAMBA393-12|GAM-258|MK657897
Leiuranus semicinctus|GAMBA392-12|GAM-257|MK658660
Leiuranus semicinctus|SCILL250-15|SCIL-250|MK657162
Leiuranus semicinctus|AUSTR595-13|AUST-597|MK657341
Leiuranus semicinctus|MARQ399-12|MARQ-400|MK566967
Myrichthys maculosus|MARQ115-12|MARQ-115|MK566982
Myrichthys maculosus|MARQ114-12|MARQ-114|MK566983
Myrichthys maculosus|AUSTR513-13|AUST-515|MK658673
Callechelys marmorata|MARQ325-12|MARQ-326|MK566817
Ariosoma scheelei|SCILL296-15|SCIL-296|MK657774
Ariosoma|AUSTR598-13|AUST-600|MK657721
Ariosoma|MARQ317-12|MARQ-318|MK566802
Conger|MOOP021-17|MOOP-024|MK658142
Conger cinereus|AUSTR101-13|AUST-101|MK657754
Conger cinereus|MARQ285-12|MARQ-286|MK566875
Moringua|SCILL300-15|SCIL-300|MK658713
Moringua|GAMBA315-12|GAM-178|MK657256
Moringua|SCILL299-15|SCIL-299|MK657674
Moringua|GAMBA314-12|GAM-177|MK657871
Moringua|GAMBA313-12|GAM-176|MK658531
Moringua|AUSTR381-13|AUST-383|MK658167
Moringua|MOH200-16|MOH-200|MK657725
Gorgasia galzini|MARQ324-12|MARQ-325|MK566929
Gorgasia galzini|MOH113-16|MOH-113|MK658351
Gorgasia galzini|MARQ323-12|MARQ-324|MK566930
Gorgasia galzini|MOH112-16|MOH-112|MK657036
Scolecenchelys macroptera|SCILL295-15|SCIL-295|MK657811
Schismorhynchus labialis|SCILL020-15|SCIL-020|MK658185
Scolecenchelys laticaudata|GAMBA317-12|GAM-180|MK657788
Scolecenchelys laticaudata|GAMBA316-12|GAM-179|MK657481
Scolecenchelys gymnota|SCILL251-15|SCIL-251|MK657617
Scolecenchelys gymnota|GAMBA457-12|GAM-324|MK657265
Myrophis microchir|SCILL336-15|SCIL-336|MK658432
Myrophis microchir|GAMBA390-12|GAM-255|MK657188
Myrophis microchir|MARQ321-12|MARQ-322|MK566989
Scolecenchelys gymnota|AUSTR144-13|AUST-145|MK658147
Schultzidia johnstonensis|MARQ427-12|MARQ-429|MK567098
Schultzidia johnstonensis|MOOP001-17|MOOP-001|MK657332
Schultzidia johnstonensis|MARQ171-12|MARQ-172|MK567099
Eviota infulata|GAMBA085-12|GAM-854|MK656979
Eviota infulata|GAMBA477-12|GAM-344|MK658319
Eviota infulata|SCILL057-15|SCIL-057|MK658659
Eviota infulata|GAMBA084-12|GAM-853|MK657284
Eviota infulata|GAMBA589-12|GAM-458|MK657482
Eviota infulata|GAMBA474-12|GAM-341|MK658664
Eviota infulata|GAMBA473-12|GAM-340|MK658579
Aseraggodes whitakeri|AUSTR123-13|AUST-123|MK657895
Gnathophis|MARQ315-12|MARQ-316|MK566926
Gnathophis|MARQ314-12|MARQ-315|MK566927
Gnathophis|MARQ313-12|MARQ-314|MK566925
Parascorpaena mossambica|GAMBA056-12|GAM-824|MK657272
Parascorpaena mossambica|GAMBA196-12|GAM-057|MK657586
Parascorpaena mossambica|GAMBA725-12|GAM-598|MK657855
Parascorpaena mossambica|AUSTR374-13|AUST-376|MK658175
Parascorpaena mossambica|AUSTR373-13|AUST-375|MK657069
Parascorpaena mossambica|AUSTR104-13|AUST-104|MK657054
Parascorpaena mossambica|AUSTR103-13|AUST-103|MK657298
Parascorpaena mossambica|MARQ269-12|MARQ-270|MK567016
Brosmophyciops pautzkei|GAMBA028-12|GAM-796|MK658066
Brosmophyciops pautzkei|AUSTR074-13|AUST-074|MK657975
Ostorhinchus rubrimacula|AUSTR121-13|AUST-121|MK658399
Ostorhinchus rubrimacula|AUSTR120-13|AUST-120|MK658456
Ostorhinchus rubrimacula|AUSTR004-13|AUST-004|MK658038
Ostorhinchus rubrimacula|AUSTR003-13|AUST-003|MK658197
Eviota dorsimaculata|SCILL099-15|SCIL-099|MK658451
Eviota dorsimaculata|MOH002-16|MOH-002|MK657201
Eviota dorsimaculata|MARQ473-12|MARQ-475|MK566910
Eviota dorsimaculata|MARQ448-12|MARQ-450|MK566911
Sebastapistes mauritiana|SCILL214-15|SCIL-214|MK657582
Sebastapistes mauritiana|AUSTR327-13|AUST-329|MK658657
Diplogrammus goramensis|GAMBA679-12|GAM-550|MK657275

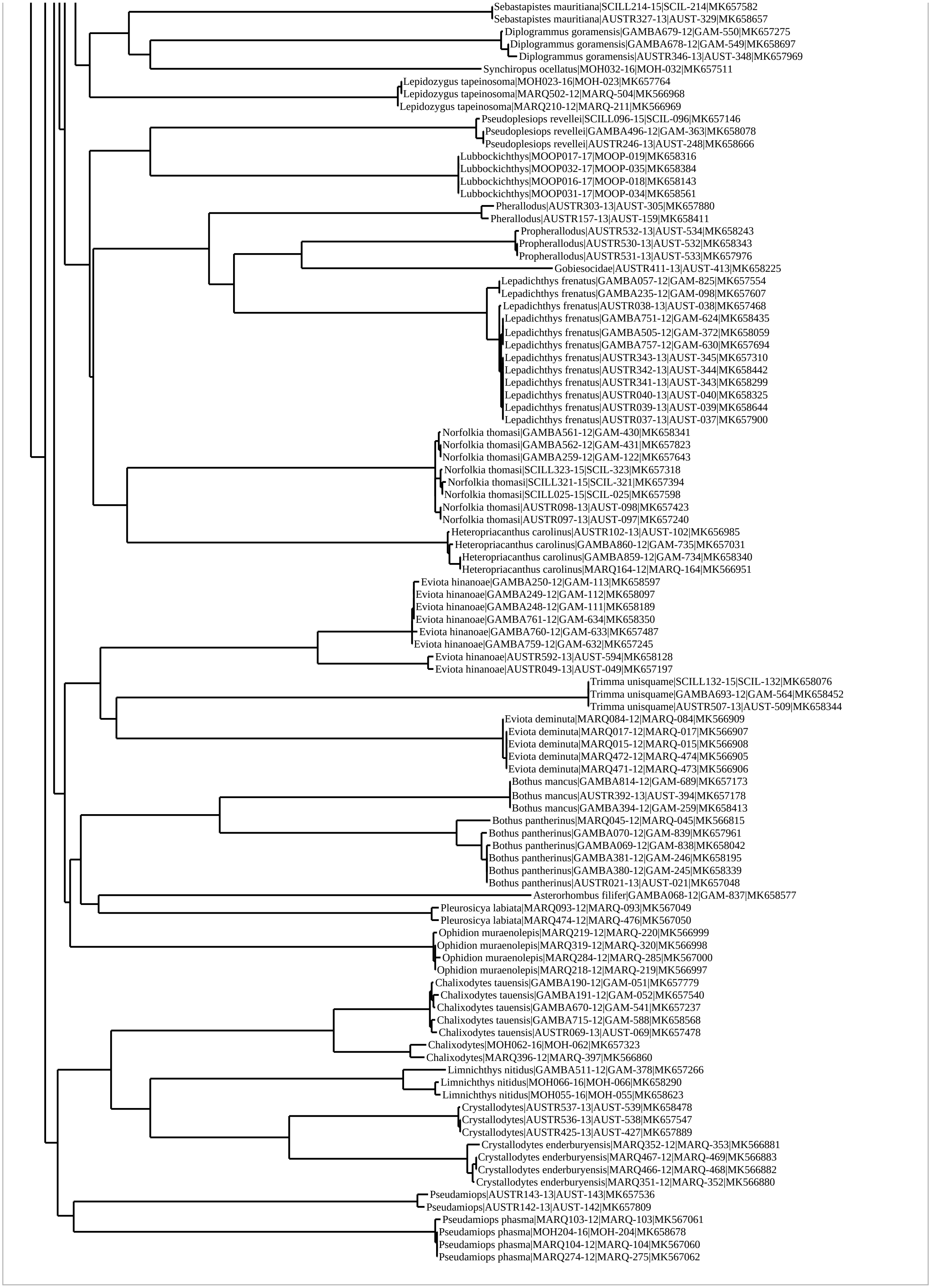
Sebastapistes mauritiana|SCILL214-15|SCIL-214|MK657582
Sebastapistes mauritiana|AUSTR327-13|AUST-329|MK658657
Diplogrammus goramensis|GAMBA679-12|GAM-550|MK657275
Diplogrammus goramensis|GAMBA678-12|GAM-549|MK658697
Diplogrammus goramensis|AUSTR346-13|AUST-348|MK657969
Synchiropus ocellatus|MOH032-16|MOH-032|MK657511
Lepidozygus tapeinosoma|MOH023-16|MOH-023|MK657764
Lepidozygus tapeinosoma|MARQ502-12|MARQ-504|MK566968
Lepidozygus tapeinosoma|MARQ210-12|MARQ-211|MK566969
Pseudoplesiops revellei|SCILL096-15|SCIL-096|MK657146
Pseudoplesiops revellei|GAMBA496-12|GAM-363|MK658078
Pseudoplesiops revellei|AUSTR246-13|AUST-248|MK658666
Lubbockichthys|MOOP017-17|MOOP-019|MK658316
Lubbockichthys|MOOP032-17|MOOP-035|MK658384
Lubbockichthys|MOOP016-17|MOOP-018|MK658143
Lubbockichthys|MOOP031-17|MOOP-034|MK658561
Pherallodus|AUSTR303-13|AUST-305|MK657880
Pherallodus|AUSTR157-13|AUST-159|MK658411
Propherallodus|AUSTR532-13|AUST-534|MK658243
Propherallodus|AUSTR530-13|AUST-532|MK658343
Propherallodus|AUSTR531-13|AUST-533|MK657976
Gobiesocidae|AUSTR411-13|AUST-413|MK658225
Lepadichthys frenatus|GAMBA057-12|GAM-825|MK657554
Lepadichthys frenatus|GAMBA235-12|GAM-098|MK657607
Lepadichthys frenatus|AUSTR038-13|AUST-038|MK657468
Lepadichthys frenatus|GAMBA751-12|GAM-624|MK658435
Lepadichthys frenatus|GAMBA505-12|GAM-372|MK658059
Lepadichthys frenatus|GAMBA757-12|GAM-630|MK657694
Lepadichthys frenatus|AUSTR343-13|AUST-345|MK657310
Lepadichthys frenatus|AUSTR342-13|AUST-344|MK658442
Lepadichthys frenatus|AUSTR341-13|AUST-343|MK658299
Lepadichthys frenatus|AUSTR040-13|AUST-040|MK658325
Lepadichthys frenatus|AUSTR039-13|AUST-039|MK658644
Lepadichthys frenatus|AUSTR037-13|AUST-037|MK657900
Norfolkia thomasi|GAMBA561-12|GAM-430|MK658341
Norfolkia thomasi|GAMBA562-12|GAM-431|MK657823
Norfolkia thomasi|GAMBA259-12|GAM-122|MK657643
Norfolkia thomasi|SCILL323-15|SCIL-323|MK657318
Norfolkia thomasi|SCILL321-15|SCIL-321|MK657394
Norfolkia thomasi|SCILL025-15|SCIL-025|MK657598
Norfolkia thomasi|AUSTR098-13|AUST-098|MK657423
Norfolkia thomasi|AUSTR097-13|AUST-097|MK657240
Heteropriacanthus carolinus|AUSTR102-13|AUST-102|MK656985
Heteropriacanthus carolinus|GAMBA860-12|GAM-735|MK657031
Heteropriacanthus carolinus|GAMBA859-12|GAM-734|MK658340
Heteropriacanthus carolinus|MARQ164-12|MARQ-164|MK566951
Eviota hinanoae|GAMBA250-12|GAM-113|MK658597
Eviota hinanoae|GAMBA249-12|GAM-112|MK658097
Eviota hinanoae|GAMBA248-12|GAM-111|MK658189
Eviota hinanoae|GAMBA761-12|GAM-634|MK658350
Eviota hinanoae|GAMBA760-12|GAM-633|MK657487
Eviota hinanoae|GAMBA759-12|GAM-632|MK657245
Eviota hinanoae|AUSTR592-13|AUST-594|MK658128
Eviota hinanoae|AUSTR049-13|AUST-049|MK657197
Trimma unisquame|SCILL132-15|SCIL-132|MK658076
Trimma unisquame|GAMBA693-12|GAM-564|MK658452
Trimma unisquame|AUSTR507-13|AUST-509|MK658344
Eviota deminuta|MARQ084-12|MARQ-084|MK566909
Eviota deminuta|MARQ017-12|MARQ-017|MK566907
Eviota deminuta|MARQ015-12|MARQ-015|MK566908
Eviota deminuta|MARQ472-12|MARQ-474|MK566905
Eviota deminuta|MARQ471-12|MARQ-473|MK566906
Bothus mancus|GAMBA814-12|GAM-689|MK657173
Bothus mancus|AUSTR392-13|AUST-394|MK657178
Bothus mancus|GAMBA394-12|GAM-259|MK658413
Bothus pantherinus|MARQ045-12|MARQ-045|MK566815
Bothus pantherinus|GAMBA070-12|GAM-839|MK657961
Bothus pantherinus|GAMBA069-12|GAM-838|MK658042
Bothus pantherinus|GAMBA381-12|GAM-246|MK658195
Bothus pantherinus|GAMBA380-12|GAM-245|MK658339
Bothus pantherinus|AUSTR021-13|AUST-021|MK657048
Asterorhombus filifer|GAMBA068-12|GAM-837|MK658577
Pleurosicya labiata|MARQ093-12|MARQ-093|MK567049
Pleurosicya labiata|MARQ474-12|MARQ-476|MK567050
Ophidion muraenolepis|MARQ219-12|MARQ-220|MK566999
Ophidion muraenolepis|MARQ319-12|MARQ-320|MK566998
Ophidion muraenolepis|MARQ284-12|MARQ-285|MK567000
Ophidion muraenolepis|MARQ218-12|MARQ-219|MK566997
Chalixodytes tauensis|GAMBA190-12|GAM-051|MK657779
Chalixodytes tauensis|GAMBA191-12|GAM-052|MK657540
Chalixodytes tauensis|GAMBA670-12|GAM-541|MK657237
Chalixodytes tauensis|GAMBA715-12|GAM-588|MK658568
Chalixodytes tauensis|AUSTR069-13|AUST-069|MK657478
Chalixodytes|MOH062-16|MOH-062|MK657323
Chalixodytes|MARQ396-12|MARQ-397|MK566860
Limnichthys nitidus|GAMBA511-12|GAM-378|MK657266
Limnichthys nitidus|MOH066-16|MOH-066|MK658290
Limnichthys nitidus|MOH055-16|MOH-055|MK658623
Crystallodytes|AUSTR537-13|AUST-539|MK658478
Crystallodytes|AUSTR536-13|AUST-538|MK657547
Crystallodytes|AUSTR425-13|AUST-427|MK657889
Crystallodytes enderburyensis|MARQ352-12|MARQ-353|MK566881
Crystallodytes enderburyensis|MARQ467-12|MARQ-469|MK566883
Crystallodytes enderburyensis|MARQ466-12|MARQ-468|MK566882
Crystallodytes enderburyensis|MARQ351-12|MARQ-352|MK566880
Pseudamiops|AUSTR143-13|AUST-143|MK657536
Pseudamiops|AUSTR142-13|AUST-142|MK657809
Pseudamiops phasma|MARQ103-12|MARQ-103|MK567061
Pseudamiops phasma|MOH204-16|MOH-204|MK658678
Pseudamiops phasma|MARQ104-12|MARQ-104|MK567060
Pseudamiops phasma|MARQ274-12|MARQ-275|MK567062
